# Post-Translational Modifications Remodel Proteome-Wide Ligandability

**DOI:** 10.1101/2025.07.31.667978

**Authors:** Weichao Li, Qijia Wei, Paolo Governa, Manuel Llanos, Tzu-Yuan Chiu, Jacob M. Wozniak, Appaso M. Jadhav, Clara Gathmann, Matthew Holcomb, Jacob Cravatt, Ashok Dongre, Mia L. Huang, Stefano Forli, Christopher G. Parker

## Abstract

Post-translational modifications (PTMs) vastly expand the diversity of human proteome, dynamically reshaping protein activity, interactions, and localization in response to environmental, pharmacologic, and disease-associated cues. While it is well established that PTMs modulate protein function, structure, and biomolecular interactions, their proteome-wide impact on small-molecule recognition—and thus druggability—remains largely unexplored. Here, we introduce a chemical proteomic strategy to delineate how PTM states remodel protein ligandability in human cells. By deploying broad profiling photoaffinity probes, we identified over 400 functionally diverse proteins whose ability to engage small molecules is impacted by phosphorylation or N-linked glycosylation status. Integration of binding site mapping with structural analyses revealed a diverse array of PTM-dependent pockets. Among these targets, we discovered that the phosphorylation status of common oncogenic KRAS mutants impact the action of small molecules, including clinically approved inhibitors. These findings illuminate an underappreciated, PTM-governed layer of proteome plasticity and uncover opportunities for the development of chemical probes to selectively target proteins in defined modification states.

## Introduction

Advances in genomic and transcriptomic technologies have transformed our ability to chart how biological systems respond to physiological, pathological, and pharmacological perturbations. Yet, these methods capture only a portion of the complexity of cellular regulation, as a significant fraction of human biology is governed at the proteome level through post-translational modifications (PTMs), which dynamically modulate protein structure, function, and interactions in ways not captured by genetic approaches. Over 680 PTMs have been identified in humans, ranging from the addition of simple functional groups (e.g., methylation and acetylation) to large complex biomolecules (e.g., complex glycans and small proteins)^1^. This diversity can generate hundreds of thousands of distinct proteoforms per cell, summing to millions across all human cell types ^2^. In addition to their structural variety, PTMs are often highly dynamic, being added, removed, or exchanged in response to diverse stimuli and environmental cues. When these regulatory processes go awry, they frequently drive pathological outcomes, underscoring the critical need to further understand how PTMs affect protein function and how this dysregulation can be therapeutically targeted ^3^.

Towards this end, extensive efforts have been directed toward global profiling of PTM dynamics using mass spectrometry (MS)-based proteomics ^4^. Because many PTMs are substoichiometric and transient, enrichment techniques such as immunoprecipitation, lectin-based capture, bioorthogonal labeling, and metabolic incorporation are commonly employed to isolate modified proteins for MS identification and quantification ^5–7^. While these collective approaches have significantly advanced our knowledge of PTM prevalence and dynamics, they offer limited insight into how PTMs alter protein structure, activity, or pharmacological tractability. To begin bridging this gap, powerful chemical proteomic methods, including activity-based protein profiling (ABPP), have been deployed to globally assess enzyme activity, cysteine reactivity status, and covalent liganding across various biological states ^8–13^. However, these approaches depend on electrophilic probes targeting nucleophilic side chains, which limit their scope to enzymes or sites with reactive residues. Moreover, the use of ABPP to profile proteomic states has largely been restricted to lysates^14,15^, where protein interactions and compartmentalization are disrupted—potentially obscuring PTM-regulated binding events or generating artifacts due to probing of non-native conditions. To better understand how PTMs or other factors alter protein state, new sensitive strategies are needed that are agnostic to protein functional class or composition and operate in intact systems.

Among the most prevalent and functionally consequential PTMs are phosphorylation and N-linked glycosylation. Phosphorylation orchestrates signaling pathways by tuning protein activity, localization, and interactions, whereas N-linked glycosylation can regulate protein folding, stability, trafficking, and cell-cell recognition^16,17^. Dysregulation of these PTMs is a hallmark of diverse diseases, including cancers, autoimmune disorders, and metabolic syndromes. Correspondingly, the machinery responsible for installing, removing, and recognizing PTMs (e.g., kinases/phosphatases, glycosyltransferases/glycosidases, and acetyltransferases/deacetylates) are often drug targets ^18–20^. Yet pharmacological targeting of PTM-modifying enzymes often leads to pleiotropic effects, typically affecting the manifold substrates of these enzymes rather than only disease-driving pathways. An alternative strategy is to directly assess how PTMs alter the ligandability of proteins—i.e., their ability to engage small molecules—across biological contexts. Such an approach could reveal disease-relevant proteoforms with differential druggability, enabling the design of more selective therapeutic agents.

Here, we present a chemoproteomic strategy that leverages broadly interactive, fragment-based photoaffinity probes to map reversible small-molecule–protein interactions as a function of phosphorylation and glycosylation status directly in cells (Fig. 1a). Applying this approach, we uncovered hundreds of proteins—spanning diverse functional classes—that display PTM-dependent interactions with drug-like small molecules. Proteome-wide binding site mapping revealed both direct and indirect mechanisms by which PTMs reshape small-molecule accessibility. Notably, we found that phosphorylation status of KRAS reconfigures ligand engagement, altering binding of clinically approved inhibitors to numerous oncogenic mutants. Our findings unveil previously underappreciated layers of regulation in protein-small molecule interactions and highlight critical implications for drug targeting.

**Fig. 1.**
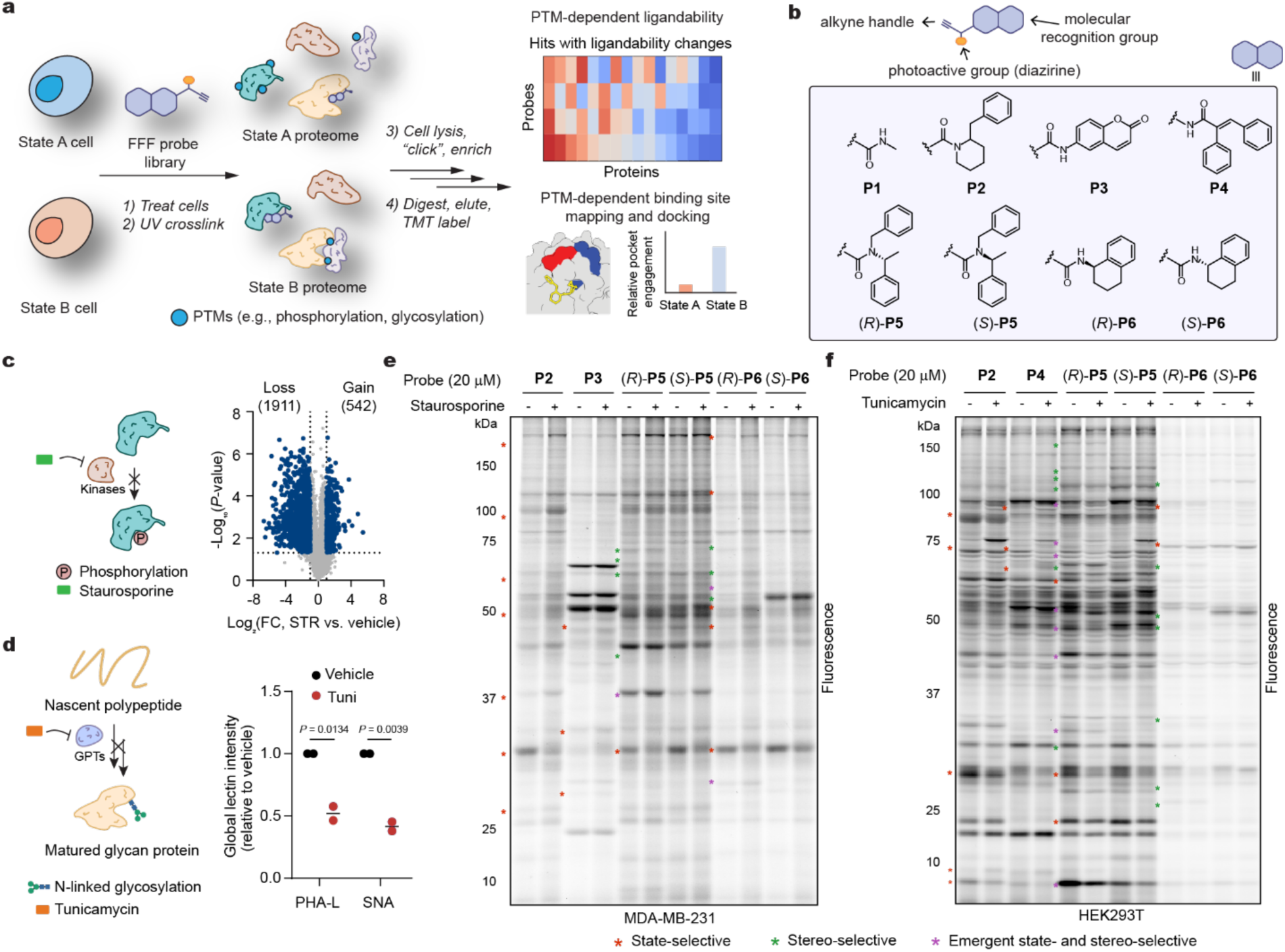
Chemoproteomic mapping of PTM-dependent ligandability changes in cells. **a**, Schematic of the experimental design for identifying PTM-dependent ligandability events. **b**, Structures of the fully-functionalized fragment (FFF) probe library used in this study, which consist of a small molecule fragment, diazirine photoreactive group and a clickable alkyne handle. **c**, Phosphorylation model system. MDA-MB-231 cells treated with staurosporine (STR, 250 nM, 3 h) result in 2,453 observed phosphorylation-related changes. **d**, N-glycosylation model system. HEK293T cells treated with tunicamycin (Tuni, 10 µg ml^−1^, 24 h) results in alterations in glycosylation as monitored with lectins PHA-L and SNA, which probe for the presence of complex-type N-linked glycans. PHA-L, *Phaseolus vulgaris* leucoagglutinin; SNA, *Sambucus nigra* agglutinin. **e**-**f**, Gel-based profiling of probe labeling under different phosphorylation states in MDA-MB-231 cells (**e**, soluble fraction) and N-linked glycosylation-dependent changes in HEK293T cells (**f**, soluble fraction). Red asterisks indicate representative state-selective probe–protein interactions, green asterisks denote stereoselective enantio-probe–protein interactions, and purple asterisks highlight interactions that are both state-selective and stereoselective. Proteomic and immunoblot quantifications were derived from two independent biological replicates and are presented as mean ± s.d. Statistical significance was assessed using two-tailed Student’s *t*-test.

## Results

### PTM-dependent ligandability profiling in human cells

To investigate changes in protein ligandability as a function of phosphorylation status, we aimed to broadly perturb phosphorylation while minimizing potential convoluting proteomic alterations. We found that treatment of MDA-MB-231 triple-negative breast cancer cells with staurosporine (STR), a pan-kinase inhibitor^21^, at low exposures (250 nM, 3 hours), leads to broad alteration of phosphorylation levels with minimal effects on cellular viability or changes in protein abundances (Fig. 1c, Extended Data Fig. 1a-d and Supplementary Data 1). Using MS-based quantitative phosphoproteomics, we monitored 28,000 phosphorylation modifications on serine, tyrosine, or threonine residues, and identified significant (>2-fold) decreases in phosphorylation at 1,911 sites and increases at 542 sites (Fig. 1c, Extended Data Fig. 1e-h and Supplementary Data 1).

We next set out to assess the impact of phosphorylation changes on protein ligandability in cells. Towards this end, we chose to profile a library of eight fully functionalized fragment (FFF) probes, comprising of a control probe (**1**), three nonchiral probes (**2**, **3**, **4**) and two pairs of enantiomerically paired probes ((*R*/*S*)-**5** and (*R*/*S*)-**6**) for target enrichment (Fig. 1b). These probes were chosen for their previously reported broad proteomic interactions, utility in identifying authentic ligandable sites on proteins and demonstration that they can be progressed into selective modulators of protein function^22–26^. MDA-MB-231 cells were treated with STR or vehicle, incubated with FFF probes (20 μM), and subsequently irradiated to capture probe-bound proteins. Harvested cells were lysed, fractionated, and conjugated to a rhodamine-azide tag using copper-catalyzed azide-alkyne cycloaddition (CuAAC) chemistry^27^, enabling visualization of probe-protein interactions via gel-based fluorescence scanning. We observed significant alterations in probe labeling, indicative of both decreased and increased binding across numerous proteins under STR treatment (Fig. 1e and Extended Data Fig. 2a). Notably, for the enantiomerically paired probes, in addition to numerous labeling events that were stereoselective, we also observed instances of “emergent” stereoselective binding events, where stereoselectivity is apparent only under one state but not the other.

To assess the impact of N-linked glycosylation on protein ligandability, we used tunicamycin (Tuni), a natural product which inhibits N–linked glycosylation through blockade of UDP-*N*-acetylglucosamine (GlcNAc) transfer to dolichol phosphate, a key step of N-linked protein glycosylation ^28^. Using HEK293T cells, which have been extensively characterized in previous glycosylation studies ^29^, we screened various conditions to maximize alterations in N-linked glycosylation while minimizing effects on cell viability (Fig. 1d and Extended Data Fig. 1i-j). We performed similar gel-based profiling studies with the FFF probes as described above, and, analogous to our phosphorylation perturbation, we observed numerous Tuni-dependent, stereo-selective and emergent stereo-selective probe-protein interactions across all the probes (Fig. 1f and Extended Data Fig. 2b), suggesting these conditions would be suitable to identify glycosylation-dependent ligandability events.

Next, we sought to identify and quantify phosphorylation-dependent ligandability events through MS-based proteomics. MDA-MB-231 cells were treated with STR followed by the FFF probes (100 μM) as described above. Probe-labeled proteins were then conjugated to an azide-biotin tag, isolated, and comparatively quantified by streptavidin enrichment and TMT-based quantitative proteomics ^24,25^. Proteins were considered probe targets if they exhibited >5-fold enrichment relative to control probe (**1**). Probe targets that display >2-fold change (*p* < 0.05) between conditions were classified as state-dependent ligandability events. Among 5,056 protein targets identified across all probes, 235 (5%) exhibited altered ligandability after STR treatment with at least one probe (Extended Data Fig. 3a, Supplementary Fig. 1a and Supplementary Data 1), with 107 (46%) possessing increased probe binding and 128 (54%) decreased probe binding. Similarly, we mapped N-linked glycosylation-dependent ligandability events in HEK293T cells. Using the filtering criteria established for phosphorylation, we identified a total of 4,421 probe targets, 225 of which exhibited significant ligandability changes (Extended Data Fig. 3b, Supplementary Fig. 1b and Supplementary Data 1), with 142 (63%) increased and 83 (37%) decreased in probe binding. By clustering the identified proteins based on their functional and clinical classifications^26^, we found that targets exhibiting state-dependent ligandability span a wide range of protein families (Fig. 2a, b and Supplementary Data 1). Most state-dependent targets were detected only by one or two probes (Fig. 2c), and were largely independent of changes in protein abundance (Extended Data Fig. 3c-e), indicating molecular recognition as a major driver of state-dependent ligandability events.

**Fig. 2.**
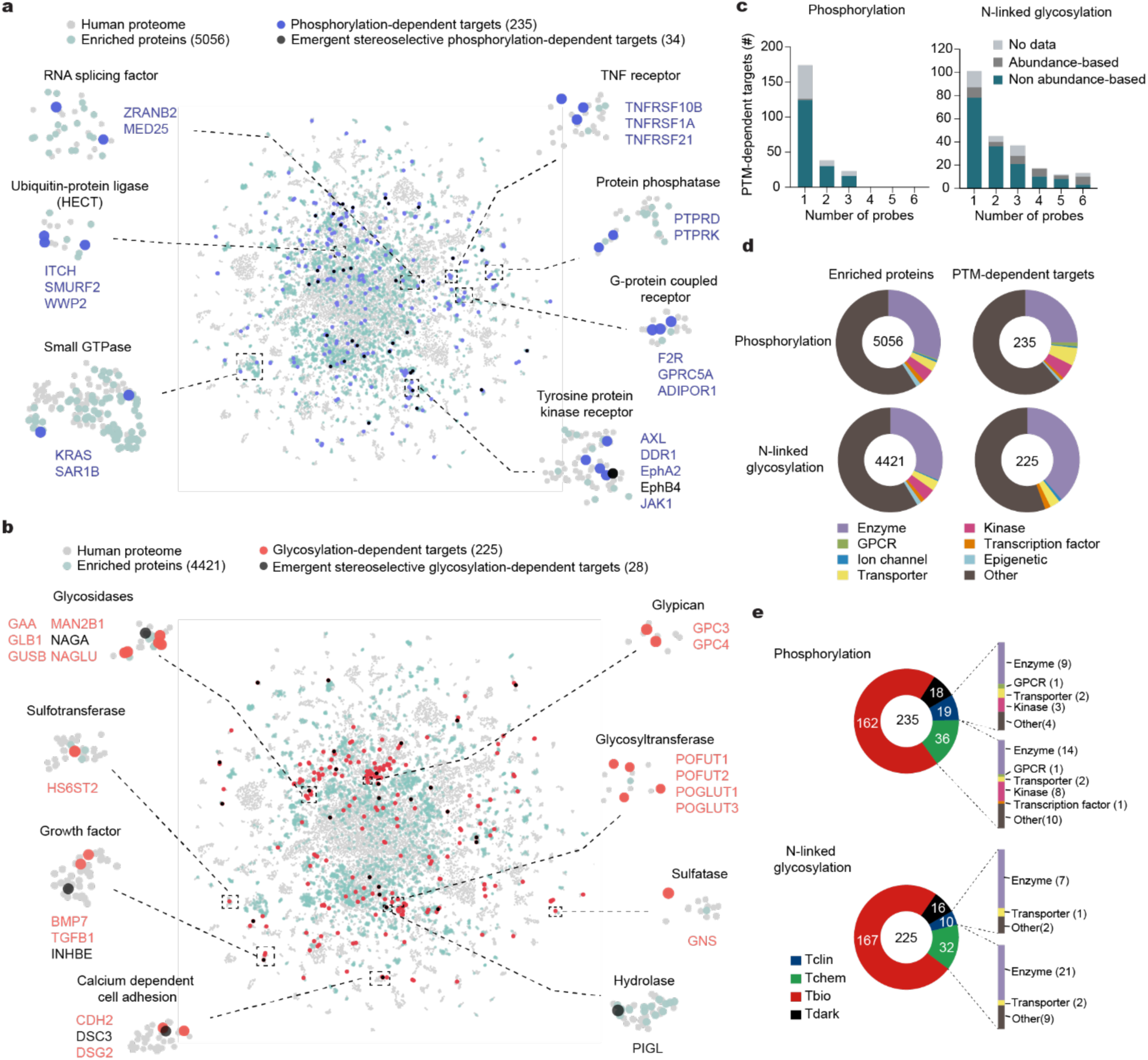
Proteome-wide ligandability changes associated with PTM status. **a**-**b**, Protein functional clustering of probe (100 µM) enriched proteins and PTM-dependent targets. Green dots represent probe enriched proteins (> 5-fold enrichment over control probe, *p* < 0.05, *n* = 2); Blue and red dots denote phosphorylation-dependent and glycosylation-dependent targets, respectively, defined by >2-fold difference between DMSO and STR (blue) or Tuni (red) treatments (*p* < 0.05, *n* = 2). Black dots represent proteins with state-specific stereoselective (emergent) interactions. **a**, phosphorylation-associated ligandability profiles. **b**, N-linked glycosylation-associated ligandability profiles. **c**, Number of PTM-dependent targets detected by number of probes. Dark grey corresponds to targets with detected protein abundance changes (> 2-fold, *p* < 0.05). Light grey represents proteins without abundance detection. **d**, Illuminating the Druggable Genome (IDG) functional classes of enriched protein and PTM-dependent targets. **e**, IDG Target Development Level (TDL) of PTM-dependent targets, based on four categories: Tclin (targets of approved drugs), Tchem (targets with known bioactive compounds), Tbio (biologically characterized targets lacking drug-like modulators), and Tdark (poorly characterized proteins with limited functional information). IDG Functional pathways highlighted for Tclin and Tchem categories. Associated datasets are provided in Supplementary Tables 1-4 and 6-7.

PTM-dependent targets include druggable protein classes such as enzymes and ion channels, but also historically more challenging targets, including epigenetic regulators and transcription factors (Fig. 2d) ^30^. Gene Ontology (GO) analysis revealed that proteins displaying PTM-dependent ligandability are enriched in a wide range of cellular processes. Among the top enriched pathways in phosphorylation-dependent targets were protein localization, lysosome organization, ion homeostasis, and mRNA transport, whereas saccharide metabolic processes and the ERAD pathway, were prominently enriched among N-glycosylation-dependent targets (Extended Data Fig. 3f). Critically, targets displaying differential ligandability were enriched (74% in phosphorylation and 62% in N-linked glycosylation) with proteins possessing the corresponding PTMs (Extended Data Fig. 3g), suggesting that many ligandability events emanate directly from changes in their PTM status. PTM-dependent targets include therapeutically relevant proteins (29, classified as Tclin), such as KRAS and HMGCR, and ligandable proteins (68, classified as Tchem), such as SOD1 and SLC2A1. However, the vast majority of PTM-dependent targets (∼78%) were found to lack well-annotated ligands (Fig. 2e).

### Features of proteins displaying PTM-dependent ligandability

We next sought to orthogonally validate the state-dependent interactions for 10 representative targets through probe pull down and immunoblotting. Among the STR-dependent targets, we confirmed the observed broad reduced ligandability of the tyrosine kinase JAK1 across multiple chemotypes (**P2,** (*R*)-**P6**, and (*S*)-**P6**) (Fig. 3a-b). In contrast, the E1 subunit of the V-ATPase proton pump, ATP6V1E1, despite being enriched by all probes, displayed increased ligandability exclusively with (*R*/*S*)-**P6**, suggesting that **P6** may possess a distinct binding mode that is affected by STR (Fig. 3c-d). Additional examples include the major protein kinase C substrate MARCKS (Extended Data Fig. 4a) and the transmembrane glycoprotein CD44 (Extended Data Fig. 4b), which displayed increased and decreased ligandability with **P2**, respectively, as well as the E3 ubiquitin-protein ligase WWP2, which exhibited decreased binding ability with (*R*/*S*)-**P6** following STR treatment (Extended Data Fig. 4c).

**Fig. 3.**
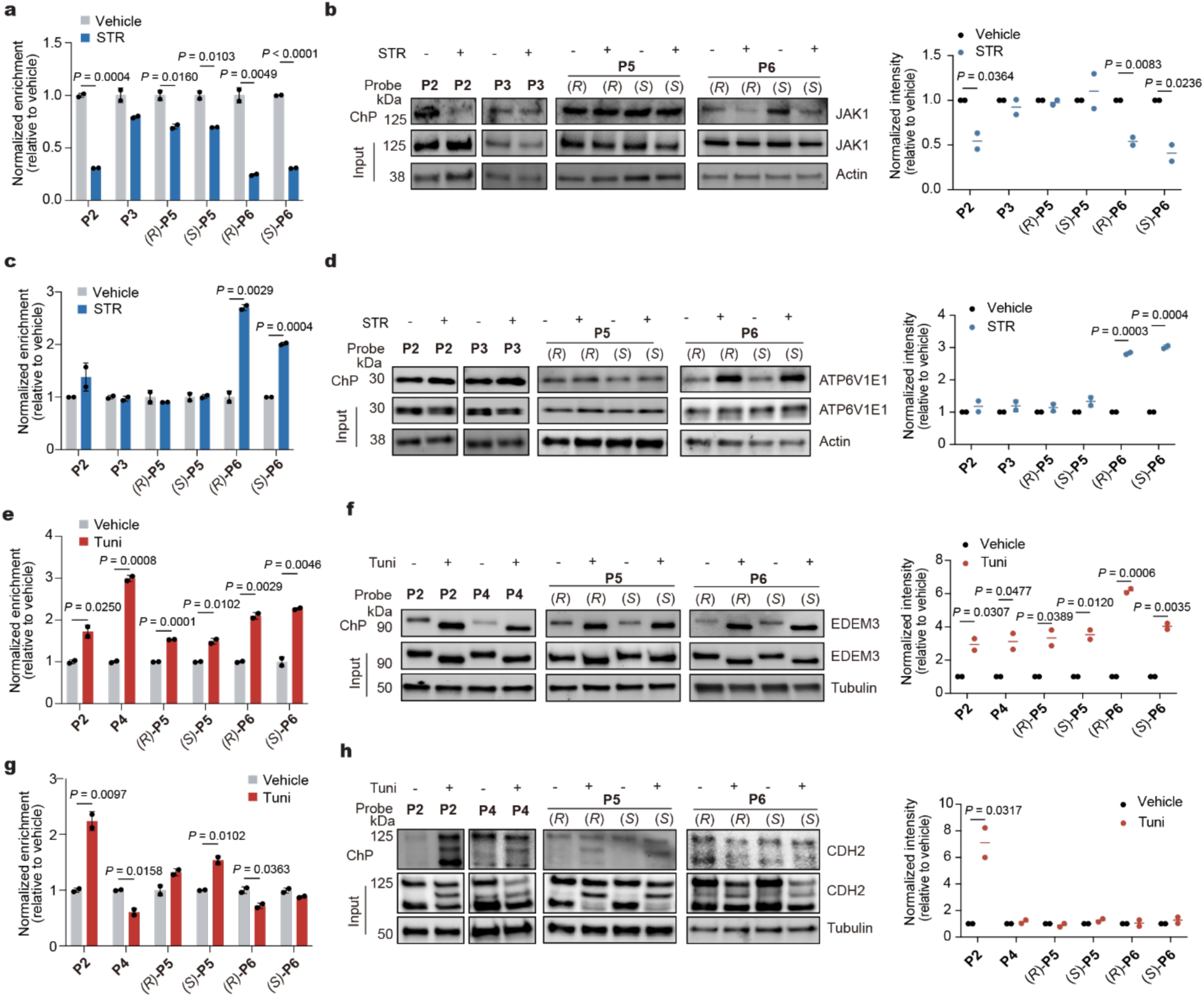
Analysis and validation of PTM-dependent ligandability changes. Proteomic profiles and corresponding chemo-precipitation (ChP) validation for selected targets exhibiting PTM-sensitive probe (100 µM) binding: JAK1 (**a-b**), ATP6V1E1 (**c-d**), EDEM3 (**e-f**) and CDH2 (**g-h**). Panels (**a**, **c**, **e**, **g**) show quantification of chemoproteomic profiling; panels (**b**, **d**, **f**, **h**) present validation by ChP (left) and correlated quantification across biological replicates (right). Proteomic data were obtained from two independent biological replicates and are presented as mean ± s.d. Gel images reflect representative results from two independent experiments. Statistical significance was assessed using two-tailed Student’s *t*-test. Associated datasets are provided in Supplementary Tables 1-8.

Among proteins displaying phosphorylation-dependent binding, 31% (72) possessed phosphosites quantified in our phosphoproteomic experiments, allowing for more in-depth analysis. Of these targets, 34 displayed changes in phosphorylation level and ligandability of the same directionality (Extended Data Fig. 4d-e). For example, we observed significantly reduced phosphorylation at S686, S697 and S706 on the cell surface receptor CD44, which also displayed a concomitant decrease in binding with probe **P2**, while we observed increase phosphorylation on S144 of chromatin associated protein NUCKS1 and increased binding with **P2**. We also observed 22 instances of ligandability changes that occur directionally opposite to observed phosphorylated changes. For example, the calmodulin-binding protein MARCKS, displayed increased binding with probe **P2** and decreased phosphorylation at S81 and T150. Among these targets, we identified 16 proteins without significant phosphorylation changes, including CLN3, a lysosomal protein with strong genetic links to Batten disease^31^, where all reported phosphorylation sites were quantified in our phosphoproteomic experiments but were unaltered. Such instances indicate that differences in probe labeling likely occur indirectly from changes in phosphorylation status of the liganded target.

Among the glycosylation-dependent targets, we confirmed broadly increased ligandability across all probes for EDEM3 (Fig. 3e-f), an ER-localized glycoprotein with seven annotated N-linked glycosites ^32^. We also validated CDH2, a cell membrane-localized adhesion glycoprotein (Fig. 3g-h) and DDOST, a subunit of the oligosaccharyl transferase (OST) complex, which in contrast to EDEM3, displayed selective increased binding to **P2** (Extended Data Fig. 5a) upon Tuni treatment. We noted a subset of targets (e.g., HSPA5, HYOU1, PDIA4) that were preferentially enriched by all six probes. Further inspection revealed that these events are largely related to changes in protein abundance and consist of members of the unfolded protein response (UPR), in-line with previous reports demonstrating that tunicamycin activates this pathway (Extended Data Fig. 5b)^33^. Therefore, we excluded these targets from further analysis. We also identified 21 functionally diverse proteins that exhibit differential ligandability under both phosphorylation and glycosylation perturbations, often in divergent directions (Extended Data Fig. 5c-d), underscoring the complexity of PTM influence over small molecule-protein interactions.

We and others have shown that integration of stereochemical features into probe design facilitates the confident assignment of authentic ligandable pockets on proteins ^23,34,35^. As preluded in our gel-based profiles (Fig. 1e-f and Extended Data Fig. 2), we identified 62 ‘emergent’ stereoselective liganding events across both PTM perturbations (Fig. 2a-b and Extended Data Fig. 6a-b), wherein stereoselective binding was found to occur in only one condition or state (34 in phosphorylation; 28 in glycosylation). Instances of emergent stereoselective events in the phosphorylation dataset include CETN2, HIGD1A and EphB4 (Extended Data Fig. 6c-e), as well as AS3MT, SLC25A4 and MPV17 (Extended Data Fig. 6f-h) in the glycosylation dataset. These data suggest that dynamic protein states can give rise to conditional stereoselective liganding events.

### Characterization of binding sites displaying PTM-dependent ligandability

We recently developed a chemical proteomic workflow to determine, with high resolution, proteome-wide binding sites of photoaffinity probes in cells and integrated this strategy with multiplexed quantitation ^24,36^. Using this method under the PTM perturbations described above, we identified differentially labeled binding sites (*p* < 0.05) in 82 and 55 proteins, respectively, from the group previously shown to exhibit significant phosphorylation- and glycosylation-dependent binding (Supplementary Data 1). With this higher resolution information in hand, we calculated the minimal spatial distance between labeled sequences and annotated PTM sites based on experimental or predicted structural information. This analysis revealed that, perhaps not surprisingly, binding sites exhibiting PTM-dependency are generally proximal to corresponding annotated PTM sites, suggestive that phosphate or N-glycan modifications may directly hinder or facilitate interactions between small molecules and proteins (Fig. 4a and Supplementary Data 1). We observe that differentially liganded sites often occurred at or near annotated functional sites (<10 Å), including active sites, ion-, cofactor- and nucleotide-binding sites, implicating potential roles of PTM status on protein function for these targets (Extended Data Fig. 7a).

**Fig. 4.**
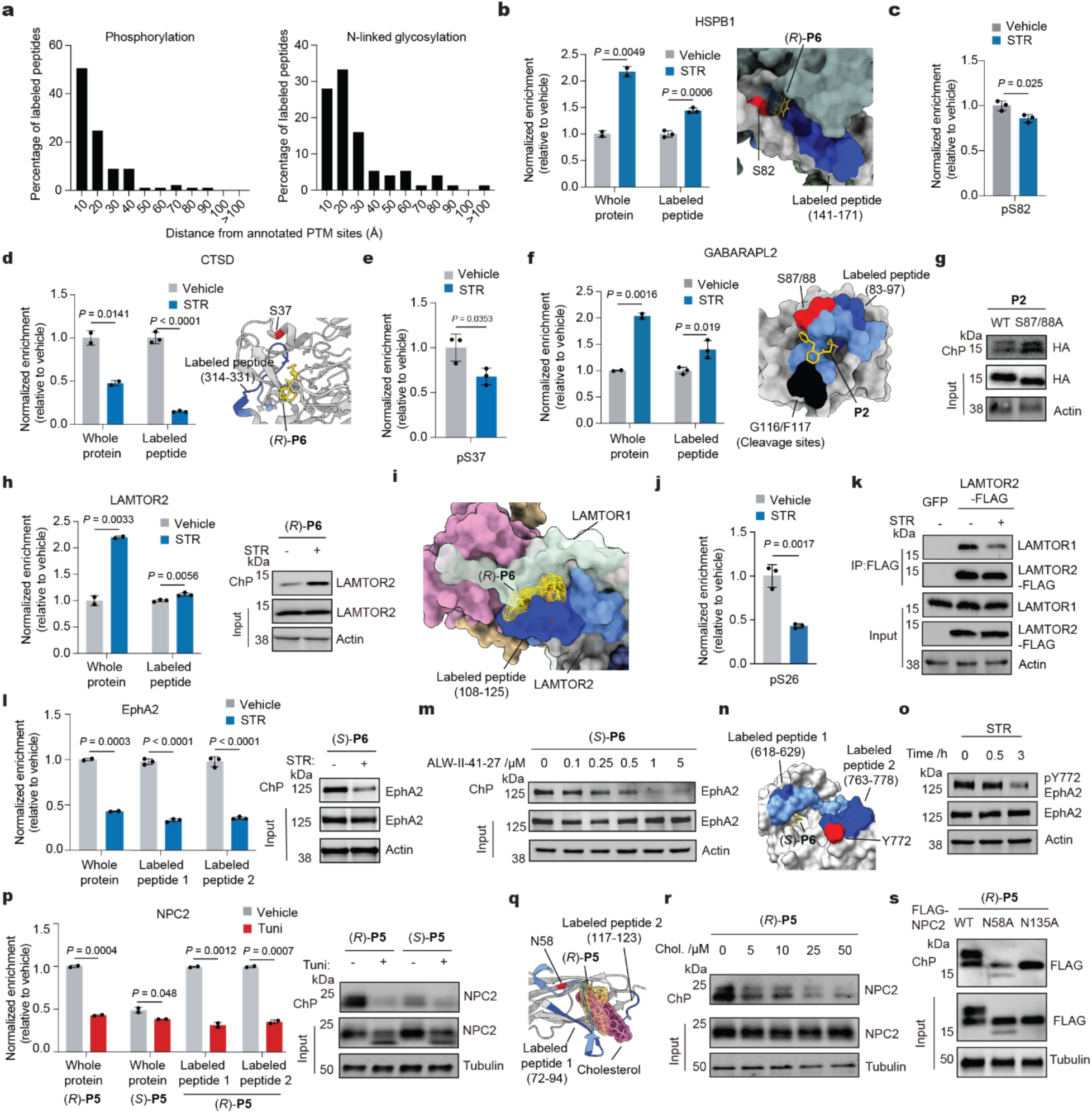
Mapping PTM-dependent binding sites in cells. **a**, Distribution of spatial distances between probe-labeled peptides and annotated phosphorylation (left) and N-linked glycosylation (right) sites. **b**, Proteomic profiling and structural docking reveal (*R*)-**P6** binding to a site on HSPB1 near S82 (PDB 6VD5). **c**, STR-induces S82 dephosphorylation in HSPB1. **d**, Proteomic profile of PTM-dependent liganding of CTSD with (*R*)-**P6** -labeled peptide (left) and docked structure in identified pocket near S37 (right, AF-P07339-F1). **e**, STR-induces dephosphorylation at S37 in CTSD. **f**, **P2** PTM-dependent labeling of a site near S87/88 on GABARAPL2 (PDB 7KL3). **g**, Chemo-precipitation of **P2** binding with GABARAPL2 with wild-type and S87/88A mutant GABARAPL2. **h**, Proteomic profile of PTM-dependent liganding (left) and ChP validation (right) of LAMTOR2 with (*R*)-**P6** after STR treatment. **i,** Docked structure of LAMTOR2 with (*R*)-**P6** (AF-Q9Y2Q5-F1), overlaps with the LAMTOR1 interface in the Ragulator complex (PDB 5X6V). **j**, STR-induces dephosphorylation at S26 in LAMTOR1. **k,** Immunoprecipitation (IP) reveals STR-induced reduction in LAMTOR1-LAMTOR2 complex formation. **l**, Proteomic profile of PTM-dependent liganding of EphA2 with (*S*)-**P6** -labeled peptide (left) and ChP validation (right) after STR treatment. **m**, Competitive inhibition of (*S*)-**P6** binding by ALW-II-41-27, an ATP-competitive EphA2 inhibitor. **n**, Docking analysis shows (*S*)-**P6** and ALW-II-41-27 share a binding site near ATP pocket of EphA2 (PDB 5EK7). **o**, STR-induces Y772 dephosphorylation in EphA2. **p**, Proteomic profile (left) and ChP validation (right) reveals PTM- and stereo-selective binding of (*R*)-**P5** to NPC2 with after Tuni treatment. **q**, Structural docking shows (*R*)-**P5** binds to the cholesterol-binding pocket of NPC2 near the N58 glycosylation site. **r**, Dose-dependent competition of (*R*)-**P5** by cholesterol. Chol., cholesterol. **s**, Mutation of N58 in NPC2 disrupts (*R*)-**P5** binding. Phosphorylation experiments were conducted in MDA-MB-231 cells with or without staurosporine treatment (250 nM, 3 h). Glycosylation experiments were conducted with or without tunicamycin (10 µg ml^−1^, 24 h) in HEK293T cells. All structures depict probe-labeled residues in dark blue, the remainder of each peptide in light blue, annotated PTM sites in red, functional residues in black, and probes in gold. Proteomic data represent mean ± s.d. from two or three independent biological replicates. Gel images are representative of two independent experiments. Significance was determined using a two-tailed Student’s *t*-test. Associated datasets are provided in Supplementary Tables 11-12.

We noted that differentially liganded sites largely overlapped with predicted small molecule binding pockets (Extended Data Fig. 7b). To begin to better understand molecular recognition components that might influence probe binding, we integrated our state-dependent site mapping data with molecular docking. Towards this end, we obtained atomic structures for a subset of interactions (57 structures from PDB and 45 from AlphaFold), then blindly docked every probe into every predicted pocket and cross-referenced the results to the proteomics data as previously reported ^24^. We identified low-energy probe binding modes across a variety of pockets proximal to corresponding PTMs. Among these PTM-dependent pockets include diverse functional sites, such as ATP binding sites (TGFBR1; Extended Data Fig. 7c), calcium binding sites (NUCB2, CALM3; Extended Data Fig. 7d-e) and phosphosites (COPE, HMGB1, CD63; Extended Data Fig. 7f-h) in phosphorylation-dependent pockets; protease cleavage sites (HBA1; Extended Data Fig. 7i), substrate binding site (HS6ST2; Extended Data Fig. 7j), enzyme active sites (CTSB; Extended Data Fig. 7k) and glycosites (OSTM1; Extended Data Fig. 7l) in glycosylation-dependent pockets.

In several instances, we also detected and quantified corresponding phosphorylation changes from our phosphoproteomic studies, allowing us to directly compare alterations in ligandability with changes in phosphorylation. We identified 20 binding pockets with altered ligandability located within 10 Å of phosphosites whose phosphorylation level changed upon STR treatment. Of these, 6 exhibited ligandability changes that correlated with phosphorylation changes, while the other 14 showed the opposite trend (Extended Data Fig. 7m). For instance, we observed selective increased binding of (*R*)-**P6** with the heat shock protein HSPB1 (Fig. 4b) in a pocket proximal (∼5 Å) to the S82, which decreases in phosphorylation (Fig. 4c), suggesting that the phosphate group may directly obstruct probe binding. In contrast, for CTSD, both (*R*)-**P6** binding (Fig. 4d) and phosphorylation at the nearby S37 site (∼6 Å) decreased (Fig. 4e), indicating that the phosphate group may, in some cases, facilitate probe binding. We also observed increased **P2** binding to a pocket near S87/88 on GABARAPL2, phosphosites implicated in GABARAPL2’s autophagic roles^37^ (Fig. 4f). Mutation of these residues to alanine (S87/S88A) to block phosphorylation led to substantial increased **P2** binding, confirming the impact of phosphorylation on the ligandability of this site (Fig. 4g).

In addition to binding pockets proximal to PTMs themselves, we also identified several differentially liganded sites that occur near protein-protein interfaces. For instance, we observed increased (*S*)-**P5** binding to a pocket that is located at the interface of the GSTP1 homodimer and proximal to the glutathione binding site (Extended Data Fig. 7n). Interestingly, phosphorylation of GSTP1 shifts the dimer-monomer equilibrium and promotes the formation of the GSTP1-JNK complex ^38^, suggesting that dephosphorylation may result in a formation of a neopocket at the dimer interface that promotes probe binding. We also observed an increase in (*R*)-**P6** binding of the metalloprotease inhibitor TIMP3 following STR treatment (Extended Data Fig. 7o), though TIMP3 is itself not annotated to be phosphorylated. Binding site inspection and molecular docking revealed, though we identified a differentially probe modified peptide on TIMP3, (*R*)-**P6** likely binds to a pocket on TNF-α converting enzyme (TACE), whose activity is inhibited upon TIMP3 binding ^39^. Interestingly, inhibition of the ERK/p38 MAPK pathway increases the association between TIMP3 and TACE, presumably through modulation of TACE phosphorylation^40^. Consistent with this, we observed a significant decrease in TACE phosphorylation at S791 (Extended Data Fig. 7p), suggesting increased (*R*)-**P6** could emanate from increased stability of this complex. In another notable example, we discovered increased binding of (*R*)-**P6** to LAMTOR2, a subunit of the Ragulator-Rag complex ^41^, upon STR treatment (Fig. 4h and Extended Data Fig. 7q). Site mapping and docking analysis revealed that the probe likely binds to a pocket on LAMTOR2 that is occupied by LAMTOR1 in Ragulator complex (Fig. 4i). Though several phosphosites have been reported in the Ragulator complex, primarily on LAMTOR1, we observed decreased phosphorylation only at S26 (Fig. 4j), which is not located near (*R*)-**P6** labeled site, prompting us to investigate whether STR treatment may impact complex stability. Indeed, we observed substantially reduced interactions between LAMTOR1 and LAMTOR2 upon STR treatment (Fig. 4k), implying that ligandability differences under these conditions result from disruption of the Ragulator complex, exposing a binding pocket. From the glycosylation dataset, we observed an increased binding of **P2** to calreticulin (CALR), a chaperone with specificity for monoglucosylated oligosaccharides, such as those of HLA-A, a component of the MHC I complex^42^. Notably, **P2** site mapping on CALR revealed increased binding at the glycoprotein-interaction region, and structural docking further showed that **P2** occupies the same pocket as the N-linked glycan on N110 of HLA-A (Extended Data Fig. 7r), suggesting that it may obstruct probe binding to CALR.

As previously noted, we observed that a subset of proteins exhibited ligandability changes under both the STR and Tuni treated conditions (Extended Data Fig. 5c-d). We generally observe that these events occur at distinct binding sites, however, we identified instances where the same pocket displayed opposing ligandability changes depending on the treatment. For example, we mapped the binding site of both **P2** and (*S*)-**P6** to the same pocket near the intracellular death domain of TNFRSF10B (DR5), the receptor for the cytotoxic ligand TNFSF10/TRAIL (Extended Data Fig. 7s). Upon treatment with STR, we observe substantially decreased binding to **P2**; however, we also observed increased binding with (*S*)-**P6** upon treatment with Tuni (Extended Data Fig. 7s). Interestingly, though there are no Uniprot-annotated PTMs on either TNFRSF10B or TNFSF10, both possess Asn-X-Ser/Thr (X = any amino acid but Pro) canonical N-glycosylation sequons as well as predicted phosphosites ^43^, alluding to their existence^44^. More broadly, these observations highlight that pocket ligandability can be highly dynamic and that a specific PTM status can impose differential effects on pocket ligandability.

Among mapped binding pockets displaying PTM-dependent ligandability, several have known ligands, leading us to inquire whether changes in PTM status could affect binding of these ligands. For example, we identified a selective decrease in binding of probe (*S*)-**P6** with the receptor tyrosine kinase EphA2 in the presence of STR (Fig. 4l). We identified two (*S*)-**P6** labeled peptides on EphA2 located near the ATP binding pocket and binding site of ALW-II-41-27 (Extended Data Fig. 8a), an ATP-competitive EphA2 inhibitor ^45^. We first validated this finding via immunoblotting (Fig. 4l and Extended Data Fig. 8b) and confirmed that the observed decrease in probe binding was not due to competitive binding with STR itself (Extended Data Fig. 8c). However, co-incubation of (*S*)-**P6** with increasing concentrations of ALW-II-41-27, revealed clear blockade of probe labeling, indicating they target the same pocket (Fig. 4m), which was further supported by binding site mapping and molecular docking (Fig. 4n). Interestingly, it was reported that ALW-II-41-27 inhibitory activity decreases when Y772 on EphA2 is dephosphorylated^46^, aligning with our observation that STR treatment lowers Y772 phosphorylation and correspondingly decreases (*S*)-**P6** binding (Fig. 4o).

Among the targets displaying glycosylation-dependent ligandability changes, we identified NPC intracellular cholesterol transportation 2 (NPC2) (Fig. 4p). NPC2 plays a pivotal role in facilitating the egress of lipoprotein-derived cholesterol from lysosomes, and mutations of this gene, along with NPC1, are associated with Niemann-Pick C disease, an autosomal recessive disorder ^47^. We observed a stereoselective, decreased binding with (*R*)-**P5** in Tuni-treated cells and observed no changes with the enantiomer (*S*)-**P5** (Fig. 4p and Extended Data Fig. 8d). Binding site mapping and docking of (*R*)-**P5** on NPC2 revealed engagement within the cholesterol binding pocket (Fig. 4q). In line with this observation, co-incubation with increasing concentrations of cholesterol resulted in a dose-dependent decrease in (*R*)-**P5** labeling (Fig. 4r and Extended Data Fig. 8e). First, we suspected that reduced glycosylation might affect NPC2 lysosomal localization, however, we observed limited changes in localization upon Tuni treatment (Extended Data Fig. 8f). Two primary N-linked glycosylation sites on NPC2 have been reported at N58 and N135 ^48^. Inspection of the NPC2 structure indicates that N58 is proximal to the mapped (*R*)-**5** site and the cholesterol binding pocket (Fig. 4q), prompting us to hypothesize that N58 glycosylation might directly impact probe binding. Indeed, we observe a substantial decrease in (*R*)-**P5** binding to the N58A NPC2 mutant but not the N135A mutant (Fig. 4s), suggesting that N-glycosylation on N58 of NPC2 may play a crucial role in small molecule and substrate binding of this pocket.

### Phosphorylation status alters KRAS inhibitor activity

Among targets with functional ligands displaying phosphorylation-dependent ligandability, we identified a significant reduction of **P3** binding to the small GTPase KRAS (Fig. 5a, Extended Data Fig. 9a and Supplementary Data 1). RAS family proteins, including KRAS, are molecular switches, receiving signals from growth factor receptors and relaying them to downstream effector pathways, which regulate processes like cell growth and proliferation ^49^. Somatic mutations on KRAS are the most common activating lesions found in cancer cells, with the most frequent occurring at codons 12 (G12) 13 (G13) and 61 (Q61), resulting in abnormal activation of the RAS pathway ^50^. As a key oncogene, substantial efforts have been made over the past decades to drug KRAS, particularly targeting oncogenic variants ^49,51^.

**Fig. 5.**
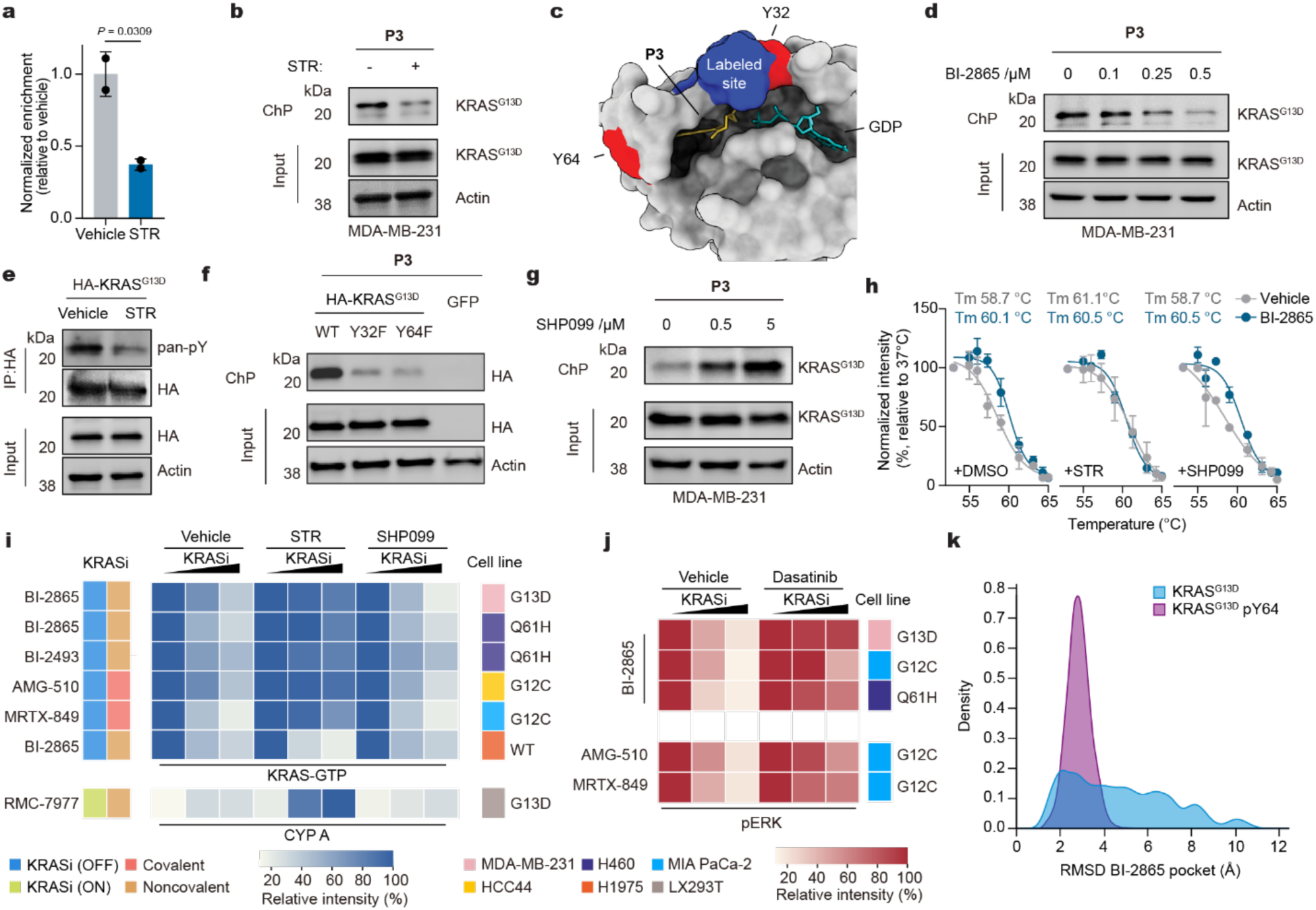
KRAS phosphorylation status alters druggability. **a**-**b**, Proteomic profile (**a**) and chemo-precipitation validation (**b**) of **P3** binding to KRAS^G13D^ following STR treatment (250 nM, 3 h) in MDA-MB-231 cells. **c**, Docked structure of KRAS with **P3** highlights probe interaction near the switch-II pocket (PDB 8B00). Residues labeled by **P3** are highlighted in blue, phosphorylation sites (Y32 and Y64) in red, **P3** in yellow, and GDP in cyan. **d**, Dose-dependent competition between **P3** and the pan-KRAS inhibitor BI-2865 in MDA-MB-231 cells. **e**, KRAS phosphorylation level decreased following treatment with staurosporine (250 nM, 3 h). HA-tagged KRAS was immunoprecipitated, phosphorylation status was assessed by immunoblotting using a pan-phosphotyrosine (pY) antibody. **f**, ChP shows reduced **P3** binding to Y32F and Y64F mutants of KRAS^G13D^. **g,** Increased **P3** engagement with KRAS^G13D^ upon SHP2 inhibition by SHP099 (500 nM, 30 min) in MDA-MB-231 cells. **h**, Cellular thermal shift assay (CETSA) reveals decreased KRAS stabilization by BI-2865 in the presence of STR, while SHP099 has no observable effect. **i**, Ras-binding domain (RBD) assay shows that STR diminishes the ability of KRAS OFF-inhibitors to disrupt the active (GTP-bound) KRAS-RAF interaction across multiple KRAS-mutant cell lines, while SHP099 restores the effect. Immunoprecipitation (IP) reveals that STR enhances KRAS-CYPA ternary complex formation in the presence of the KRAS ON-state inhibitor RMC-7977. **j**, Total pERK (T202/Y204) levels assessed by immunoblotting in cells co-treated with dasatinib (1 µM) and indicated KRAS inhibitors restores pERK across various KRAS mutations in multiple cell lines. **k**, Density plots showing the distribution of BI-2865 binding competent conformations. Results are representative of two biological replicates. Proteomic and immunoblotting data are presented as mean ± s.d. from two or three independent biological replicates. Statistical significance was determined using a two-tailed Student’s *t*-test. See Supplementary Tables 2 and 13 for related datasets.

We first confirmed the identified **P3** ligandability change against both endogenous KRAS^G13D^ in MDA-MB-231 cells^52^ (Fig. 5b and Extended Data Fig. 9b) and recombinantly expressed HA-KRAS^G13D^ in Lenti-X 293T cells (Extended Data Fig. 9c). We observe similar phosphorylation-dependent engagement with G12C and Q61H mutants (Extended Data Fig. 9c-e). Interestingly, we observed directionally opposite phospho-dependent **P3** ligandability changes on both endogenous and recombinantly expressed KRAS^WT^ relative to the oncogenic mutants (Extended Data Fig. 9f). We mapped the binding site of **P3** to be near the switch-II (SWII) pocket (Fig. 5c), the binding site of many KRAS inhibitors ^53^, including the FDA-approved AMG-510 (Sotorasib) and MRTX-849 (Adagrasib) which covalently target the G12C, as well as the non-covalent pan KRAS inhibitor BI-2865 ^54–56^. Molecular docking yielded a high ranking **P3** binding pose that overlaps with several KRAS inhibitors (Extended Data Fig. 9g-h), which was subsequently confirmed by competition experiments in cells with BI-2865 and AMG-510, where both inhibitors dose-dependently block probe binding in the concentration range of their reported potencies (Fig. 5d and Extended Data Fig. 9i). Like these SWII inhibitors, we observed that GppNHP, a non-hydrolyzable analog of GTP, blocks **P3** binding to KRAS^G13D^, while GDP had no observable effect, demonstrating that **P3** preferentially binds to the GDP-bound state (Extended Data Fig. 9j).

Further inspection of **P3**’s binding mode revealed its proximity to Y32 and Y64 (<10 Å), raising the possibility that phosphorylation of these sites may play a role in the observed differential binding. To explore this hypothesis, we first confirmed that treatment with STR led to dephosphorylation of KRAS^G13D^ (Fig. 5e). We next generated dephosphorylated mimicking Y32F and Y64F mutants and observed ablation of **P3** binding to both G13D and G12C mutants, consistent with the effects observed with STR treatment of tyrosine counterparts (Figure 5f and Extended Data Fig. 9k). Furthermore, pharmacological inhibition of the KRAS phosphatase SHP2 with SHP099 ^57,58^ resulted in a modest increased interaction between KRAS and **P3** in both MDA-MB-231 (KRAS^G13D^) cells (Fig. 5g) and HCC44 (KRAS^G12C^) cells (Extended Data Fig. 9d). Conversely, inhibition of c-Src kinase with dasatinib, which phosphorylates KRAS at Y32 and Y64 ^58,59^, resulted in a similar reduction of engagement (Extended Data Fig. 9l). Collectively these data confirm that phosphorylation status at Y32 and Y64 impact **P3** binding near the SWII pocket.

We next wondered whether phosphorylation would also affect the interaction between KRAS and SWII pocket inhibitors. We first employed cellular thermal shift assays (CETSA) ^60^ to evaluate how inhibitor impact on KRAS stability might change upon STR treatment. We observed a significant thermal stabilization of KRAS in MDA-MB-231 cells treated with BI-2865, however, this stabilization was effectively diminished when the cells were co-treated with STR or dasatinib (Fig. 5h and Extended Data Fig. 9m-n). In contrast, treatment with SHP099 restored and even enhanced the stabilizing effects (Fig. 5h and Extended Data Fig. 9m), indicating that KRAS phosphorylation status influences the inhibitor-mediated thermal stabilization. SWII inhibitors block interactions with c-Raf ^56,61,62^, prompting us to hypothesize that phosphorylation status would also impact this activity. We observed a dose-dependent disruption of the RAS-RAF interaction in cells treated with BI-2865, however, this disruption was ablated upon STR treatment (Fig. 5i and Extended Data Fig. 10a), an effect that was observable for at least 8 hours (Extended Data Fig. 10b). Critically, BI-2865 did not disrupt the KRAS-RAF interaction in Y32F (KRAS^G13D-Y32F^) and Y64F (KRAS^G13D-Y64F^) mutants ^61^ (Extended Data Fig. 10c) consistent with the hypothesis that KRAS phosphorylation at these sites impacts inhibitor binding. Notably, similar patterns were observed with another non-covalent inhibitor (BI-2493) as well as covalent inhibitors (AMG-510 and MRTX-849), wherein the activity of these inhibitors were effectively abolished in cells treated with STR (Fig. 5i and Extended Data Fig. 10a) or dasatinib (Extended Data Fig. 10d-e). We observe similar effects with BI-2865 or BI-2493 treated H460 cells harboring the Q61H mutation, which has impaired intrinsic and GAP-mediated GTP hydrolysis^63–65^, suggesting the effects of phosphorylation on inhibitor binding may not be fully attributed to GDP-GTP cycling (Extended Data Fig. 10a).

Intriguingly, treatment of cells with RMC-7977, which targets GTP-bound KRAS via ternary complex with cyclophilin A (CYPA) to block the RAS-RAF interaction ^66^, led to an increase in CYPA binding upon STR treatment as did KRAS^G13D-Y32F^ and KRAS^G13D-Y64F^ mutations (Fig. 5i and Extended Data Fig. 10g-h). These findings suggest that phosphorylation plays a distinct role in modulating the activity of KRAS-OFF and -ON inhibitors. In line with our observations with **P3** binding, this effect appears to be specific to mutant KRAS, as STR had no effect on BI-2865’s ability to disrupt the KRAS-RAF interaction in H1975 cells, which express KRAS^WT^ (Fig. 5i and Extended Data Fig. 10a). Finally, we observed a restoration of downstream phospho-ERK signaling when KRAS G12, G13 or Q61 mutant cells are co-treated with KRAS inhibitors and dasatinib, demonstrating that phosphorylation status influences both inhibitor binding and functional suppression (Fig. 5j and Extended Data Fig. 10i).

Phosphorylation of Y32 and Y64 has been recently shown to impact KRAS structural conformation, impacting interactions with the effector RAF and GTP cycling ^58,63,67^, suggesting that the observed ligandability differences are, in part, consequence of phosphorylation affecting the SWII binding site conformation. We performed enhanced sampling molecular dynamics simulations, which revealed that phosphorylation at Y64 in GDP-bound KRAS^G13D^ significantly alters conformational dynamics, decreasing SWII flexibility while increasing SWI flexibility (Extended Data Fig. 10j). This shift in conformational equilibrium enhances the prevalence of BI-2865 binding-competent states in phosphorylated KRAS^G13D^ compared to its non-phosphorylated counterpart (Fig. 5k). To perform a more accurate energetic comparison, we employed metadynamics simulations to obtain Free Energy Surfaces (FES) for these KRAS variants, including also the Y32-phosphorylated proteoform. Phosphorylation either at Y32 or Y64 induces a drastic shift in the conformational population, reshaping the energy landscape in a way that essentially locks KRAS^G13D^ in a closed-like state that is compatible with BI-2865 binding, supporting the observed ligandability changes (Extended Data Fig. 10k).

## Discussion

While previous strategies have systematically cataloged PTM prevalence and dynamics through mass spectrometry^5^, translating such high dimensional datasets into actionable knowledge regarding protein function, chemical tractability, or druggability, has remained relatively underexplored^68^. To date, only a limited number of examples have been identified—often through serendipitous discovery—in which PTMs modulate protein ligandability. For instance, the unphosphorylated form of Abelson tyrosine kinase (Abl) exhibits over 150-fold greater sensitivity to inhibition by the clinically approved drug imatinib and acetylation cyclophilin A substantially antagonizes the immunosuppressive effects of the natural product cylosporin.^69,70^ Similarly, α1-acid glycoprotein (AGP) displays highly dynamic binding affinities for the anticoagulant warfarin across various glycoforms^71^. Our chemical proteomic approach directly addresses this gap by enabling proteome-wide assessment of how phosphorylation, N-linked glycosylation, and, in principle, any PTM can influence protein–small molecule interactions.

The implications of our findings extend broadly across biological and therapeutic contexts. We revealed that PTM-dependent ligandability events involve diverse functional classes, including traditionally challenging drug targets such as transcription factors and epigenetic regulators. Our data point to a wide range of mechanisms, beyond abundance changes, by which PTMs may affect protein ligandability. We observe a strong enrichment of differentially liganded sites located proximal to the corresponding altered PTM (Fig. 4a-g), suggesting that the PTM directly shapes ligand-binding pockets, either by altering pocket accessibility orthosterically or influencing conformational dynamics critical for ligand engagement. We also identify differentially ligand sites that are not located near PTMs, suggesting PTM alterations impact binding indirectly (Extended Data Fig. 7m-s), such as the case of LAMTOR2 where STR treatment appears to disrupt Ragulator complex formation, creating a ‘neo-pocket’ (Fig. 4h-k). Moreover, the observed PTM-dependent alterations in ligandability are often proximal to annotated functional pockets (Extended Data Fig. 7a-k), including enzymatic active sites, substrate-binding pockets, protein-protein interaction interfaces, as well as functionally unannotated sites. This close alignment of ligandability changes with functional domains suggests that ligandability can be a powerful proxy for protein status—reporting shifts in conformation, interactions, and other functions beyond direct small molecule binding.

Critically, our study also identified clinically relevant targets, such as KRAS, whose phosphorylation state profoundly influences inhibitor efficacy. This observation raises important questions regarding how chemical probes and clinically approved drugs behave in native systems under different states. Multiple studies have shown that SHP2 inhibitors impair growth and induce death of KRAS-driven cancers and the combination of SHP2 inhibitors with KRAS inhibitors have resulted in enhanced efficacy in clinical settings ^72,73^. These findings have been attributed to synergistic mediation of upstream RTK signaling and stimulation of KRAS GTP cycling *via* GEF/GAP modulation ^58,74–82^. Extensive work has also shown that phosphorylation of KRAS directly stalls the GTPase cycle and impairs effector binding ^58,63,67^. Though our observation that phosphorylation has similar impacts to cells harboring G12/G13 mutants, as well as the Q61H mutant, suggests additional factors mediated by KRAS phosphorylation may contribute to the observed effects of ligand binding to the SWII pocket, such as conformational changes, as supported by our modeling studies (Fig. 5k and Extended Data Fig. 10j-k). We further observed that WT KRAS exhibits opposing phosphorylation-dependent ligandability patterns compared to oncogenic mutants and RMC-7977-mediated KRAS-CYPA stabilization is enhanced when KRAS is dephosphorylated, demonstrating a multi-layered regulation of pocket activity. These results underscore that PTM-driven modulation of ligandability may be consequential for therapeutic targeting, and such PTM-dependent nuances could unveil new opportunities to optimize existing therapies tailored to specific cellular states.

It is important to note several limitations of our study. Though we observed 400+ PTM-dependent ligandability events, our analysis was restricted to changes in phosphorylation and N-linked glycosylation mediated by staurosporine and tunicamycin, respectively, and thus reflects a relatively limited perspective of how PTMs and other factors might influence proteome-wide ligandability. Additionally, all experiments were conducted in HEK293T or MDA-MB-231 cells, confining our coverage to proteins expressed in these cells. We anticipate that extending this approach to additional PTMs, cellular states, and disease-relevant contexts as well as broadening such investigations to diverse tissues, will further illuminate how protein status shapes ligandability and druggability. Moreover, while we employed relatively promiscuous fragment probes, our library is small and inadequately covers broad chemical space. Future studies employing larger libraries with structurally diverse and stereochemically defined members should enable deeper examination of the proteome and improve the likelihood of identifying state-dependent ligandability events that can be advanced into chemical probes capable of modulating protein function in a state-specific manner. Collectively, our approach provides not only a new format to investigate how PTMs modulate protein status but also lays a general framework for the discovery of chemical probes to target proteins in specific functional or pathological states.

## Methods

### Cell culture and treatment

MDA-MB-231, HEK293T, H460, MIA PaCa2 and H1975 cell lines were procured from ATCC, while HCC44 cells were obtained from DSMZ, and Lenti-X™ 293T (LX293T) cells from Takara Bio. MDA-MB-231, MIA PaCa2, HEK293T, and Lenti-X™ 293T cells were cultured in Dulbecco’s Modified Eagle Medium (DMEM) supplemented with 10% (*v*/*v*) fetal bovine serum (FBS) and 2 mM L-glutamine. HCC44, H1975, and H460 cells were maintained in RPMI-1640 medium supplemented with 10% (*v*/*v*) FBS and 2 mM L-glutamine. All cell lines were incubated at 37 °C in a humidified atmosphere containing 5% CO_2_. Unless otherwise indicated, all compound treatment experiments were performed when cells are around 50-70% confluency.

### Chemical probes and materials

The chemical probes and acid cleavable biotin azide used in this study were synthesized in-house based on previously reported research from our lab^23,83,84^. Commonly used reagents were purchased from Corning, Gibco, Bio-Rad, Research Product International, Sigma-Aldrich and ThermoFisher Scientific. Biotin-PEG3-Azide (AZ104) was from Click Chemistry Tools. Sequencing grade modified trypsin (V5111) was purchased from Promega. Endoproteinase LysC (P8109S) is from New England Biolabs, Inc. Pierce streptavidin agarose (catalog no. 20353), Pierce high pH reversed-phase fractionation kit (84868), TMT10Plex isobaric label reagent set (90406), TMTpro 16plex label reagent set (A44520), halt protease and phosphatase inhibitors (78438 or 78441) were from ThermoFisher Scientific. Specific chemical reagents originated from the following sources: staurosporine (Selleckchem, S1421); ALW-II-41-27 (HY-18007), BI-2493 (HY-153723), BI-2865 (HY-153724), AMG-510 (HY-114277), MRTX-849 (HY-130149), RMC-7977 (HY-156498), SHP-099 (HY-100388), dasatinib (HY-10181) are from MedChemExpress; tunicamycin (Merck, T7765); cholesterol (Avanti Research, 700100).

### Plasmids and transient transfection

The full-length codon-optimized human NPC2, including (WT, N58A, or N135A mutation), and human GABARPL2 (WT and S87/88A) were synthesized (Twist Bioscience) and cloned into a pTWIST-CMV-puro vector individually. pBabe-HA-KRAS (75282), pRK5-p14-Flag (42330) was obtained from Addgene. Site-directed mutagenesis was generated by Phusion Flash High-Fidelity PCR (Thermo Fisher, F548S) and ligated by In-Fusion Snap Assembly (Takara Bio, 638947). All plasmids were verified by Sanger or whole plasmid sequencing. Primers for mutagenesis were as follows (5’ to 3’ orientation):

G12C: forward (F):TGGAGCTTGTGGCGTAGGCAAGAGTGCC,

reverse (R): ACGCCACAAGCTCCAACTACCACAAGTTT;

G13D, F: AGCTGGCGATGTAGGCAAGAGTGCCTTGACGA,

R: CCTACATCGCCAGCTCCAACTACCACAAGT;

Y32F, F: GGACGAATTTGATCCAACAATAGAGGATTCCTACA,

R: GGATCAAATTCGTCCACAAAATGATTCTGA;

Y64F, F: AGAGGAGTTTAGTGCAATGAGGGACCAGTAC,

R: GCACTAAACTCCTCTTGACCTGCTGTGTCG.

For transient transfection, LX293T cells were seeded and transfected by Lipofectamine 3000 Transfection Reagent (Invitrogen, L3000075). Cells were harvested at 24 or 48 hours post-transfection for subsequent analysis.

### Proliferation assay

Cells were plated at 20,000 cells per well in 96-well plates. The media was first aspirated after the cells were attached, and the cells were then treated in triplicate with media containing different concentrations of staurosporine (24 nM-50 µM) for MDA-MB-231 cells and tunicamycin (1-100 µg ml^−1^) for HEK293T cells. The same volume of DMSO was used as a vehicle control. Cell proliferation was assayed using the CellTiter-Glo Luminescent Cell Viability Assay (Promega) following manufacturer’s guidelines. Data represents the average and standard deviation of triplicates in measured luminescence.

### Photoaffinity probe cell treatments

For MDA-MB-231 cells, cells were cultured to ∼90% confluency in 15-cm plates, pre-treated with staurosporine for 3 hours. For HEK293T cells, cells grew to about 70% confluency, then treated with tunicamycin for 24 hours. After aspirating the growth medium, the cells were incubated in serum-free medium containing FFF probes (20 µM for in-gel labeling, 100 µM for proteomics, site of labeling and gel-based validations) for 30 minutes at 37 °C in a 5% CO_2_ atmosphere. Following treatment, the cells were exposed to UV light (365 nm) for 20 minutes, harvested, washed with cold DPBS, and transferred to tubes. The cell suspensions were then centrifuged (400*g*, 5 min), washed with cold DPBS, and the resulting pellets were stored at −80 °C until further processing.

### Gel-based fluorescence analysis

Gel-based analysis of probe crosslinked proteins was performed as previously described^84^. Briefly, 6 μl of a freshly prepared click reaction mixture was added to each 50 μl protein sample (2 mg ml^−1^). The click mix included 3 μl of 1.7 mM TBTA tris[(1-benzyl-1H-1,2,3-triazol-4-yl)methyl]amine (TBTA) in 4:1 *t*-BuOH:DMSO, 1 μl of 50 mM CuSO_4_ in water, 1 μl of 1.25 mM tetramethylrhodamine (TAMRA) azide in DMSO, and 1 μl of freshly prepared 50 mM tris(2-carboxyethyl)phosphine (TCEP) in DPBS. Samples were mixed with the click mix and incubated at room temperature for 1 hour. Reactions were then quenched with 17 μl of 4× SDS sample buffer. Proteins (15 μg per lane) were resolved by SDS-PAGE using in-house prepared 10% acrylamide gels and visualized via in-gel fluorescence with a Bio-Rad ChemiDoc Imaging System. Images were processed using Image Lab software.

### Preparation of samples for proteomic analysis

For unenriched proteomic samples, cell pellets were resuspended in 100 μl ice-cold DPBS containing 1× protease inhibitor cocktail (Thermo Fisher Scientific, 78441) and lysed by probe sonication (Branson Sonifier; 15 ms on, 40 ms off, 15% amplitude, 7 pulses). Protein concentrations were measured using the BCA assay (Thermo Fisher Scientific). 100-200 μg of protein was denatured in 8 M urea, reduced with 5 mM DTT, and alkylated with 55 mM IAA. Proteins were precipitated with 4:1 methanol/chloroform, washed, and resuspended in 100 mM TEAB. Sequential digestion was performed using LysC (New England Biolabs, P8109S) for 2 h followed by trypsin (Promega, V5111) overnight at 37 °C in the presence of 1 mM CaCl₂. Peptides were quantified using the Pierce Quantitative Fluorometric Peptide Assay (Thermo Fisher Scientific), labeled with TMT10plex reagents (Thermo Fisher Scientific) in 30% acetonitrile, Labeling was quenched with 6 μl of 5% hydroxylamine (v/v), followed by a 15-minute incubation. Samples were acidified with 8 μl of formic acid, dried, resuspended in 0.1% TFA. Fractionation was carried out using the Pierce High pH Reversed-Phase Fractionation Kit (Thermo Fisher Scientific, 84868). MDA-MB-231 peptides were divided into 18 fractions and pooled into 9 sets; HEK293T peptides were divided into 28 fractions and pooled into 14 sets. All fractions were dried and stored at −80 °C until LC-MS analysis.

For phosphoproteomic samples, MDA-MB-231 cells were treated with 0.25 μM staurosporine for 3 h prior to harvest. Cell pellets were lysed in 1 ml ice-cold DPBS containing protease and phosphatase inhibitors (Thermo Fisher Scientific, 78441) and sonicated. Proteins were denatured with 4 M urea, reduced (DTT, 5 mM), alkylated (IAA, 55 mM) and precipitated. Pellets were resuspended in 1 M urea in 50 mM HEPES (pH 8.5), digested with LysC and then trypsin, and desalted using Sep-Pak C18 cartridges. Approximately 2 mg of peptides per sample were enriched using TiO₂ beads (GL Sciences, #5010-21315) or Fe-NTA beads (Thermo Scientific, A32992), following manufacturer protocols. For TiO₂ enrichment, peptides were resuspended in 2 M lactic acid, 50% acetonitrile and incubated with beads at a 1:4 peptide-to-bead ratio for 1 hour at room temperature with vortexing. Beads were washed three times with binding buffer and three times with wash buffer (50% acetonitrile, 0.1% TFA). Phosphopeptides were eluted with 50 mM potassium phosphate (pH 10) and desalted using Sep-Pak C18 cartridges. The eluates from the TiO_2_ or Fe-NTA enrichment were dried in a vacuum centrifuge, then labeled with TMT10plex reagents (Thermo Fisher Scientific) for 1 hour at room temperature, quenched, acidified and dried. Then peptides were resuspended in 0.1% TFA, pooled, and fractionated using the Pierce High pH Reversed-Phase Fractionation Kit into 18 fractions (7.5-95% acetonitrile), which were pooled into 9 final sets, dried, and stored at −80 °C.

Chemoproteomic samples were prepared in biological duplicates for each condition using previously described procedures^85^. Briefly, cell pellets were resuspended in 500 µl of cold DPBS and lysed via sonication. Protein concentrations were normalized to 2 mg ml^−1^ in 500 µl of ice-cold DPBS, using a DC Protein Assay (Bio-Rad), in 15-ml falcon tubes. The “click-chemistry cocktail” was then added to each lysate, consisting of TBTA (final concentration: 100 µM), TCEP (final concentration: 1 mM), biotin-azide (final concentration: 100 µM), and CuSO_4_ (final concentration: 1 mM). Samples were incubated at room temperature for 1 hour. Ice-cold 4:1 MeOH/CHCl_3_ (2.5 ml) and ice-cold DPBS (1 ml) were added to each sample to quench the click reaction, followed by centrifugation (3,200g, 4 °C, 10 min) to precipitate proteins. The precipitates were washed with 4:1 MeOH/CHCl_3_ and resuspended in 500 µl of 6 M urea in DPBS, along with 10 µl of 10% SDS. Protein reduction was performed by adding 50 µl of a freshly prepared 1:1 solution of TCEP (25 µl, 200 mM in DPBS) and K_2_CO_3_ (25 µl, 600 mM in DPBS), and incubate at 37 °C for 30 minutes. Following reduction, proteins were alkylated by adding 70 µl of freshly prepared iodoacetamide (400 mM in DPBS) and incubated for 30 minutes at room temperature in dark. After alkylation, 130 µl of 10% SDS was added to the samples, followed by dilution with 5.5 ml of DPBS. A 100 µl aliquot of 50% streptavidin agarose slurry (Thermo Fisher Scientific, 20349) was added to each tube, and samples were rotated for approximately 1.5 hours at room temperature to enrich probe-labeled proteins. The beads were then centrifuged down to tube bottom (750*g*, 2 min, 4 °C) and washed once with 0.2% SDS in DPBS (5 min), twice with DPBS, once with water, and transferred to LoBind microcentrifuge tubes with 100 mM TEAB (pH 8.5).

Bound proteins were digested overnight with trypsin (Promega, V5111) in digestion buffer (100 ml, 100 mM TEAB (pH 8.5), 100 µM CaCl_2_) at 37 °C. The resulting peptides were transferred to new tubes for each condition and labeled with the respective tandem mass tags (TMT; Thermo Fisher Scientific) 10plex or 16plex reagents (8 µl, 20 µg µl^−1^) for 1 h, quenched, acidified and dried through SpeedVac vacuum concentrator. The dried peptides were redissolved in 0.1% TFA in H_2_O, combined, and proceeded to fractionated into 12 fractions by Pierce high pH Reversed-Phase Fractionation Kit. The fractions were pooled pairwise into 6 final fractions, dried, and stored at - 80 °C until mass spectrometry analysis.

The preparation of site-of-labeling (SoL) samples followed our previously reported protocol^24^. Briefly, cell pellets were resuspended in cold DPBS (1000 µl) and lysed via sonication. 3 mg of the lysates were taken and diluted into 1ml of DPBS in 15 ml falcon tubes. Each sample was then added with TBTA (final concentration: 100 µM), TCEP (final concentration: 1 mM), cleavable biotin azide tag (light L-Valine tag, final concentration: 100 µM), and CuSO_4_ (final concentration: 1 mM) for 1 hour. Cold methanol (4 ml) was added, and the protein was allowed to precipitate overnight. The precipitated proteins were pelleted by centrifugation (4,000g, 10 min), and the supernatant was discarded. The pellets were washed with 4:1 (*v*/*v*) MeOH/CHCl_3_, resuspended by sonication, and centrifuged again (4,000g, 10 min). The pellets were then resuspended in freshly prepared 6 M urea solution (1000 µl, in DPBS) and a solution of SDS (100 µl, 10 % *w*/*v*), followed by reduction and alkylation steps as described above. The samples were diluted with DPBS (11 ml), streptavidin-agarose slurry (200 µl, 50 %, Pierce) was added, and the tubes were rotated for 1.5 hours at room temperature. The beads were pelleted by centrifugation (500g, 5 min) and washed with 0.2 % SDS in DPBS (1×), DPBS (2×), water (1×), and 100 mM TEAB (2 × 5 ml, pH 8.5 in H_2_O). The beads were then transferred to 1.5 ml LoBind microcentrifuge tubes, added with sequencing-grade modified trypsin (4 µg, 100 mM TEAB, pH 8.5, 100 µM CaCl_2_) was added and incubated overnight at 37 °C. The beads were further washed with SDS in DPBS once (0.2 % *w*/*v*), NaCl in DPBS (2×, 150 mM), and water (2×) by centrifugation (400g, 5 min). Elution was performed with a buffer of formic acid/water (3 % v/v, 200 µl) for 1 hour with gentle shaking. The supernatant was collected, and the beads were re-eluted with the same buffer, followed by an additional elution with acetonitrile/water (50 % v/v, with 3 % formic acid, 200 µl). The supernatants were combined and dried using vacuum centrifugation. The resulting residue was reconstituted in TEAB (100 mM) and labeled with TMT reagents (Thermo Fisher) across replicates as per the manufacturer’s instructions. TMT-labeled samples were pooled, dried, fractionated into 12 fractions by Pierce high pH Reversed-Phase Fractionation Kit, combined into 6 final fractions, dried, and stored at −80 °C until mass spectrometry analysis.

### LC-MS analysis of proteomic samples

LC-MS analysis was conducted following our previously published protocol^24^. For phosphor, unenriched and enriched proteomics, samples were resuspended in MS sample buffer (20 µl, 5% acetonitrile, 0.1 % formic acid in water) prior to LC-MS analysis, following published methoeds^85^.

In short, 3 µl of each sample was loaded onto an Acclaim PepMap 100 precolumn (75 µm × 2 mm) and separated on a PepMap RSLC analytical column (2 µm, 100 Å, 75 µm × 25 cm) using an UltiMate 3000 RSLCnano system (Thermo Fisher Scientific). Buffer A (0.1% formic acid in H_2_O) and buffer B (0.1% formic acid in acetonitrile) were used in a 220-minute gradient at a flow rate of 300 µl min^−1^. The gradient consisted of 2% buffer B for the first 10 min, a gradual increase to 30% buffer B over the next 192 min, 60% buffer B for 5 min, followed by a ramp to 95% buffer B in 1 min, held for 5 min, then back to 2% buffer B in 1 min, and re-equilibrated for 6 min.

Eluents were analyzed using a Thermo Fisher Scientific Orbitrap Fusion Lumos mass spectrometer with a 3-second cycle time and a nano-LC electrospray ionization voltage of 2,000 V. Data were acquired with Xcalibur software (v4.1.50). MS1 spectra were captured with a scan range of 375-1,500 m/z, a maximum injection time of 50 milliseconds, and dynamic exclusion enabled (1 repeat, exclusion duration of 20 seconds). The resolution was set to 120,000 with an AGC target of 1 × 10^6^. Peptides chosen for MS2 analysis were fragmented by collision-induced dissociation (CID) at 30% collision energy at 1.6 m/z isolation window. Fragment ions were recorded in the ion trap (AGC target 1.8 × 10^4^, maximum injection time of 120 ms). MS3 spectra were generated using high-energy collision-induced dissociation (HCD) mode with 65% collision energy, and synchronous precursor selection (SPS) was set as 10.

For SoL proteomics, samples were resuspended in 30 μl of MS sample buffer, 10 μl was injected onto the analytical column. The loading phase involved 2% buffer B (0.1% formic acid in acetonitrile) for 10 min, followed by elution with a gradient of 2-35% buffer B over 215 min, 35-55% over 10 min, 55–95% over 5 min, 95% buffer B for 2 min, then decreasing to 2% buffer B over 1 min. After holding at 2% for 2 min, the gradient was ramped to 95% buffer B in 1 min, held at 95% for 2 min, decreased again to 2% in 1 min, and held at 2% for an additional 11 min, totaling 260 minutes of run time.

Eluents were analyzed using a ThermoFisher Orbitrap Fusion Lumos mass spectrometer. MS1 scans were performed with a resolution of 120,000, a scan range of 375-2,000 m/z, and an RF lens setting of 60%, with a dynamic exclusion window of 30 seconds. MS2 scans utilized the orbitrap mass analyzer with a resolution of 15,000 and a first mass of 120. Peptide isolation and fragmentation were carried out with quadrupole isolation of 1.6 m/z, an AGC target of 5 × 10⁴, a maximum injection time of 110 ms, and a high-energy collision-induced dissociation (HCD) energy of 28%. For MS3, the orbitrap mass analyzer was set to a resolution of 50,000 with an AGC target of 1.5 × 10⁵, a maximum injection time of 120 ms and an HCD collision energy of 65% (SPS 10).

### Proteomic result analysis

Proteomic analysis was carried out using Proteome Discoverer (v3.0, Thermo Fisher Scientific) following previously established protocols^85^. For phosphor, unenriched and enriched proteomic results, peptide identification was conducted using the SEQUEST HT algorithm, precursor mass tolerance is 10 ppm and fragment mass tolerances is 0.6 Da, allowing for 2 missed cleavage sites. Variable modifications included oxidation (M, +15.994915), carbamidomethyl (C, +57.02146) and TMT-tag modifications (K and N-terminal, +229.1629 for 10plex, +304.207 for 16plex tags) were used as fixed modifications. The spectral data were searched against the Homo sapiens proteome database (Uniprot, 2018; 42,358 sequences), applying a 1% false discovery rate (FDR) with Percolator. MS3 peptide quantification was performed using a mass tolerance of 20 ppm.

For unenriched proteomic analysis, abundances in each channel were normalized to the average of the median signal intensity across all channels. For phorsphoproteomic analysis, no normalization was applied to peptide abundances. For enriched proteomic results, abundances in each channel were normalized to endogenously biotinylated proteins (PCCA, PC, MCCC1, or ACACA). For phospho-dependent experiments **P2** and **P6**, no normalization was applied to protein abundances.

Site of labeling analysis was conducted according to the previously published protocol^24^, using Proteome Discoverer (v3.0, Thermo Fisher Scientific) for data processing. Peptide sequences were identified by comparing experimental fragmentation patterns with proteome databases via the SEQUEST HT algorithm. The precursor mass tolerance was set to 10 ppm, fragment mass tolerance to 0.05 Da, and up to two missed cleavage sites were allowed. Static modifications included TMT at the N-terminus (+229.163), while variable modifications specified were carbamidomethyl (C, +57.02146), oxidation (M, +15.994915), TMT tag (K and N-terminal, +229.1629 for 10plex, +304.207 for 16plex tags), and FFF-tags (**P2**: +481.310; **P3**: +501.274; **P4**: +467.217; **P5**: +517.305; **P6**: +453.274) on all amino acids. The spectral data were searched against each probe’s enrichment database from the whole protein profiling experiment with a 0.01 false discovery rate (FDR) using the target-decoy approach. Further validation was performed using Percolator, applying the target-decoy strategy with a ‘Separate’ setting and a maximum delta Cn threshold of 0.1. TMT quantification was carried out at the MS3 level, with a tolerance of 20 ppm. Only peptides with an average signal-to-noise ratio (S/N) greater than 10 were retained for further analysis.

For MSFragger SoL searches, an open search workflow was used^86^. In brief, precursor mass tolerance was set to 0–520 Da, MS1 tolerance was set to 20 ppm. Full trypsin specificity was designated with a maximum of two missed cleavages. Peptide length was set from 4 to 50, peptide mass range set 500 to 5000. Oxidation (M, +15.9949), (C, +57.02146), TMT tag (K, +229.1629 for 10plex, +304.207 for 16plex tags) as variable modification, TMT tag (N-terminal, +229.1629 for 10plex, +304.207 for 16plex tags) as fixed modifications.

To ensure data reliability, proteins reported required at least two unique peptides. TMT ratios calculated in Proteome Discoverer were transformed using log2(x), and statistical significance of the ratios was assessed with two-tailed Student’s t-tests across two biological replicates (significance threshold set at *p* < 0.05). Detailed proteomics data are available in Supplementary Tables 1-14.

### Gel-based analysis of probe crosslinked targets in cells

Cells were treated as the “Photoaffinity probe cell treatments” section described. Probe labelled cell pellets were lysed in ice-cold DPBS supplemented and 1× Halt protease inhibitors cocktail (Thermo Fisher Scientific, 78438) and lysed by sonication. Protein concentrations were normalized and click reactions were performed as described in chemoproteomic part in “Preparation of samples for proteomic analysis” section. In brief, after the “click-chemistry” reaction, 2 ml of ice-cold MeOH was added to each sample to precipitate proteins and incubated overnight in −20 °C. The samples were then centrifuged (3,200*g,* 4 °C, 10 min) and washed with 4:1 MeOH/CHCl_3_, resuspended by sonication and centrifuged as above. Pellets were resuspended in freshly prepared urea solution (500 µl DPBS, 6 M) with SDS (100 µl of 10% *w/v*). After incubating the samples for 30 minutes at 37 °C to fully dissolve the protein pellet, samples were diluted with DPBS (5.5 ml), added with a streptavidin-agarose slurry (100 µl, 50%, Pierce) and rotated for 1.5 hour at room temperature. The beads were pelleted by centrifugation (500*g*, 5 min, 4 °C) and washed once with DPBS with 0.2% SDS, three times with DPBS, then boiled with 2× SDS sample buffer for 15 minutes. Boiled samples and saved input samples were resolved by SDS-PAGE and followed by immunoblotting analysis.

### Knowledge-based and structure-based enrichment analyses

Protein enrichment analysis was conducted and modified following an established protein enrichment clustering workflow^26^. In brief, enriched proteins were analyzed and clustered based on protein classifications (PANTHER database 19.0 and Uniprot “Family and domain databases”)^87^. Significant and emerging significant hits were identified based on ligandability changes. Proteins were visualized in clusters reflecting functional and classification similarities. Downstream analyses included GO term and IDG category enrichment, using PANTHER family annotations and Illuminating the Druggable Genome database annotations, respectively. Clustering and plotting were performed using Python scripts adapted from prior work^26^.

Protein information for the labeled peptides was used to extract additional information from Uniprot, such as structural features (ACT_SITE, BINDING, METAL, DNA_BIND and NP_BIND), presence in DrugBank, gene ontology (GO) keywords and PDB and AF structure identities. Protein structures were batch downloaded from PDB or AlphaFold using command line prompts in terminal. Proteomic, pocket, site and structure data were processed together using R scripts adapted from previously published work^24^.

### Immunostaining

For NPC2 localization, HEK293T cells were seeded on 0.02% poly-L-lysine (Sigma-Aldrich, P1274) coated 8-well plates (ibidi, #80827) at 37 °C overnight. Cells were treated with tunicamycin for another 24h, fixed in Bouin’s solution (Sigma) for 10 min and permeabilized with ice-cold methanol at −20 °C for 10 min. After fixation/permeabilization, cells were washed for 5 min with PBS and then blocked with 3% BSA/PBST (PBS with 0.05% Triton X-100) for 10 min. Primary antibodies were diluted with blocking buffer (Rabbit anti-NPC2 1:100, Mouse anti-LAMP1 1:500) and incubated with cells for 1 h. Follow three washes with PBST, cells were incubated with Alexa Fluor-labeled secondary antibodies (1:500) for an additional hour. Cells were then washed three times with PBST, stained with NucBlue Probe (Thermo Fisher Scientific, R37606) for nuclear imaging. Confocal images were carried out on a Zeiss LSM 780 microscope using a 100× objective (NA = 1.40), and 1 Airy unit was set as a pinhole for each channel. Imaris, ImageJ and ZEN (Zeiss) software were used for image processing and analysis.

### Thermostability experiments

MDA-MB-231 cells were digested with 0.25% trypsin, and equal amounts of cells were incubated with DMSO or BI-2865 with or without staurosporine/SHP-099/dasatinib in fresh DMEM media for 30 min at 37 °C. Cells were then collected and washed once with binding buffer (cold DPBS containing DMSO or the indicated concentration of inhibitors and 1× Halt protease inhibitor cocktail). Cells were then resuspended in binding buffer, counted and aliquoted to a concentration of 300,000 cells per 50 μl before being transferred to PCR tubes for heat treatment. PCR tubes were heated at the indicated temperature gradient on a thermal cycler (Bio-Rad) for 3 min, followed by incubation at 25 °C for 3 min. After three snap freeze–thaw cycles with liquid nitrogen, lysates were transferred to 1.5-ml Eppendorf tubes and centrifuged at 100,000g for 20 min at 4 °C to separate destabilized proteins from proteins remaining in solution. The resulting supernatant was transferred to new Eppendorf tubes, mixed with 4× SDS sample buffer, boiled for 5 min and processed for immunoblotting as described below.

### RAS-binding domain (RBD) pulldown assay

RBD experiments were performed using the Active Ras Pull-Down and Detection Kit (Thermo Scientific, 16117) following the manufacturer’s instructions. Briefly, after compound treatment for the specified duration, cells were washed with ice-cold PBS, harvested, snap-frozen in liquid nitrogen, and stored at −80 °C until use. Cells were lysed in lysis buffer, vortexed, and incubated on ice for 15 min. The lysates were then centrifuged at 14,000g for 15 min at 4 °C, and supernatants were collected. Cell lysates were incubated with Raf-GST and GST beads at 4 °C for 1.5 h with rotation, washed three times with lysis buffer, and eluted by incubating in 2× sample buffer for 15 min. For HA-KRAS^G13D^, HA-KRAS^G13D/Y32F^ or KRAS^G13D/Y64F^, the cell lysates were immunoprecipitated using anti-HA magnetic beads overnight at 4 °C. The beads were washed three times with lysis buffer and eluted by incubating in 2× sample buffer for 30 min. All samples were resolved by SDS-PAGE and processed for immunoblotting as described below.

### Immunoprecipitation and immunoblotting

For immunoblotting, cells were treated with indicated compounds for specified time, then washed with ice-cold PBS and lysed in NP-40 buffer (10 mM Tris-HCl, 150 mM NaCl, 1 mM EDTA, 1% (w/v) NP-40) supplied with protease and phosphatase inhibitors. After centrifuging at 14,000g for 10 min at 4 °C to remove cell debris, supernatants were taken, and protein concentrations were determined by a Bio-Rad DC Protein Assay (5000112). Equal amounts of protein lysates were mixed with the 4× SDS sample buffer and resolved by SDS-PAGE.

For LAMTOR2-FLAG immunoprecipitation, cells were resuspended in E1C buffer (50 mM HEPES, 250 mM NaCl, 5 mM EDTA, 0.3% (w/v) CHAPS supplied with protease inhibitor and sonicated for three times. Lysates were cleared by centrifugation and normalized to 1.5-2.0 mg ml^−1^. Then 1 µg of anti-FLAG antibody was added to supernatants and rotated overnight at 4 °C, and incubated with 20 μl protein A/G agarose beads (Medchemexpress) for an additional hour on the next day. The beads were washed three times with the E1C buffer and eluted by incubating in 2× sample buffer at 95 °C for 10 min and resolved by SDS-PAGE.

For HA-KRAS immunoprecipitation, after compound treatment for the designated time, cells were washed twice with ice-cold PBS and lysed in NP-40 buffer supplied with protease and phosphatase inhibitors. Lysates were then cleared by centrifugation, normalized, and incubated with 20 µl of anti-HA magnetic beads at 4 °C for 3 hours or overnight with rotation. The beads were washed three times with lysis buffer and eluted with 2× sample buffer.

Resolved proteins were transferred to PVDF membranes using the Bio-Rad Trans-Blot Turbo Transfer system. Membranes were blocked with 5% non-fat milk or 3% BSA in TBST, incubated overnight with primary antibodies. Blots were washed three times with TBST and incubated with secondary antibodies for 1 hour. Membrane visualization was performed using the Bio-Rad ChemiDoc Imaging System, and band density was quantified with Bio-Rad Image Lab software (v.6.1.0).

### Modeling and docking

Three-dimensional coordinates of the probes were generated from SMILES using RDKit’s ETKDG v.3 followed by UFF energy minimization. Target protein structures were obtained from the Protein Data Bank (PDB) when available; otherwise, AlphaFold2-predicted models were used. In cases where multiple PDB structures existed, the most suitable was selected based on the presence of the SoL residue coordinates, highest sequence coverage, and best resolution, with a preference for X-ray crystallography over cryo-EM. In some specific cases, the structures were manually selected. These cases include: KRAS^G13D^ and KRAS^G12C^ (PDB: 8AZV and 8B00, respectively), and EphA2 (PDB: 5EK7). Hydrogen atoms were added using Reduce, and both ligands and receptors were prepared according to the standard AutoDock protocol^88^. Druggable pockets were identified using Fpocket, and a docking grid box was defined for each, encompassing the pocket region with a minimum side length of 24 Å. All probes were docked into each predicted pocket using AutoDock-GPU^89^, employing default parameters for the Lamarckian Genetic Algorithm and ADADELTA local search to generate 50 poses per ligand. Pockets were ranked based on the best binding energy and the proximity (<6 Å) of the diazirine moiety to any proteomics-identified adducted residue was used as a criterion for successful labeling. Pocket prediction success was defined as the percentage of docking events meeting these criteria across the entire probe library, while site prediction success considered only probes experimentally observed in proteomics. Molecular visualization of docking results was performed using ChimeraX 1.8.

### KRAS molecular dynamics simulations

Structural models of GDP-bound KRAS were generated via homology modeling (SwissModel) to ensure consistency with K-Ras4A (UniProt-ID: P01116). Crystallographic water molecules were retained, and protonation states were assigned at pH 7.4.

Molecular Dynamics (MD) simulations were performed in OpenMM 8.1.2 using Amber ff14SB for the protein and the recently published phosaa14SB for phosphotyrosine. Ligands were parameterized via Espaloma 0.3.2 and modified 12-6-4 Lennard Jones parameters were used for divalent metal ions. Systems were solvated with TIP3P-FB water molecules and 0.15 M NaCl. A Langevin Middle integrator was used with a 4 fs time step combining HMR and SHAKE algorithms. Harmonic positional restraints were applied on all heavy atoms and the systems were gradually heated to 300 K in the NVT ensemble. Subsequently, a Monte Carlo barostat was introduced, and 15 steps of NPT equilibration were performed, progressively releasing the restraints. Finally, 50 ns of unrestrained MD simulations were conducted to ensure proper equilibration.

The experimental structure of KRAS^G13D^ bound to the BI-2865 compound (PDB: 8B00) was simulated for 100 ns, and the average of the last 50 ns was used as a reference. The root-mean-square-deviation (RMSD) to the BI-2865 binding competent conformation was calculated over the Cα atoms of residues within 4 Å of the ligand.

### Enhanced sampling protocol

An ad hoc computational sampling pipeline was implemented to explore the conformational landscape of the systems efficiently. Initially, well-tempered metadynamics simulations (WTMetaD) were conducted, biasing the RMSD of Cα atoms relative to the initial conformation. Gaussian potentials of height 0.3 kcal/mol and width 0.01 nm were deposited every 2 ps with a bias factor of 10. Three simulations of 300 ns were run in parallel, sharing the bias history (i.e., multiple-walker scheme). Then, all Cα pairwise distances (13,041 features) were calculated from the trajectories, and MoSAIC analysis was used to identify clusters of highly correlated motions. Features from top-3 bigger clusters were subjected to Principal Component Analysis, retaining the components that explained 95% of the variance. Regular space clustering in the reduced space was conducted to achieve uniform sampling, and 30 representative conformations were extracted for each system. Each state was equilibrated, followed by 200 ns of unbiased MD.

### Metadynamics free energy surfaces

WTMetaD simulations were performed with PLUMED 2.9 using G12-T35 and G35-G60 minimum distances as collective variables. Gaussians (0.5 Kcal mol height^−1^, 0.01 nm width) were deposited every 1 ps with a bias factor of 12. Four independent replicas of 100 ns were simulated per system and combined using the Mean Force Integration method to obtain the free energy surfaces.

### Statistical analysis

All fluorescent gel scanning and western blots were performed with a minimum of two biological replicates. Compound effects at different doses were normalized to vehicle treated cells and fitted with Sigmoidal 4PL curve. All statistical analysis was performed using GraphPad Prism (v9.3.0 or v10.4.0) or Microsoft Excel (v.2122). Proteomic data were statistically evaluated using two-tailed Student’s *t*-tests performed across a minimum of two biological replicates, with significance defined as *p* < 0.05.

### Data Availability

The data that support the findings in this work are available within the paper and Supplementary Information (Supplementary text and Supplementary Tables 1-14). Uncropped, full western blot images and fluorescence gels are provided in Supplementary Figs. 2-12. All raw proteomics data files have been deposited to the PRIDE repository and available under the accession PXD056973 (unenriched and phosphor-proteomics) PXD056976 (chemoproteomics) and PXD056980 (site of labeling) (https://www.ebi.ac.uk/pride/archive/projects/ PXD056973; https://www.ebi.ac.uk/pride/archive/projects/ PXD056976; https://www.ebi.ac.uk/pride/archive/projects/ PXD056980).

## Acknowledgments

C.G.P.’s research in this area is supported by National Institute of Allergy and Infectious Diseases R01AI156268 and R01AI182439 as well as 2023-332369 from the Chan Zuckerberg Initiative DAF, an advised fund of Silicon Valley Community Foundation. Q.W. was supported by Hou Wang Fellowship in the Skaggs Graduate School. M.L.H. acknowledges support from the National Institute of General Medical Sciences under R35GM142462. The authors thank Benjamin Cravatt for useful discussions and input.

## Author contributions

C.G.P., W.L., and Q.W. conceived the study, designed experiments and interpreted results. W.L. and Q.W. performed the proteomics experiments, analyzed MS data, performed validation experiments with input from A.D. and M.L.H. S.F. and P.G. performed molecular docking analysis. S.F., M.L. and M.H. performed molecular dynamics analysis. T.-Y.C, C.G. and J.C. assisted with validation experiments. J.M.W. assisted with proteomics experiments. A.M.J. synthesized compounds. C.G.P., W.L., and Q.W. wrote the manuscript with input from all authors.

## Materials & Correspondence

should be addressed to Christopher G. Parker.

## Data and materials availability

PRIDE repository and available under the accession PXD056973 (unenriched and phosphor-proteomics) PXD056976 (chemoproteomics) and PXD056980 (site of labeling). All other data needed to evaluate the conclusions in the paper are present in the paper or the Supplementary Materials.

---

**Extended Data Fig.1.**
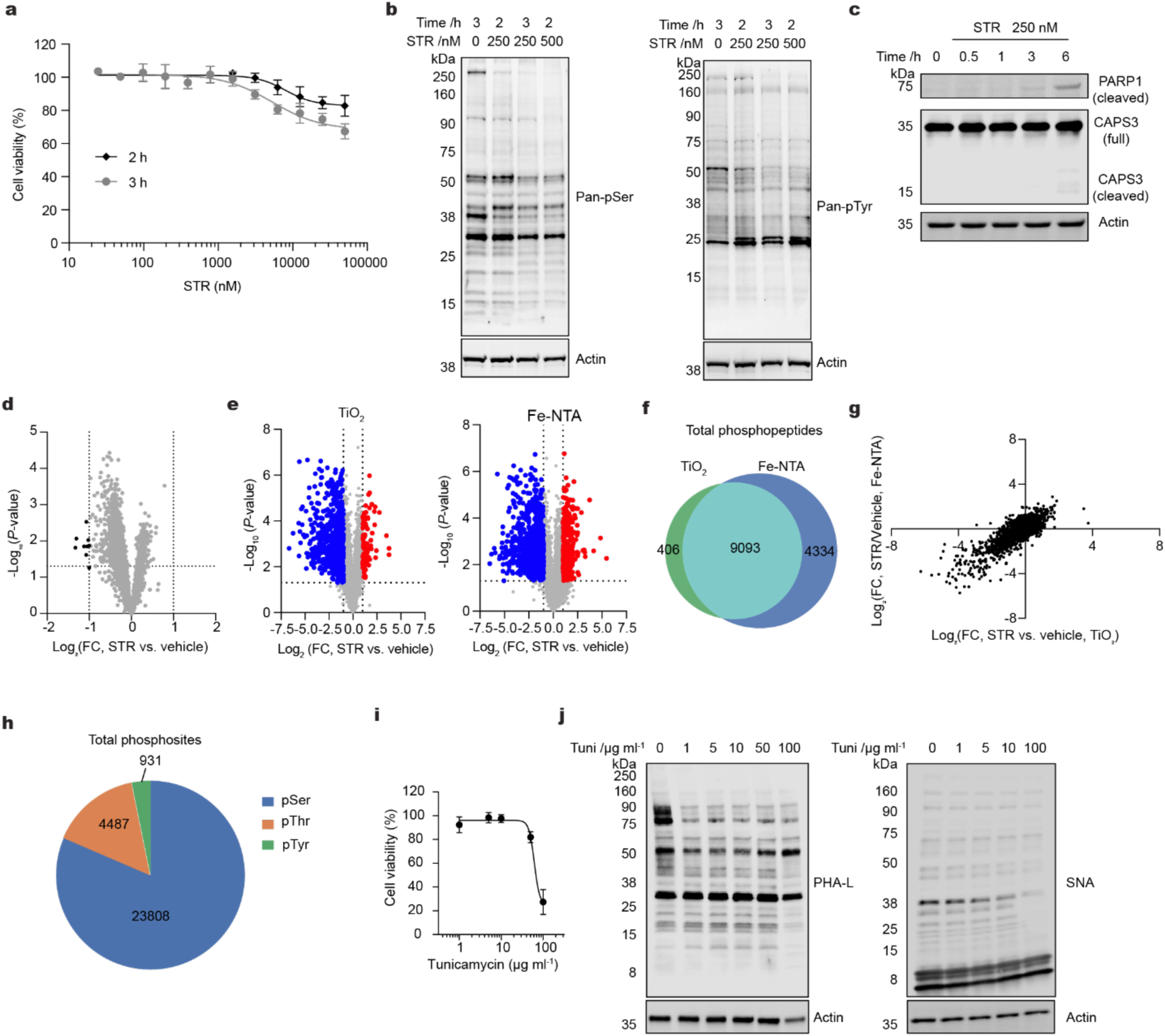
Characterization of PTM-dependent model systems. **a**, Cell viability of MDA-MB-231 cells treated with staurosporine (STR) at indicated concentrations and treatment times. **b**, Immunoblot analysis of global serine (left) and tyrosine (right) phosphorylation levels in MDA-MB-231 cells across STR concentrations and treatment durations. **c**, Apoptosis evaluation in MDA-MB-231 cells treated with 250 nM STR, as indicated by changes in the abundance of cleaved PARP1 and caspase-3 at indicated time points. **d**, Volcano plots of protein abundances comparing PTM-dependent targets displaying abundance changes after STR treatment. Vertical dashed lines represent a 2-fold change (FC) threshold, while horizontal dashed lines denote a *p*-value threshold of 0.05. Black dots highlight proteins that meet both significance thresholds. **e**, Volcano plots of phosphoproteomics experiments using different enrichment methods. Left, TiO_2_ enrichment; right, Fe-NTA enrichment. The vertical dashed line corresponds to a 2-fold change (FC) and the horizontal line corresponds to a p-value of 0.05. Colored dots correspond to phospho-peptides passing both specified thresholds, gained (red) and loss (blue). **f**, Venn diagram comparing phospho-peptides enriched by two methods. **g**, Correlation of phospho-peptide quantitation by two enrichment methods. **h**, Pie chart of enriched phospho-sites distribution, pSer (blue), pThr (orange) and pTyr (green). See data in Table S9-10. **i**, Assessment of HEK293T cell viability following treatment with tunicamycin (Tuni) at indicated concentrations for 24 h. **j**, Immunoblot analysis of global N-linked glycosylation levels in HEK293T cells treated with tunicamycin for 24 h at indicated concentrations, using *Phaseolus vulgaris* leucoagglutinin (PHA-L) and *Sambucus nigra* agglutinin (SNA) lectins. Proteomic data and cell viability data were obtained from three independent biological replicates and are presented as mean ± s.d. Gel images are representative of two independent experiments. Significance was determined using a two-tailed Student’s *t*-test. Associated datasets are provided in Supplementary Tables 5, 9-10.

**Extended Data Fig. 2.**
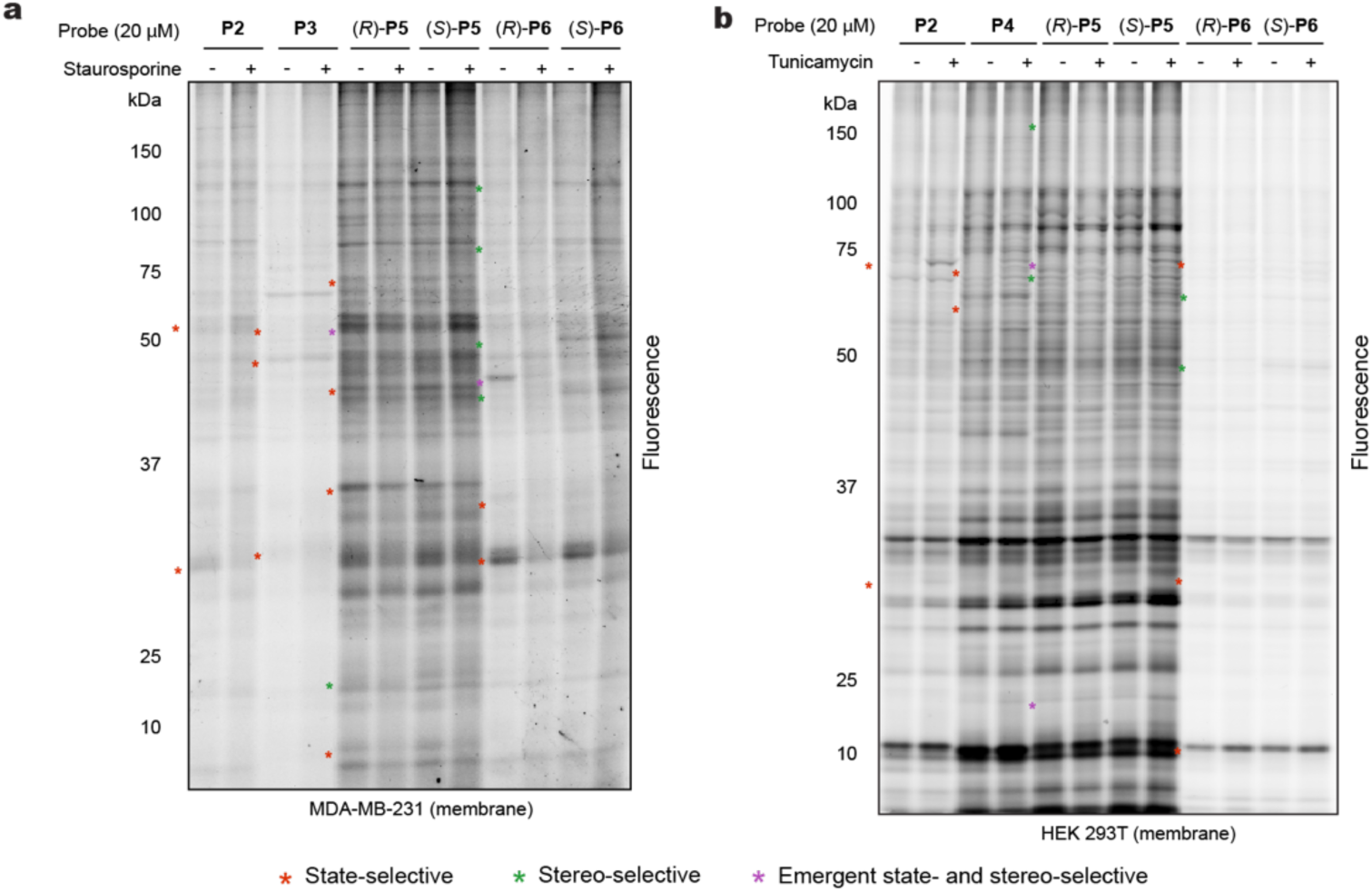
Gel profiling of probe-protein interactions (membrane fraction) **a,** Gel-based profiling of probe labeling under different phosphorylation states in MDA-MB-231 cells (membrane fraction). **b,** Gel-based profiling reveals N-linked glycosylation-dependent changes in HEK293T cells (membrane fraction). Red asterisks indicate representative state-selective probe–protein interactions, green asterisks denote stereoselective enantio-probe–protein interactions, and purple asterisks highlight interactions that are both state-selective and stereoselective. Gel images illustrate representative findings from two independent experiments.

**Extended Data Fig. 3.**
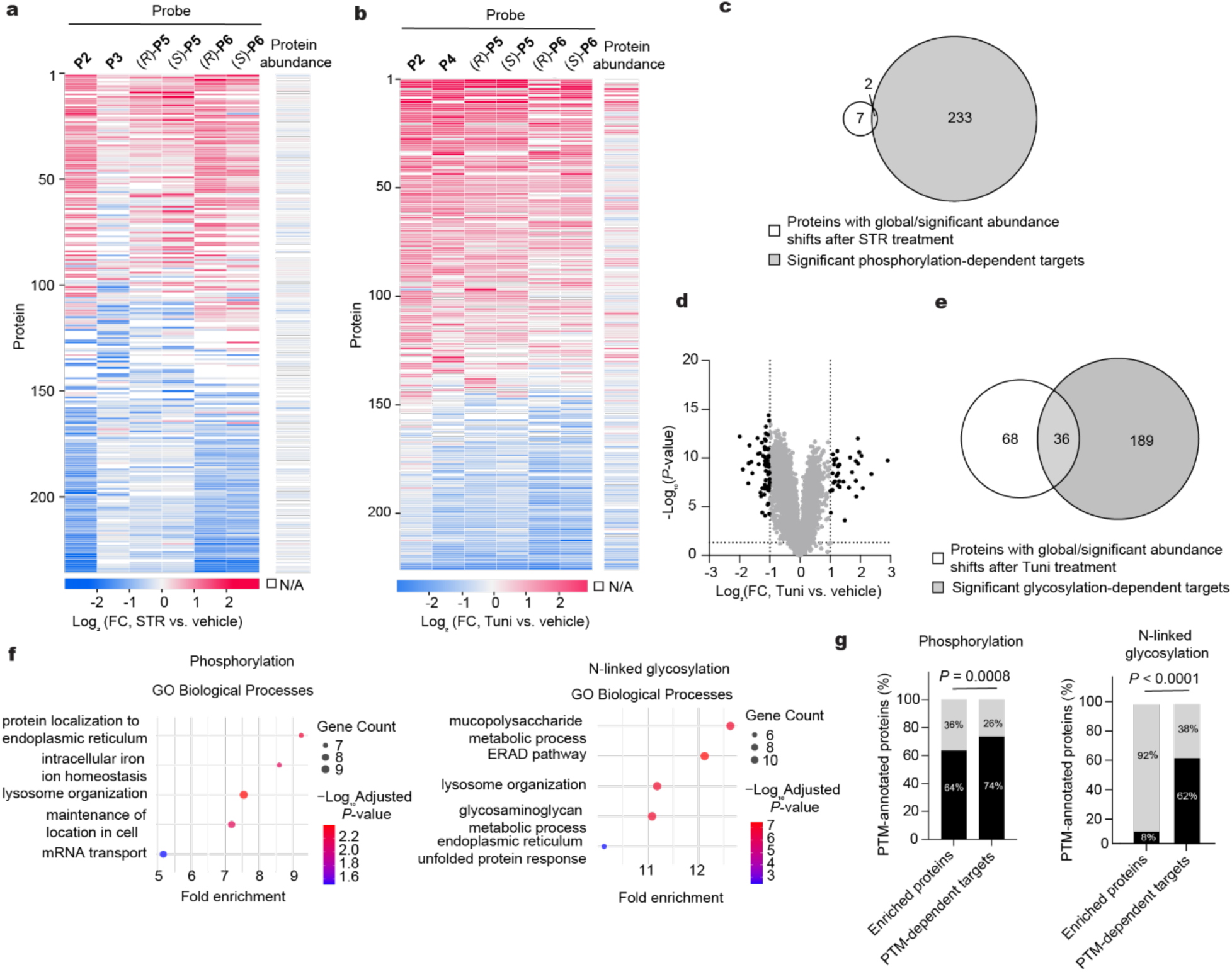
Proteomic profiling of PTM-dependent ligandability changes in cells. **a-b**, Heatmap of proteins displaying significant ligandability changes (FC >2, *p* < 0.05) after treatment with STR or Tuni. PTM-dependent targets are annotated individually by abundance level. **a**, phosphorylation; **b**, N-linked glycosylation. Observed proteins with increased (red) and decreased (blue) ligandability changes. Protein abundance changes before and after STR or Tuni treatment were detected by unenriched proteomics. N/A, not detected from the unenriched proteomics. **c**-**e**, Venn diagram (**c**, **e**) and volcano plots (**d**) of protein abundances and comparing PTM-dependent targets displaying abundance changes after STR (**c**) or Tuni (**d**-**e**) treatment. Vertical dashed lines represent a 2-fold change (FC) threshold, while horizontal dashed lines denote a *p*-value threshold of 0.05. Black dots highlight proteins that meet both significance thresholds. **f**, Gene Ontology (GO) enrichment analysis of proteins with PTM-dependent ligandability changes. Left, phosphorylation; right, N-linked glycosylation. **g,** Proportion of PTM-annotated proteins (UniProt) among enriched proteins and PTM-dependent targets, assessed by a Chi-square test. Black: proteins with annotated PTMs; shadow: proteins without PTM annotation. Proteomic data were obtained from two independent biological replicates and are presented as mean. Significance was determined using a two-tailed Student’s *t*-test. Associated datasets are provided in Supplementary Tables 1-8.

**Extended Data Fig. 4.**
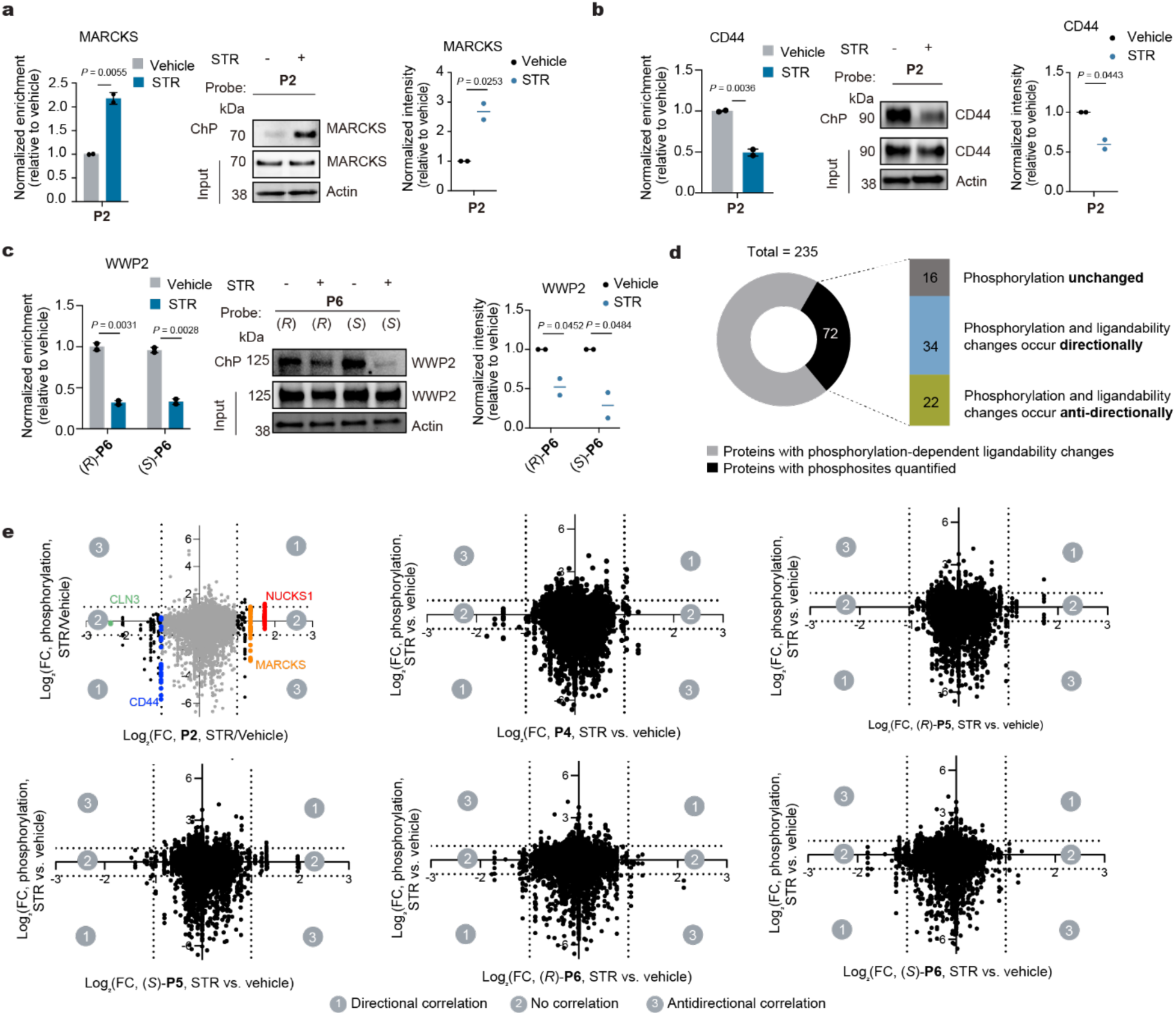
Analysis and validation of phosphorylation-dependent ligandability changes. **a-c**, Proteomic profiles and corresponding chemo-precipitation (ChP) validation for selected targets exhibiting PTM-sensitive probe binding: MARCKS (**a**), CD44 (**b**) and WWP2 (**c**). Left panels show quantitative chemoproteomic data; middle panels display ChP immunoblots; right panels show corresponding quantification across biological replicates. **d**, Distribution of phosphorylation-dependent targets with quantified phosphosites from phosphoproteomics experiments. **e**, Scatter plot depicting protein ligandability change versus phosphorylation change with indicated probes after STR treatment in MDA-MB-231 cells. Plots marked with three regions of ligandability changes: ligandability changes align with phosphorylation changes (region 1), ligandability changes occur without corresponding phosphorylation changes (region 2), and ligandability changes occur in the opposite direction of phosphorylation changes (region 3). The vertical and the horizontal dashed line represent a 2-fold change (FC) threshold. Proteomic data were obtained from two independent biological replicates and are presented as mean ± s.d. Immunoblots were quantified from two independent biological replicates and plotted as the mean. Gel images are representative of two independent experiments. Significance was determined using a two-tailed Student’s *t*-test. Associated dataset are provided in Supplementary Tables 1-4 and 9-10.

**Extended Data Fig. 5.**
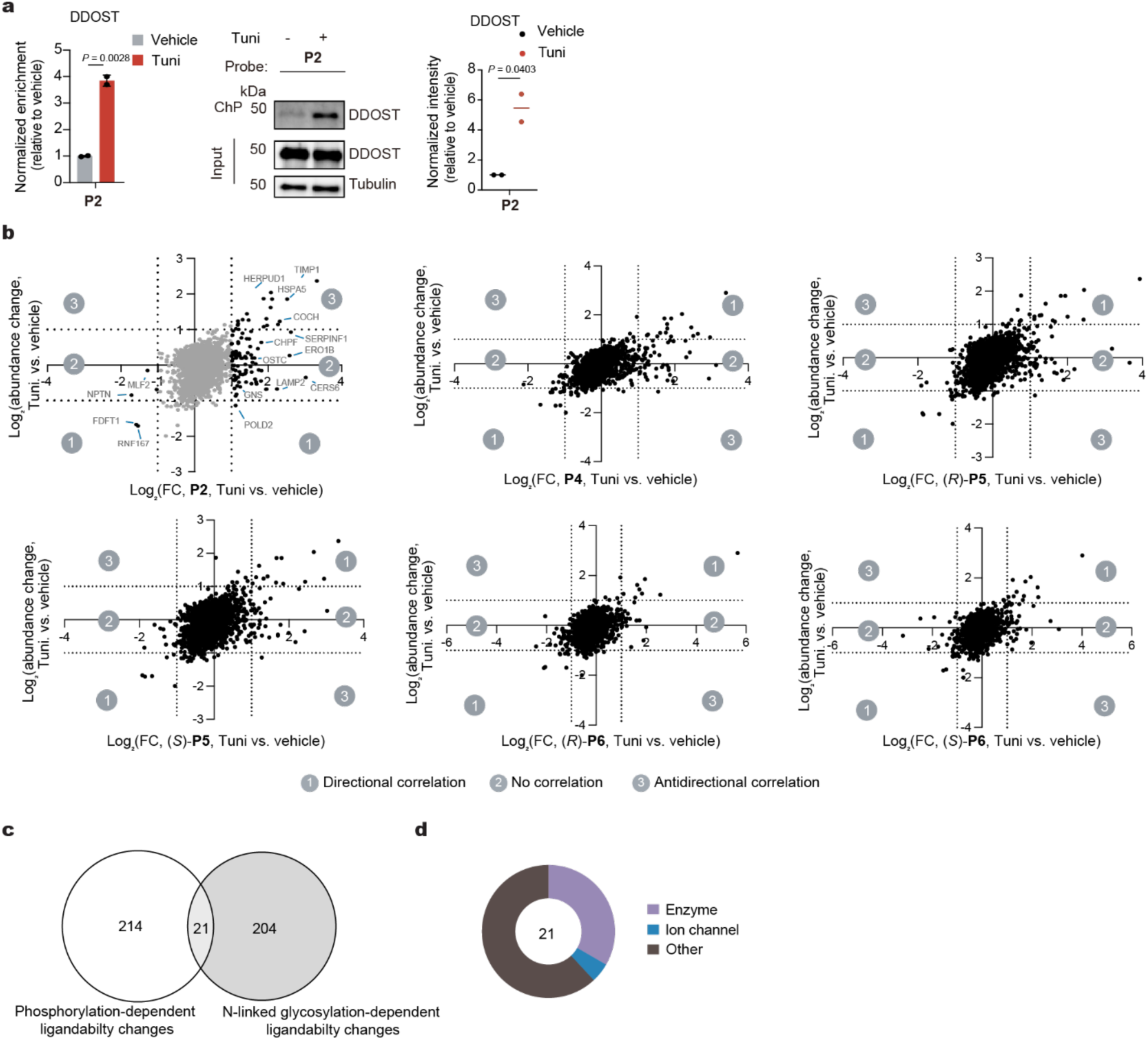
Analysis and validation of glycosylation-dependent ligandability changes. **a**, Proteomic profiles and corresponding chemo-precipitation (ChP) validation for DDOST. Left panel shows quantitative chemoproteomic data; middle panel shows ChP immunoblot results; right panel shows corresponding quantification across biological replicates. **b**, Scatter plot depicting protein ligandability change versus abundance change with indicated probes after Tuni treatment in HEK293T cells. Plots marked with three regions of ligandability changes: ligandability changes align with abundance changes (region 1), ligandability changes occur without corresponding abundance changes (region 2), and ligandability changes occur in the opposite direction of abundance changes (region 3). The vertical and the horizontal dashed line represent a 2-fold change (FC) threshold. **c**, Venn diagram comparing proteins with ligandability changes in both PTMs. **d**, IDG functional classes of proteins with ligandability changes in both PTMs. Proteomic data were obtained from two independent biological replicates and are presented as the mean. Associated datasets are provided in Supplementary Tables 1-4 and 6-8.

**Extended Data Fig. 6.**
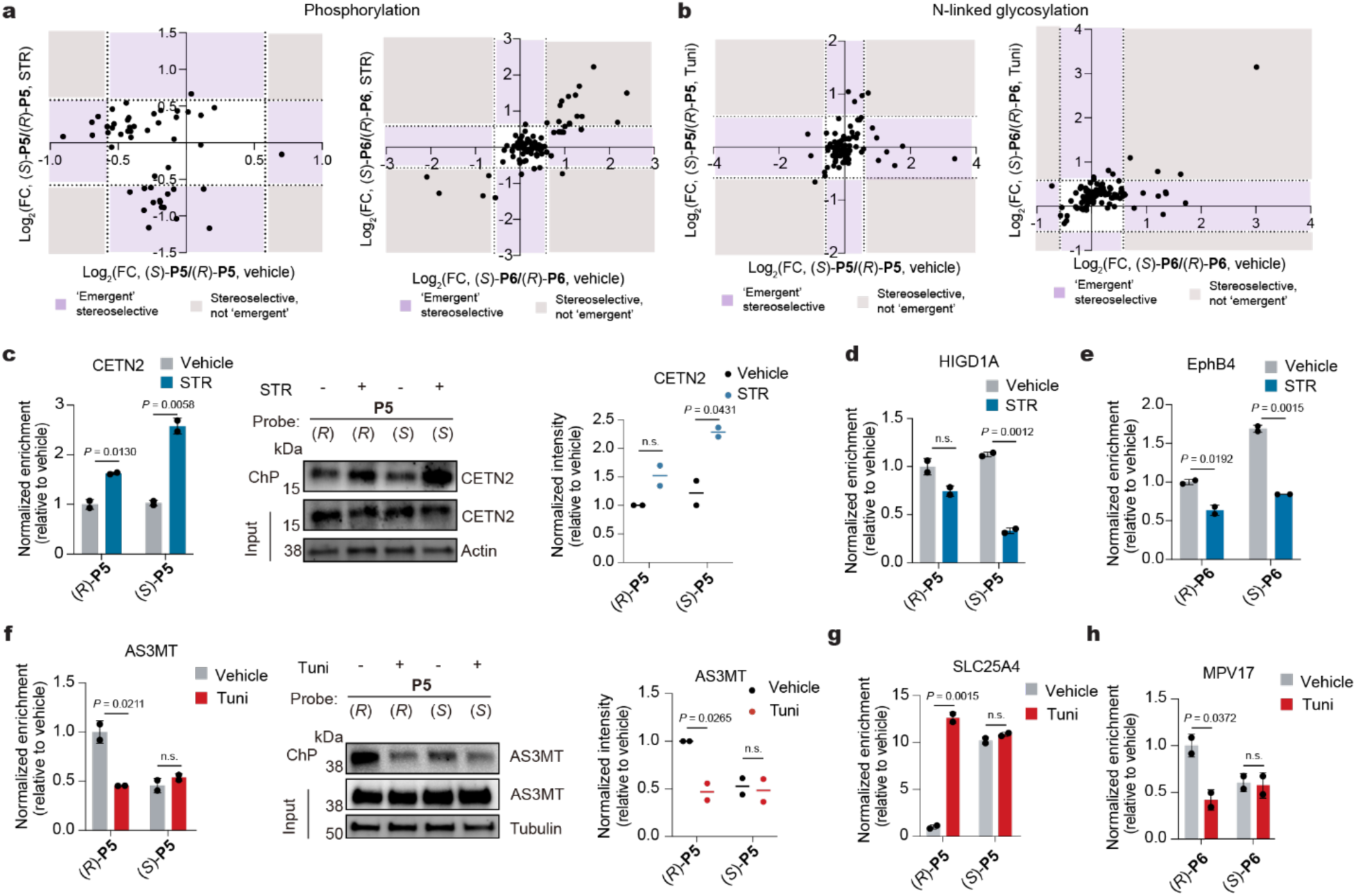
Discovery of emergent stereoselective PTM-dependent events. **a**-**b**, Scatter plots of significant PTM-dependent targets of enantioprobes to compare stereoselectivity between DMSO and STR or Tuni treated cells. The vertical and the horizontal dashed line represent a 1.5-fold change (FC) threshold. Proteins with emergent stereoselectivity (stereoselective engagement in one condition but not the other) are shown in pink area and ones with stereoselectivity in both states are in grey area. **a**, phosphorylation; **b**, N-linked glycosylation. **c**, **f**, Proteomic profiles (left), chemo-precipitation (ChP) validation (middle) and quantitation (right) of ligandability changes of CETN2 (**c**) and AS3MT (**f**). **d-e**, **g-h**, Proteomic profiles of indicated proteins. HIGD1A (**d**), EphB4 (**e**), SLC25A4 (**g**), MPV17 (**h**). Proteomic data were obtained from two independent biological replicates and are presented as mean ± s.d. Immunoblots were quantified from two independent biological replicates and plotted as the mean. Gel images are representative of two independent experiments. Significance was determined using a two-tailed Student’s *t*-test. n.s., not significant. Associated datasets are provided in Supplementary Tables 1-4 and 6-7.

**Extended Data Fig. 7.**
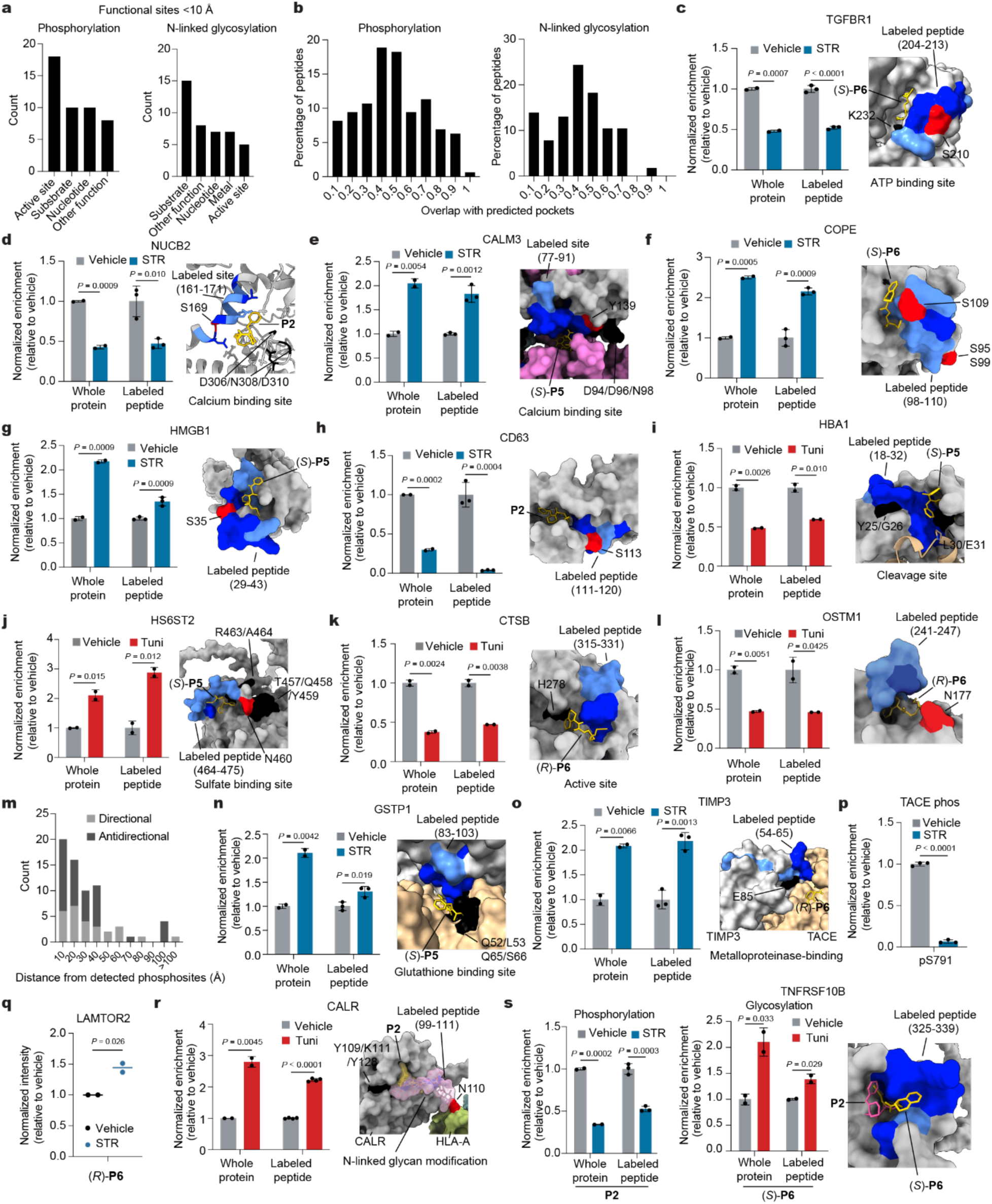
Features of pockets displaying PTM-dependent ligandability. **a**, Categorization of functional sites (Uniprot) proximal to probe-labeled peptides. **b**, Overlap of PTM-dependent probe-labeled peptides with predicted pockets (Fpocket algorithm). **c**-**l**, Proteomic profiles of indicated PTM-dependent targets and corresponding probe-labeled peptides (left) and docked structure of predicted pockets with probe labeled site highlighted (right). **c**, TGFBR1 with (*S*)-**P6** (PDB 3KCF); **d**, NUCB2 with **P2** (AF-P80303-F1); **e**, CALM3 with (*S*)-**P5** (PDB 7XNN); **f**, COPE with (*S*)-**P6** (PDB 6U3V); **g**, HMGB1 with (*S*)-**P5** (AF-P09429-F1). **h**, CD63 with **P2** (AF-P08962-F1). **i**, HBA1 with (*S*)-**P5** (PDB 1BZ1); **j**, HS6ST2 with (*S*)-**P5** (AF-Q96MM7-F1); **k**, CTSB with (*R*)-**P6** (PDB 3AI8); **l**, OSTM1 with (*R*)-**P6** (PDB 7JM7). **m**, Bins based on spatial distance of probe-labeled peptides to the detected quantified phosphosites. **n**-**o**, Proteomic profiles of indicated PTM-dependent targets and corresponding probe-labeled peptides (left) and docked structure of predicted pockets with probe labeled site highlighted (right). **n**, GSTP1 with (*S*)-**P5** (PDB 2A2R); **o**, TIMP3 with (*R*)-**P6** (PDB 3CKI). **p**, Quantified pS791 on TACE with STR treatment (250 nM, 3 h) in MDA-MB-231 cells. **q**, Quantitation of immunoblot validation of LAMTOR2 in Figure 4h. **r**, Proteomic profile of PTM-dependent binding of CALR with **P2** and corresponding probe-labeled peptides (left) and docked structure of predicted pockets (PDB 7QPD) with probe labeled site highlighted (right). **s**, Proteomic profile of PTM-dependent labeling of TNFRSF10B by indicated probes and corresponding probe labeled peptides (left, phosphorylation; middle, glycosylation) and docked structure of predicted probe binding (right). TNFRSF10B with **P2** (gold) in phosphorylation and (*R*)-**P6** (pink) in glycosylation (AF-Q8N8Q9-F1). For all structures, the primary peptide residues labeled by probes are colored dark blue and the remainder of each peptide is colored light blue. Reported PTM sites are colored red, functional sites are colored black, and probes are colored gold. Proteomic data were obtained from two or three independent biological replicates and are presented as mean ± s.d. Immunoblots were quantified from two independent biological replicates and plotted as the mean. Significance was determined using a two-tailed Student’s *t*-test. Associated datasets are provided in Supplementary Tables 1-4, 6-7 and 9-12.

**Extended Data Fig. 8.**
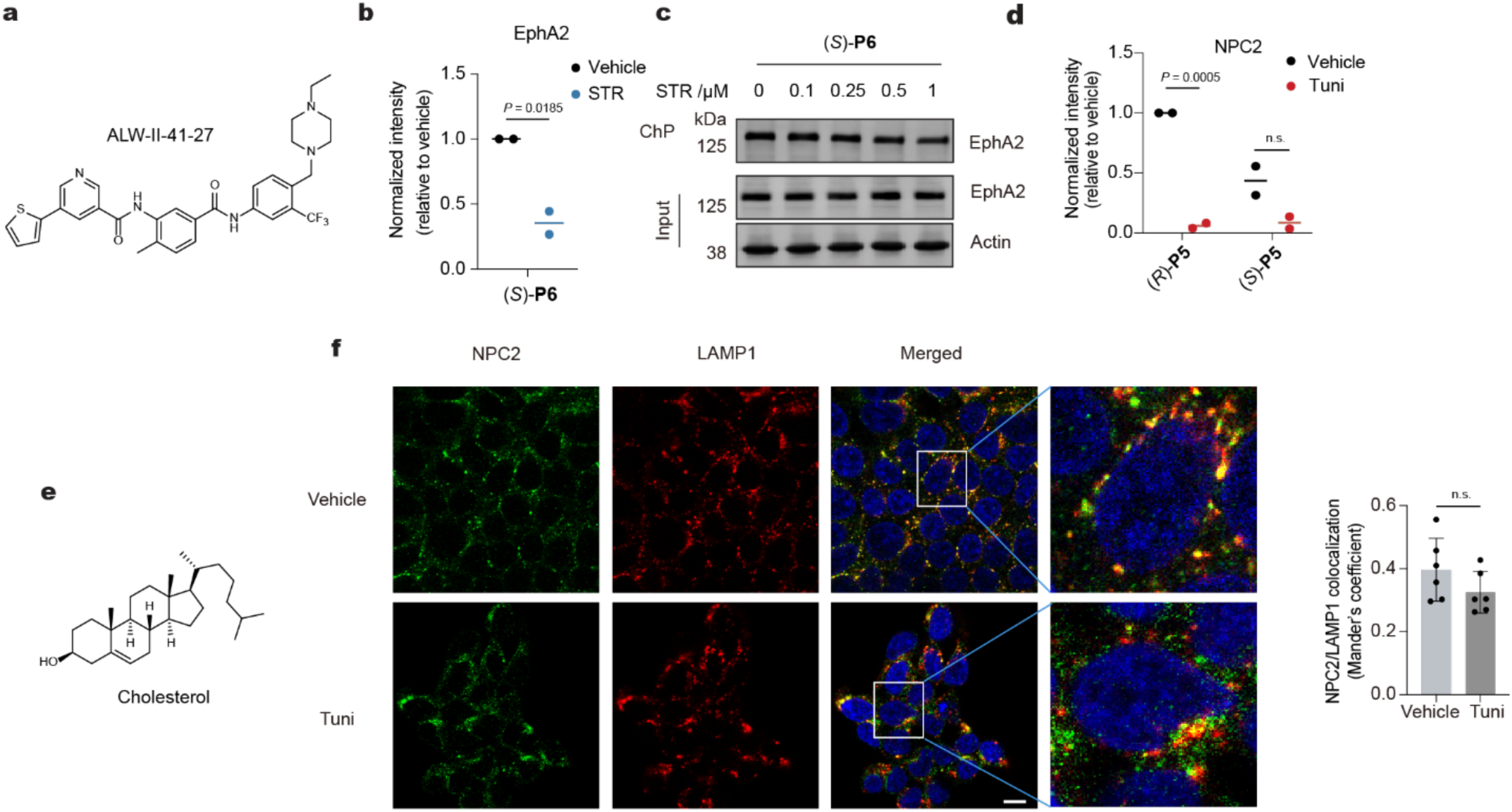
Mapping PTM-dependent binding site ligandability changes in cells. **a**, Chemical structure of ALW-II-41-27, ATP-competitive EphA2 inhibitor. **b**, Quantitation of immunoblot validation of EphA2 in Figure 4l. **c**, Chemo-precipitation (ChP) analysis of probe competition with STR indicating STR does not bind EphA2 competitively. **d**, Quantitation of immunoblot validation of NPC2 in Figure 4p. **e**, Chemical structure of NPC2 substrate cholesterol. **f**, Immunostaining of NPC2 in HEK293T cells with Tuni treatment for 24 h (left) and quantitation (right). Cells were co-stained with LAMP1 as a lysosome marker. Scale bar, 10 μm. Colocalization quantitation were based on cells from six independent fields per condition and presented as mean ± s.d. Immunoblots were quantified from two independent biological replicates and plotted as the mean. Gel images are representative of two independent experiments. Significance was determined using a two-tailed Student’s *t*-test. n.s., not significant.

**Extended Data Fig. 9.**
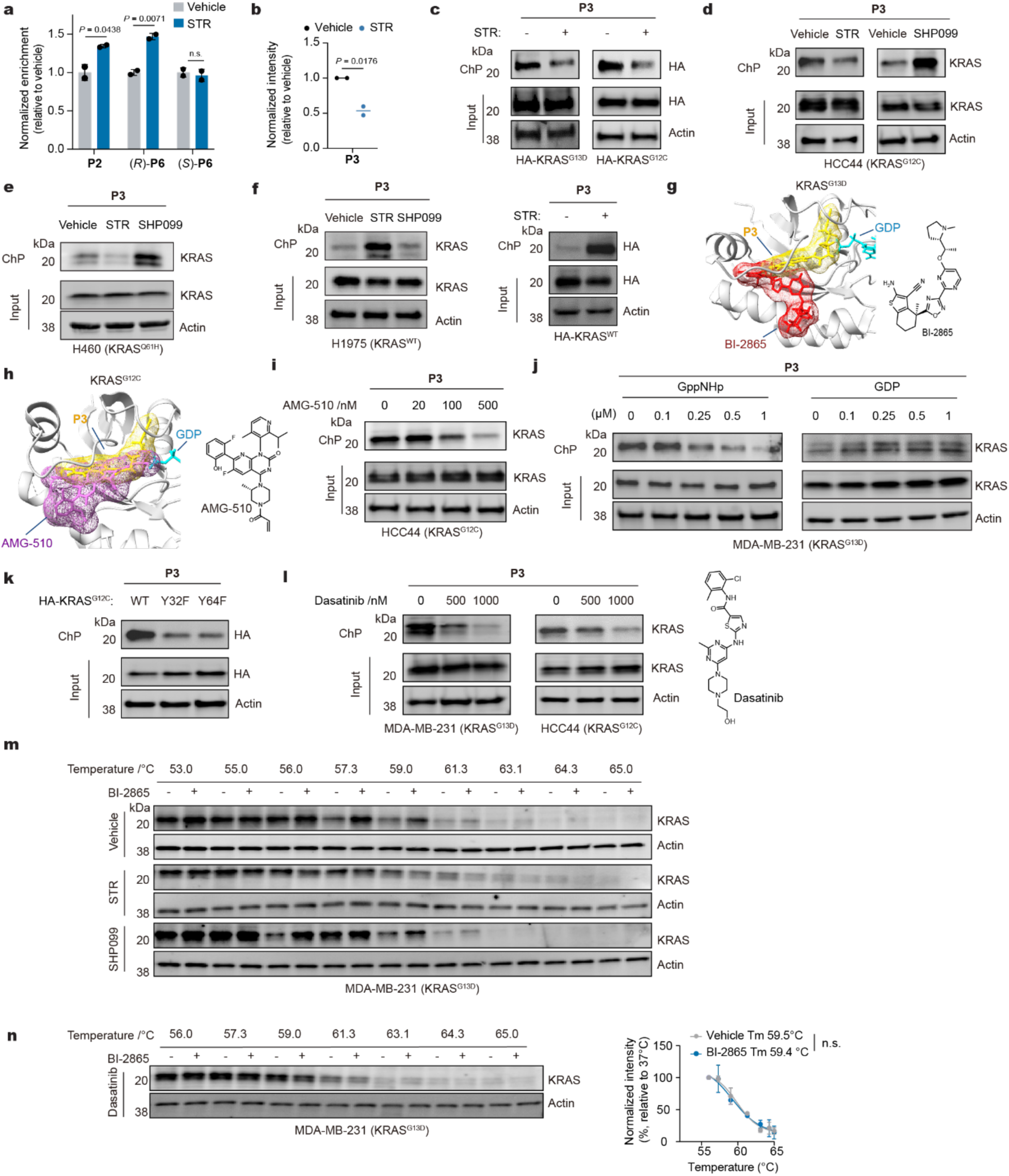
Phosphorylation status impacts KRAS ligandability. **a**, Proteomic profile of **P2** and (*R*/*S*)-**P6** binding to KRAS^G13D^ following STR treatment in MDA-MB-231 cells. **b**, Quantitation of immunoblot replicates in Fig. 5b. **c**, ChP analysis of **P3** binding in LX293T cells expressing HA-KRAS^G13D^ or HA-KRAS^G12C^ following treatment with STR (250 nM, 3 h). **d**, ChP analysis of **P3** binding to KRAS^G12C^ after treatment with STR (250 nM, 3 h) and SHP099 (500 nM, 0.5 h) in HCC44 cells. **e**, ChP analysis of **P3** binding to KRAS^Q61H^ after treatment with STR (250 nM, 3 h) and SHP099 (500 nM, 0.5 h) in H460 cells. **f**, ChP analysis of **P3** binding to KRAS^WT^ after treatment with STR (250 nM, 3 h) or SHP099 (500 nM, 0.5 h) in H1975 cells (left) or in LX293T cells expressing HA-KRAS^WT^ (right). **g**-**h**, Superimposed structure of KRAS^G13D^ docked with **P3** and BI-2865 (PDB: 8B00) and KRAS^G12C^ with **P3** and AMG510 (PDB 6OIM). **i**, ChP analysis P3 binding to KRAS^G12C^ when co-incubated with AMG510 at indicated concentrations in HCC4 cells. **j**, ChP analysis of **P3** binding to KRAS^G13D^ when co**-**incubated with increasing concentration of GppNHp (left) and GDP (right) in MDA-MB-231 cells. **k**, ChP analysis of **P3** binding to HA-KRAS^G12C^ Y32F and Y64F recombinantly expressed in LX293T cells incubated. **l**, ChP analysis of **P3** binding to KRAS^G13D^ in dasatinib-treated MDA-MB-231 cells and KRAS^G12C^ in HCC44 cells at indicated concentrations. **m**-**n**, Cellular thermal shift assay (CETSA) temperature gradient immunoblots of KRAS^G13D^ stabilization with BI-2865 (10 nM) in STR- and SHP099-treated cells (**m**) and in dasatinib treated cells (**n**). Proteomic data were obtained from two independent biological replicates and are presented as mean ± s.d. Immunoblots were quantified from two independent biological replicates and plotted as the mean. Gel images are representative of two independent experiments. Significance was determined using a two-tailed Student’s *t*-test. n.s., not significant. Associated datasets are provided in Supplementary Tables 1 and 4.

**Extended Data Fig. 10.**
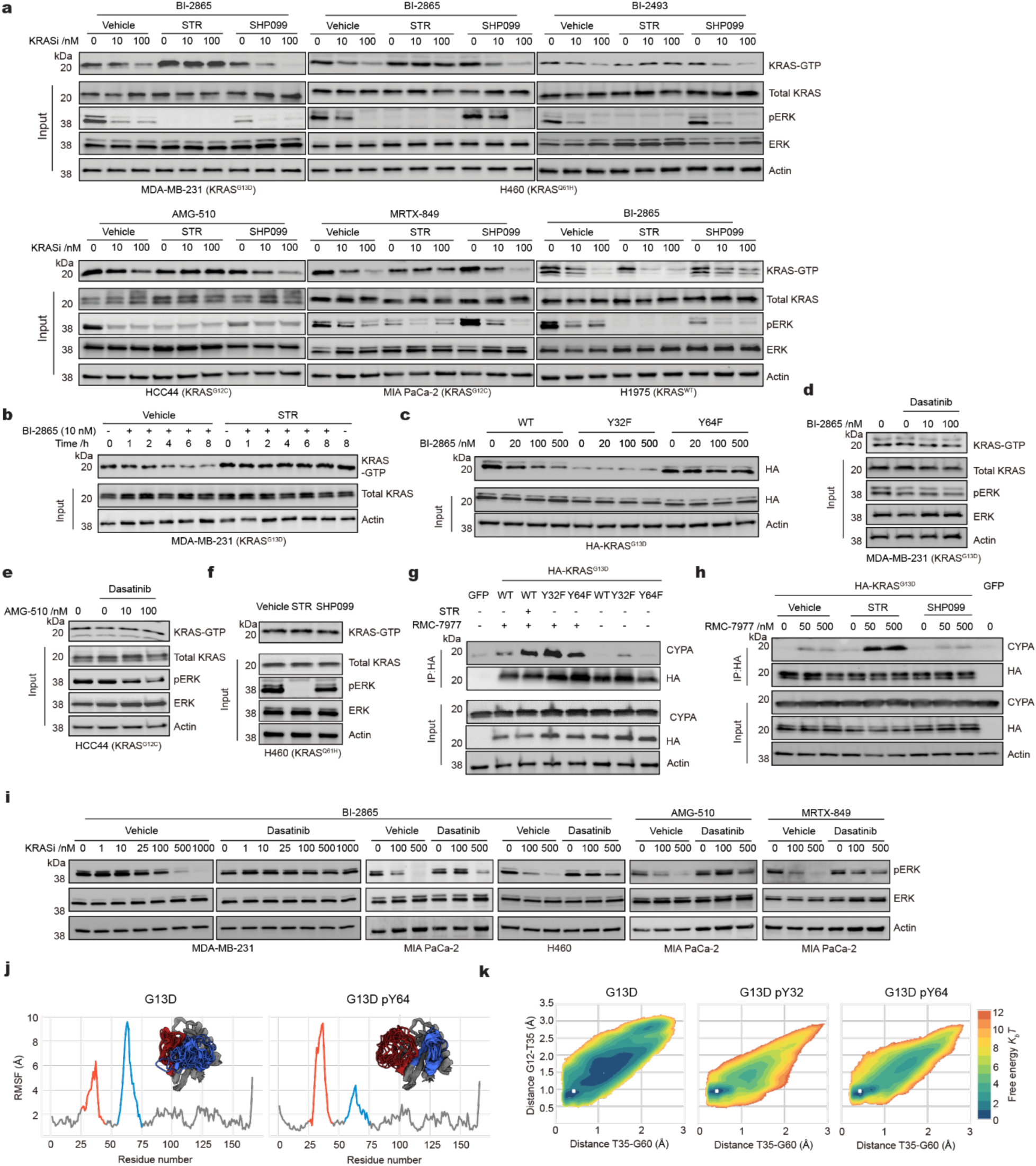
Influence of phosphorylation status on KRAS inhibitor activity. **a**, Monitoring KRAS activity using the RAS-binding domain (RBD) pull-down assay in multiple cell lines. Cells treated with staurosporine (STR, 250 nM) or SHP099 (500 nM) were incubated with indicated KRAS inhibitors, the levels of active-state (GTP-bound) of KRAS were determined via pulldown of GST-RBD and immunoblotting. **b**, Immunoblot of GTP-bound KRAS by RBD pull-down assay in MDA-MB-231 cells. Cells were pre-treated with STR (250 nM, 3 h) and co-treated with BI-2865 and STR for indicated time. **c**, Immunoblot and quantification plot of mutant GTP-bound HA-KRAS^G13D^ levels in LX293T cells expressing HA-tagged KRAS^G13D^ constructs containing Y32 and Y64 site-specific mutations. Cells were treated with BI-2865 for 5 hours followed by immunoblot after HA-tag pulldown. **d,** Immunoblot of GTP-bound KRAS by RBD pull-down assay in dasatinib-treated (1 µM, 1 h) MDA-MB-231 cells with BI-2865 (1 h). **e,** Immunoblot of GTP-bound KRAS by RBD pull-down assay in dasatinib-treated (1 µM, 1 h) HCC44 cells with AMG-510 (1 h). **f**, Immunoblot of GTP-bound KRAS^Q61H^ by RBD pull-down assay in STR (250 nM) or SHP099 (500 nM) treated in H460 cells. **g**, LX293T cells expressing HA-tagged KRAS^G13D^ Y32F and Y64F mutants were treated with RMC-7977 (500 nM, 2 hours) with STR followed by HA-pulldown and CYPA immunoblot. **h**, LX293T cells expressing HA-tagged KRAS^G13D^ were treated with STR (250 nM) or SHP099 (500 nM), and RMC-7977 with indicated concentration for 2 hours followed by HA-pulldown and CYPA immunoblot. **i,** Immunoblot of pERK signaling levels restored following dasatinib pre-treatment (1 µM, 1 h), and KRAS inhibitors co-treatment in indicated cell lines (1 h for MDA-MB-231, MIA PaCa-2; 0.5 h for H460). **j**, Per-residue root-mean-square-fluctuation (RMSF) plots quantitatively illustrate the differential flexibility of SW-I (red) and SW-II (blue) regions for KRAS^G13D^ and KRAS^G13D^ pY64. Representative structural ensembles extracted from clustering unbiased simulations are shown for each system, highlighting the conformational heterogeneity of these KRAS variants. **k**, FES for KRAS^G13D^ and the two potential phosphorylated variants, Y32 and Y64. As a reference, the BI-2865 binding-competent conformation is shown as a white cross. For the phosphorylated variants, a single energy minimum exists and matches the BI-2865 conformation. Results are representative of two or three biological replicates.

