## Supplemental Information for "Post-Translational Modifications Remodel Proteome-Wide Ligandability"

This PDF file includes:

- Supplementary Figures 1-12
- Supplementary Table 1-15
- Supplementary note

Other Supplementary Material for this manuscript includes the following:

- Supplementary Tables 1 to 14 (Supplementary Data 1, provided in excel file)

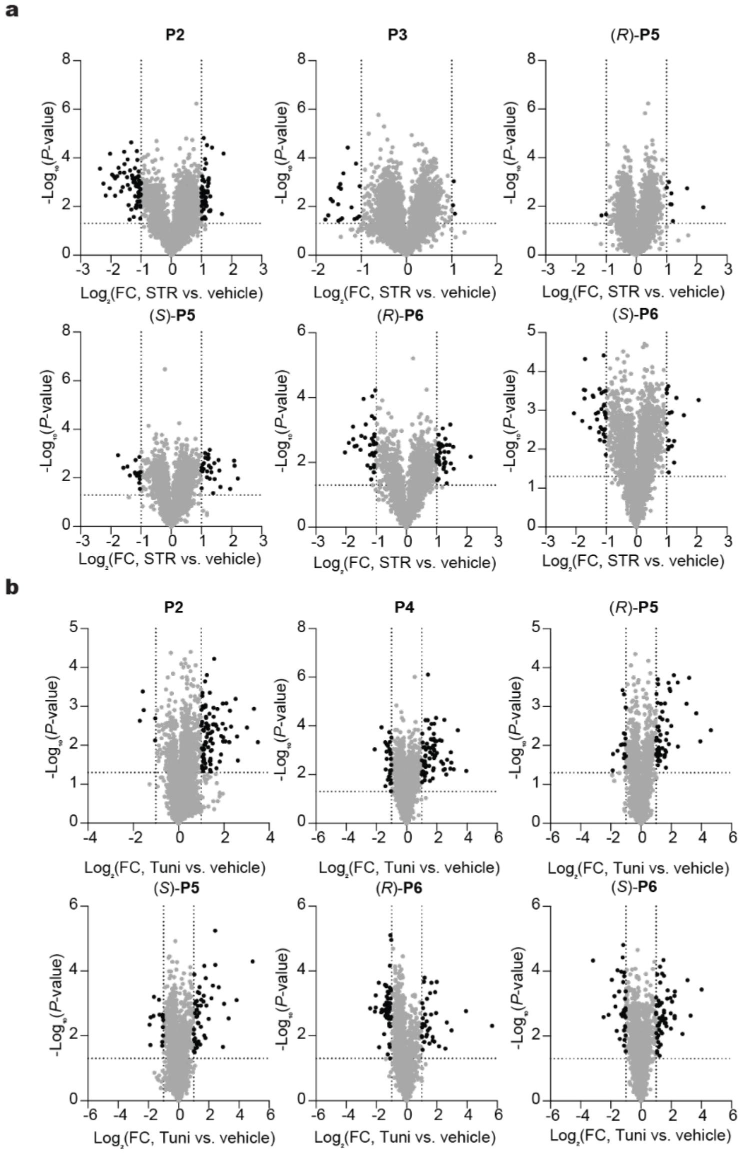

**Supplementary Fig.1 Proteomic profiling of PTM-dependent ligandability changes in cells**

**a-b**, Volcano plots illustrating PTM-dependent ligandability changes detected with the FFF-probes. **a**, Phosphorylation-dependent changes; **b**, N-linked glycosylation-dependent changes. Vertical dashed lines represent a 2-fold change (FC) threshold, while horizontal dashed lines denote a *p*-value threshold of 0.05. Black dots highlight proteins that meet both significance thresholds.

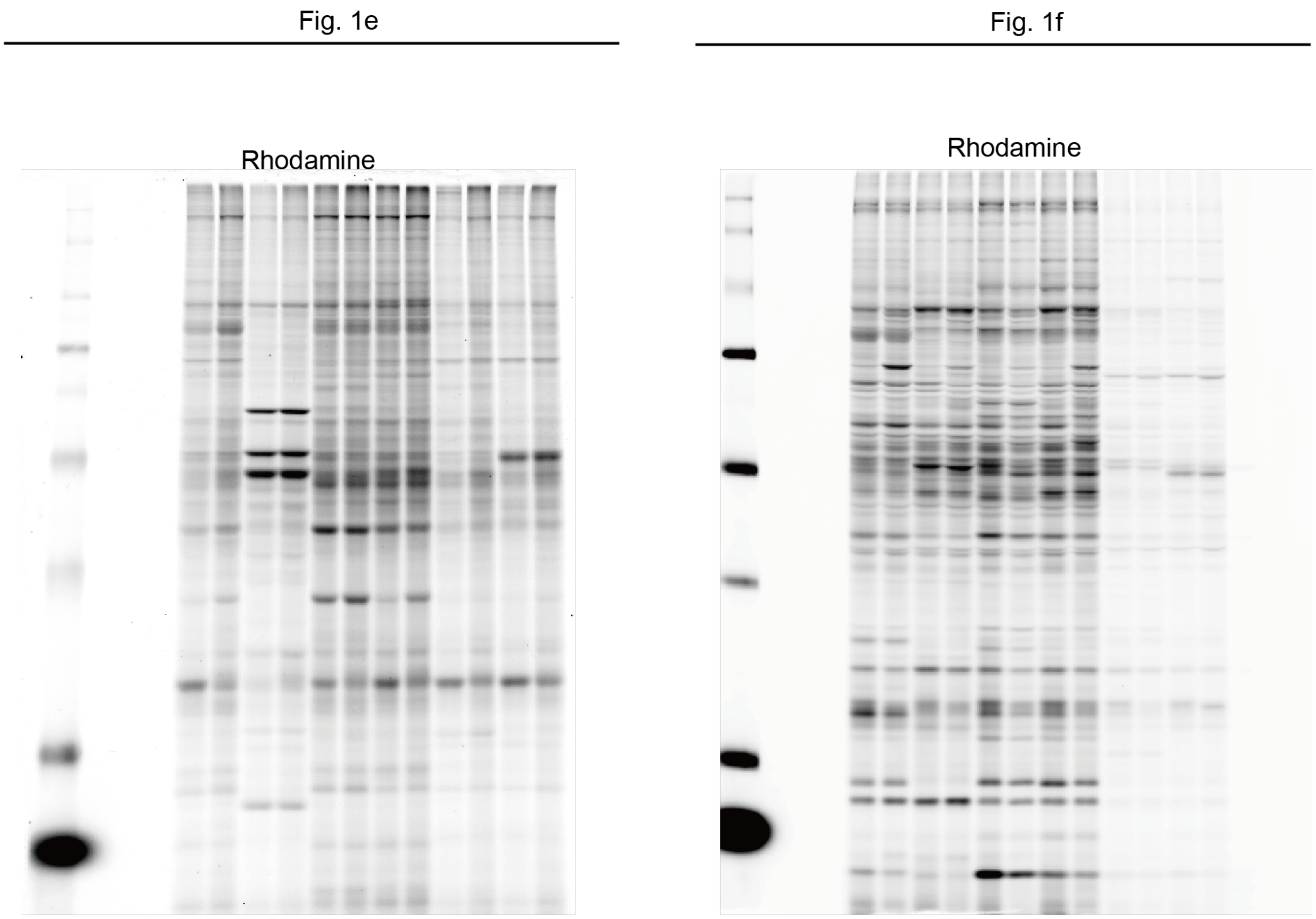

**Supplementary Fig. 2. Uncropped western blot images in Fig. 1.**

Fig. 3b

P2, JAK1 (ChP)

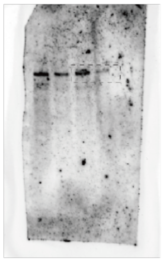

P2, JAK1 (input)

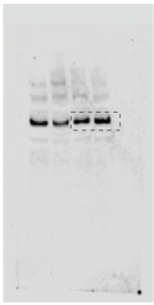

P2, Actin (input)

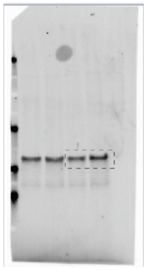

Left, P5, JAK1 (ChP)
Right, P3, JAK1 (ChP)

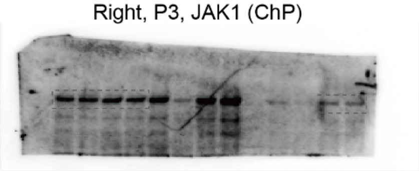

Left, P5, JAK1 (input)
Right, P3, JAK1 (input)

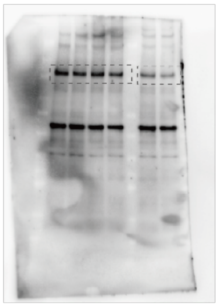

Left, P5, Actin (input)
Right, P3, Actin (input)

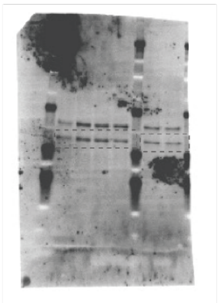

P6, JAK1 (ChP)

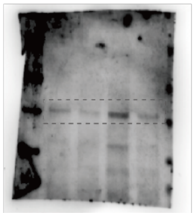

P6, JAK1 (input)

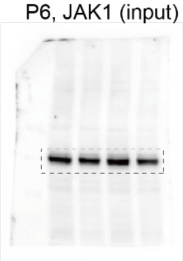

P6, Actin (input)

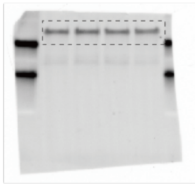

Fig. 3d

Left, P3, ATP6V1E1 (ChP)
Right, P2, ATP6V1E1 (ChP)

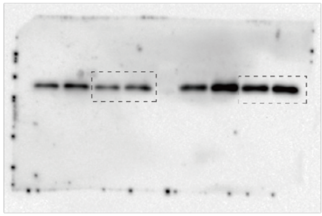

Left, P3, ATP6V1E1 (input)
Right, P2, ATP6V1E1 (input)

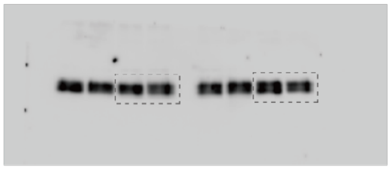

Left, P3, Actin (input)
Right, P2, Actin (input)

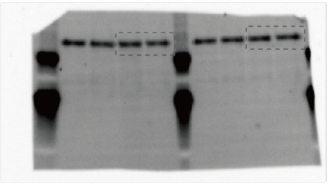

P5, ATP6V1E1 (ChP)

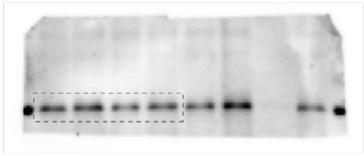

P5, ATP6V1E1 (input)

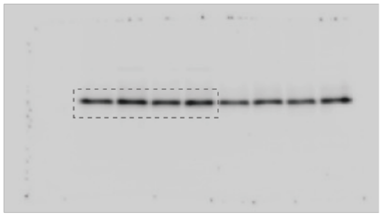

P5, Actin (input)

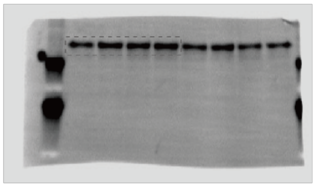

P6, ATP6V1E1 (ChP)

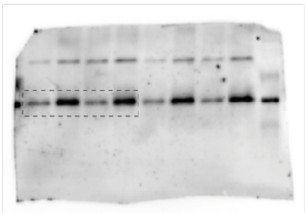

P6, ATP6V1E1 (input)

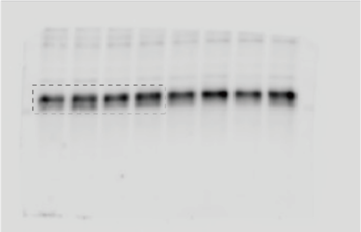

P6, Actin (input)

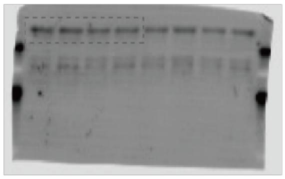

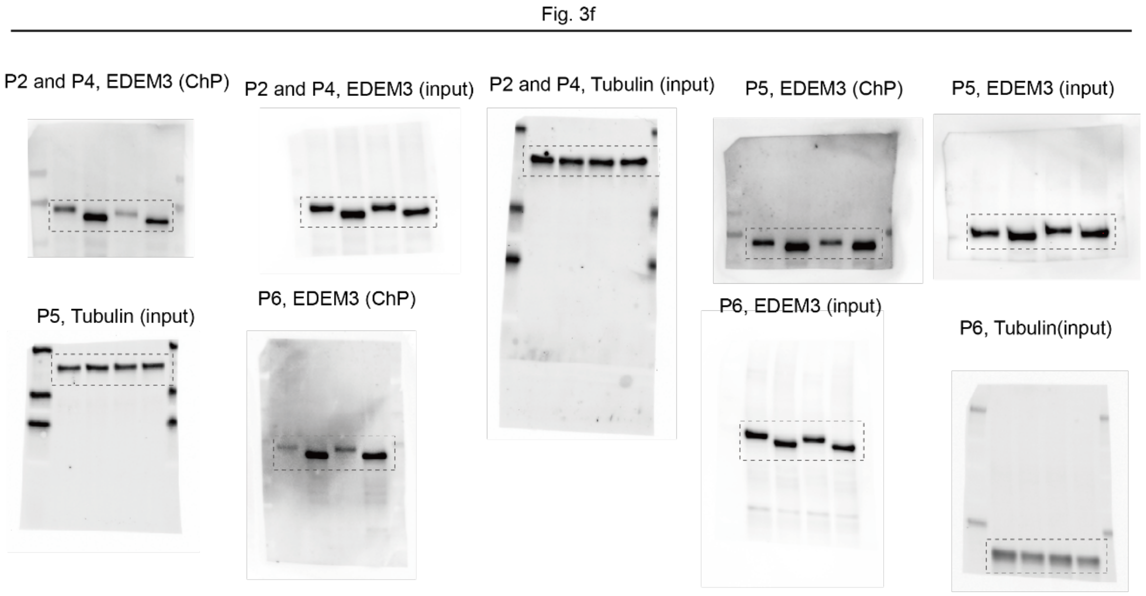

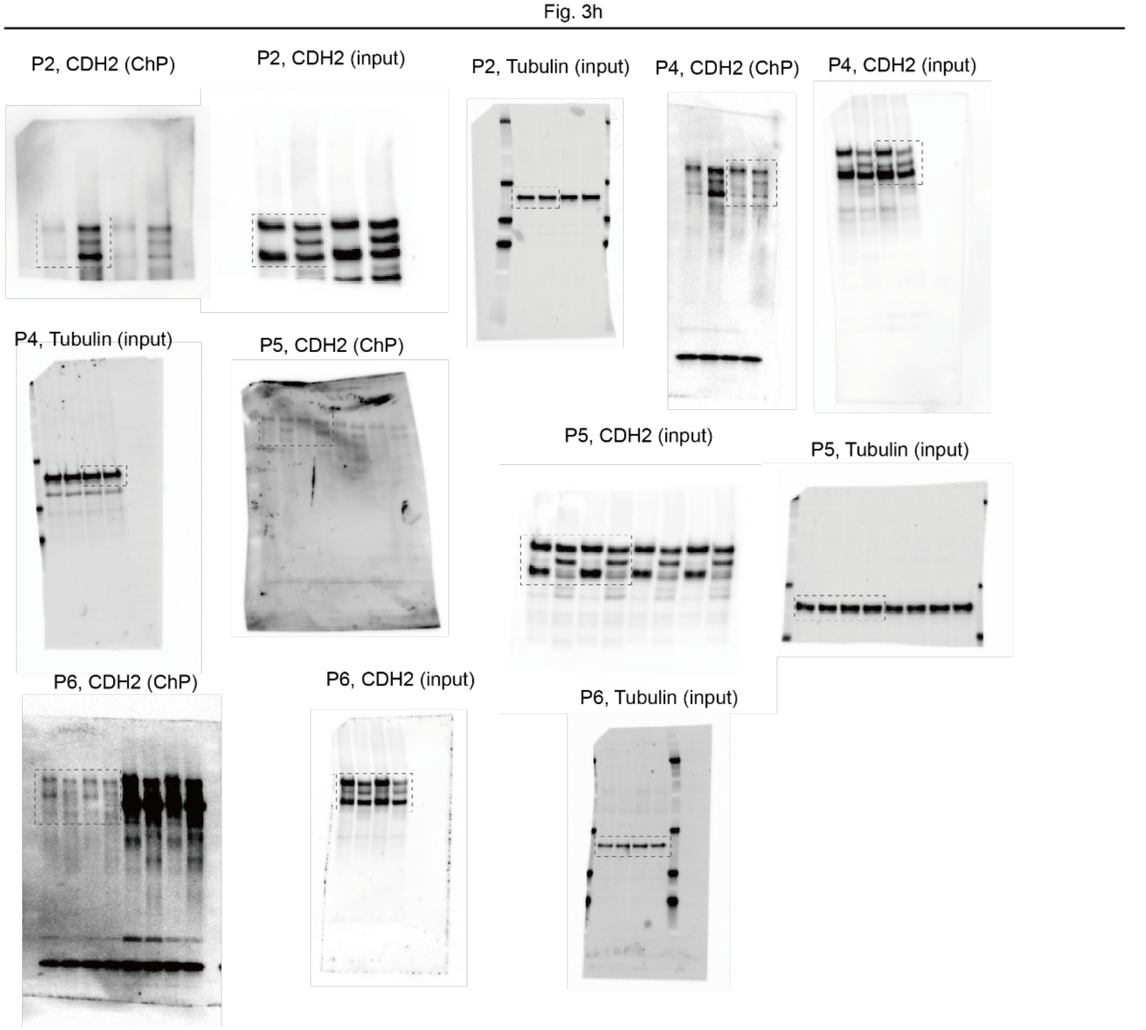

**Supplementary Fig. 3. Uncropped western blot images in Fig. 3.**

Fig. 4g

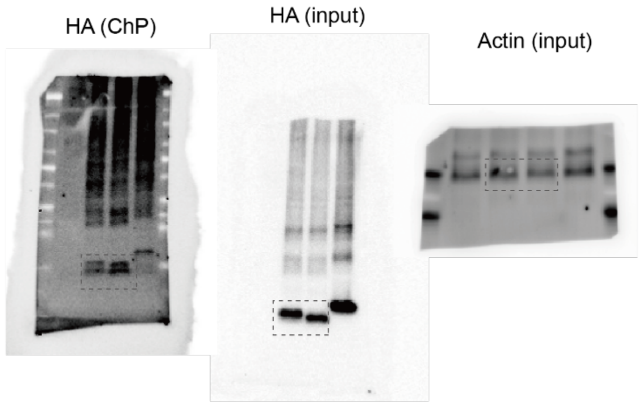

Fig. 4h

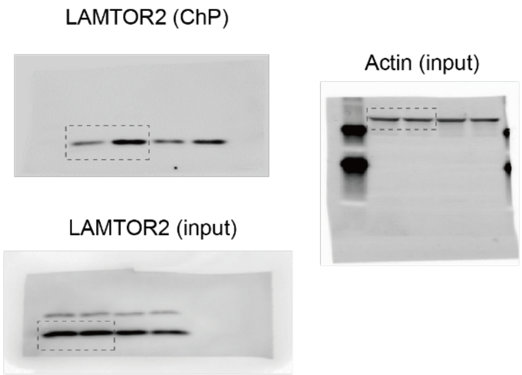

Fig. 4k

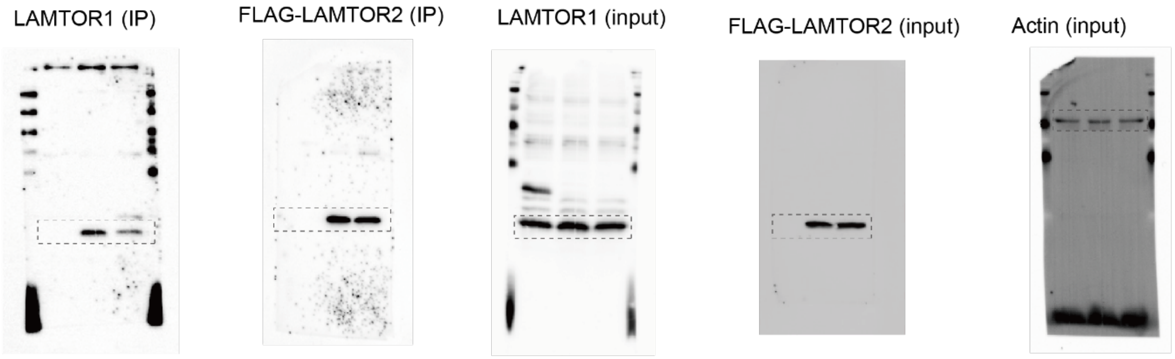

Fig. 4l

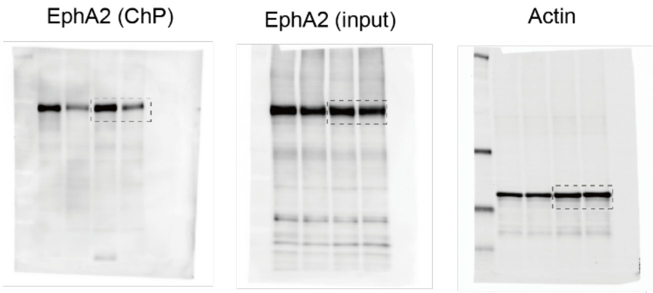

Fig. 4o

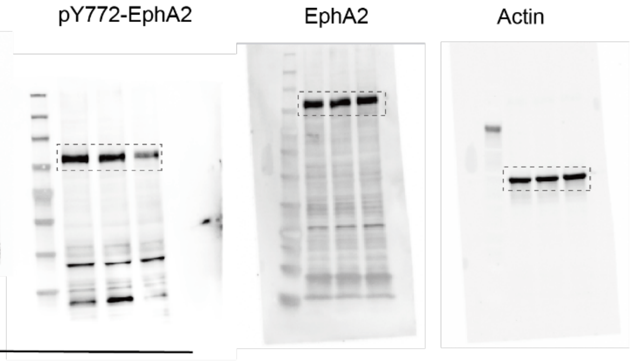

Fig. 4m

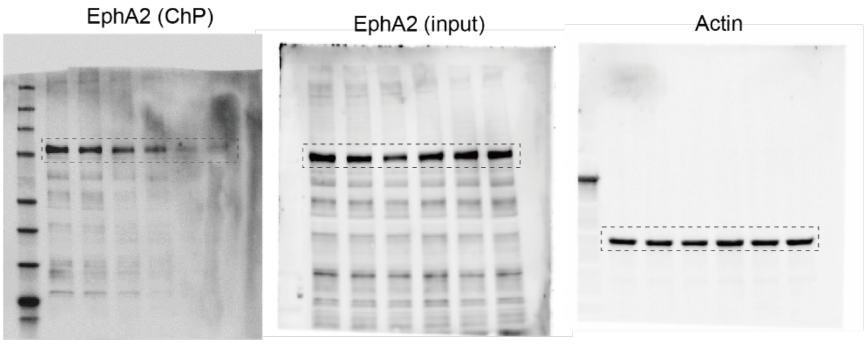

Fig. 4p

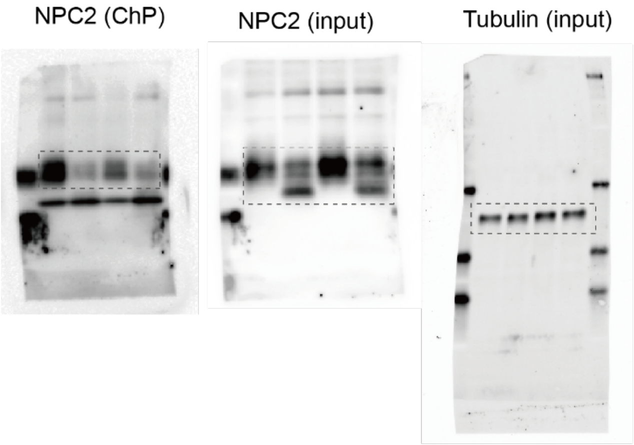

Fig. 4r

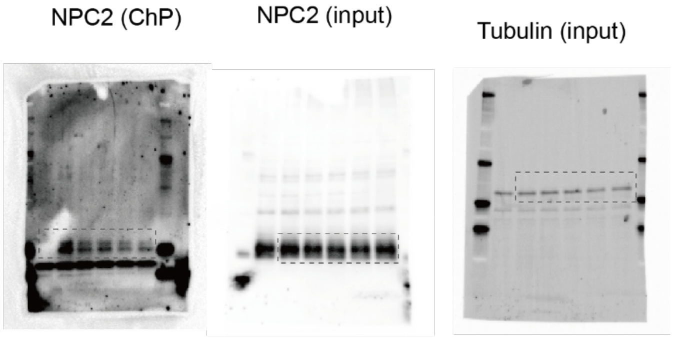

Fig. 4s

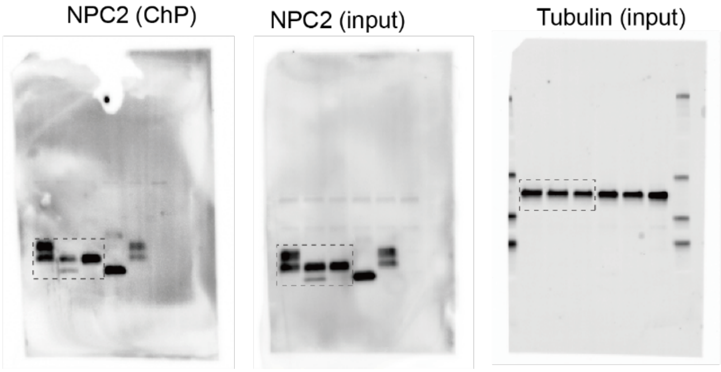

**Supplementary Fig. 4. Uncropped western blot images in Fig. 4.**

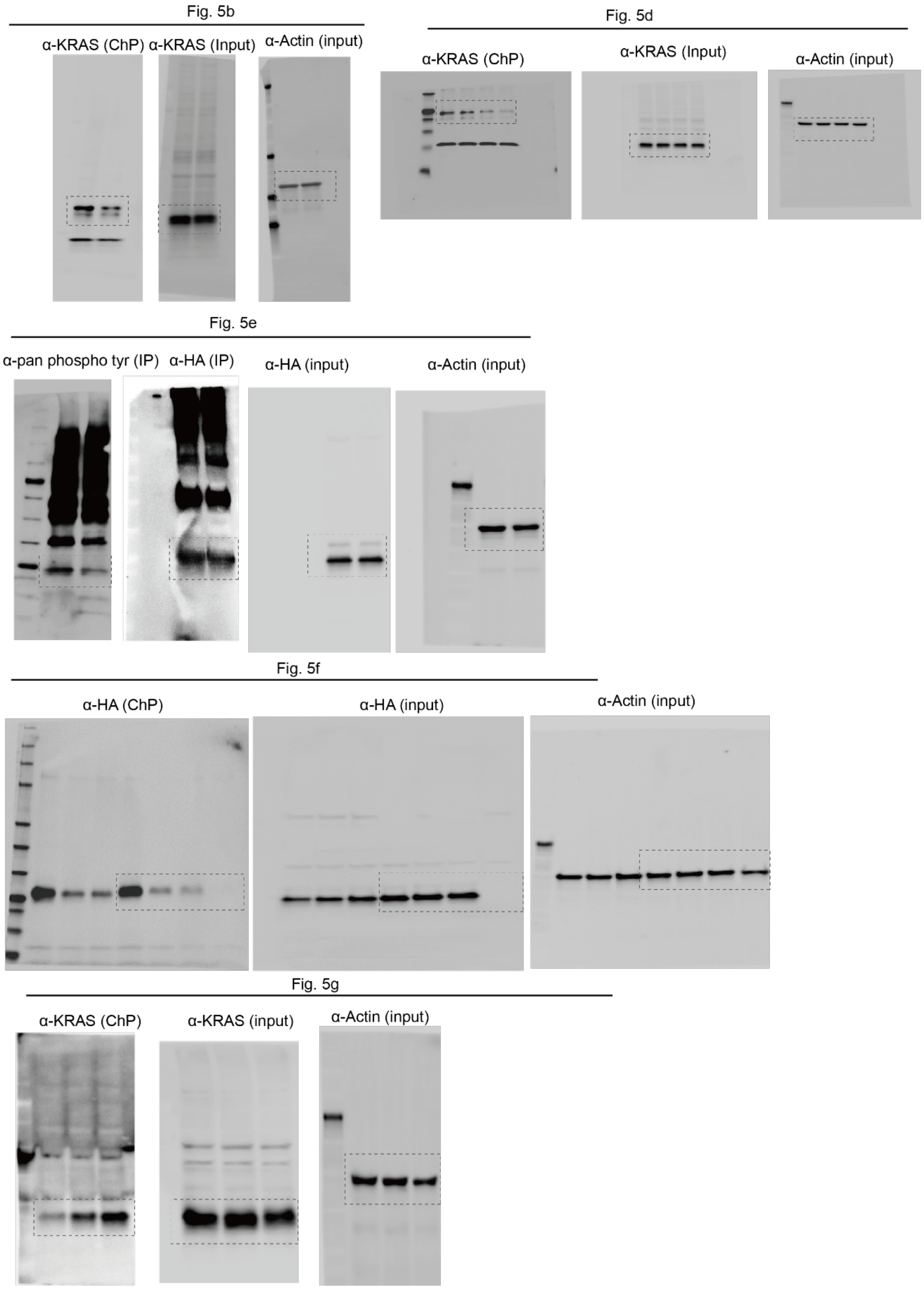

**Supplementary Fig. 5. Uncropped western blot images in Fig. 5.**

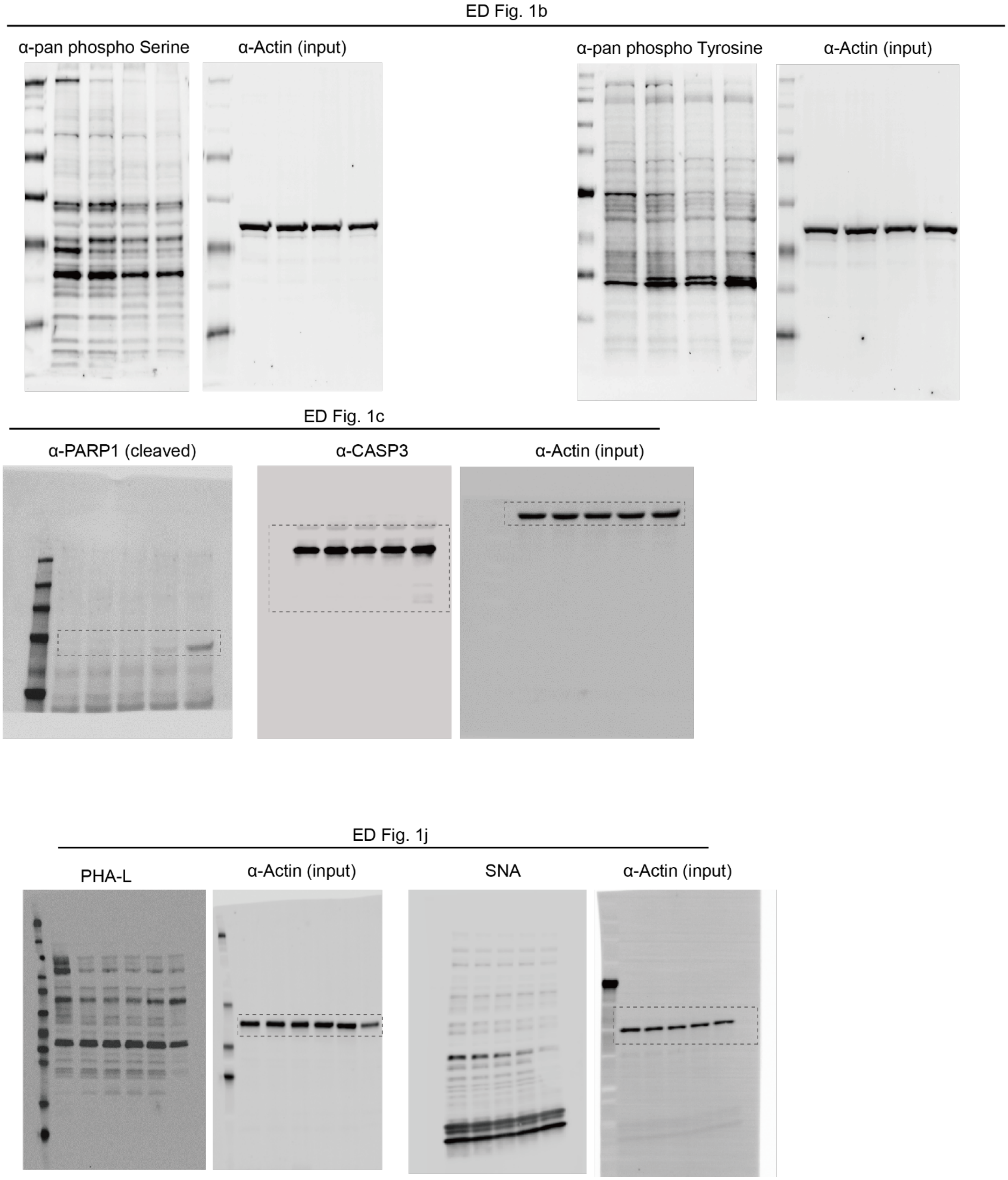

**Supplementary Fig. 6. Uncropped western blot images in Extended Data Fig. 1.**

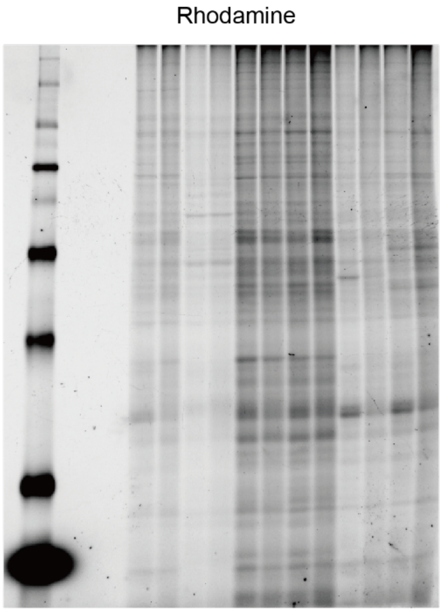

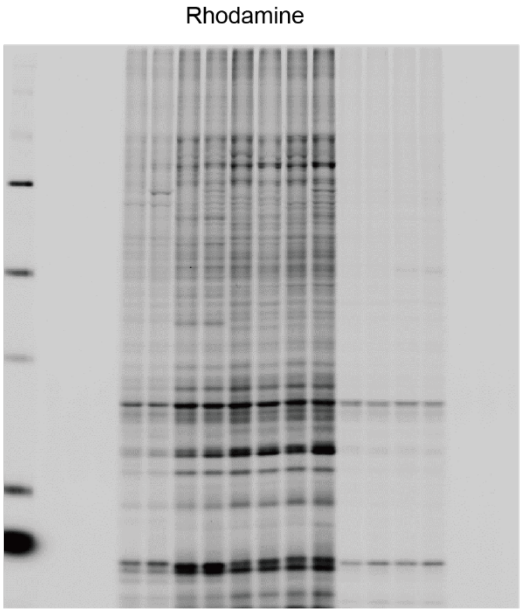

**Supplementary Fig. 7. Uncropped western blot images in Extended Data Fig. 2.**

ED Fig. 4a

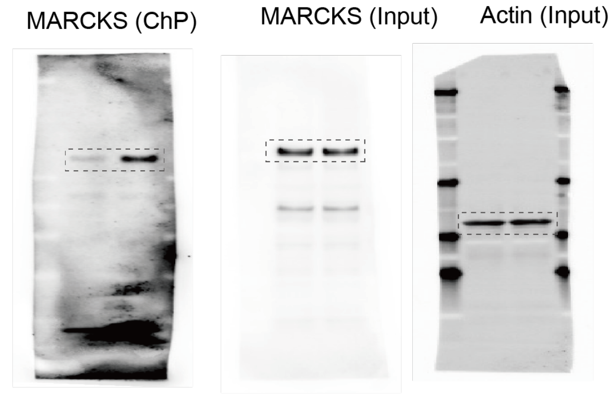

ED Fig. 4b

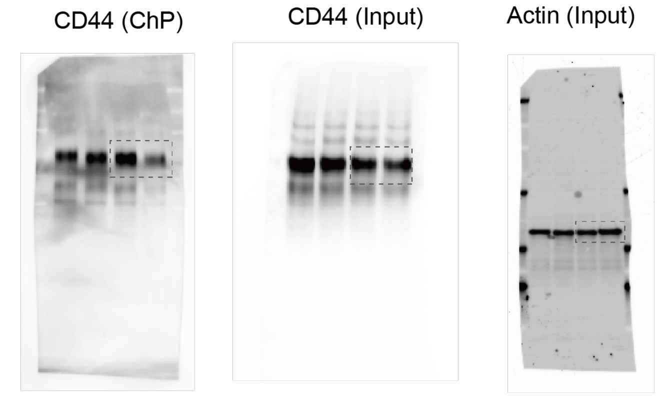

ED Fig. 4c

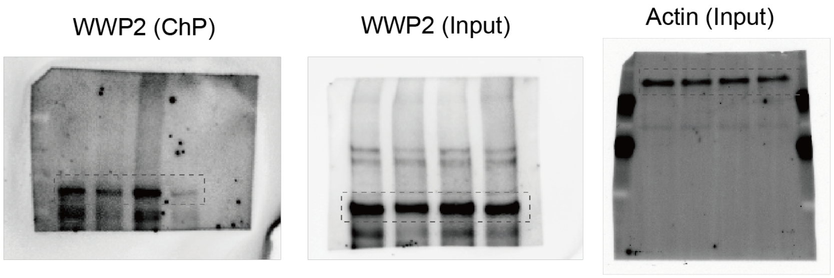

ED Fig. 5a

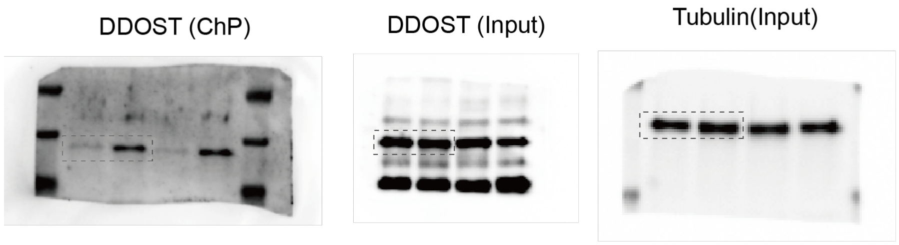

**Supplementary Fig. 8. Uncropped western blot images in Extended Data Fig. 5.**

ED Fig. 6c

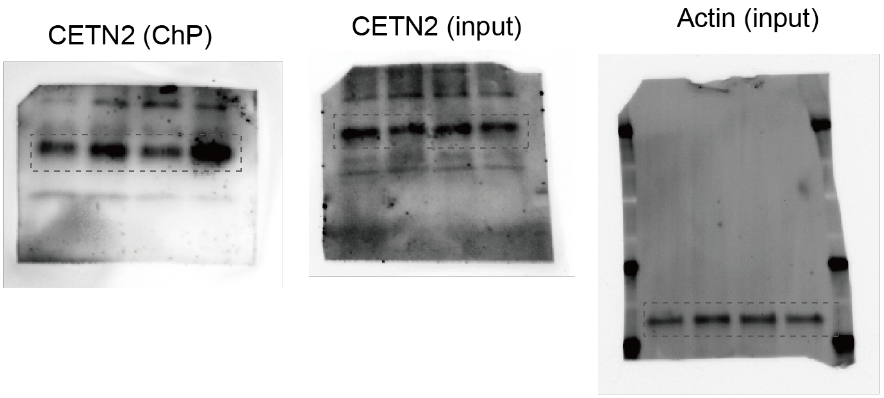

ED Fig. 6f

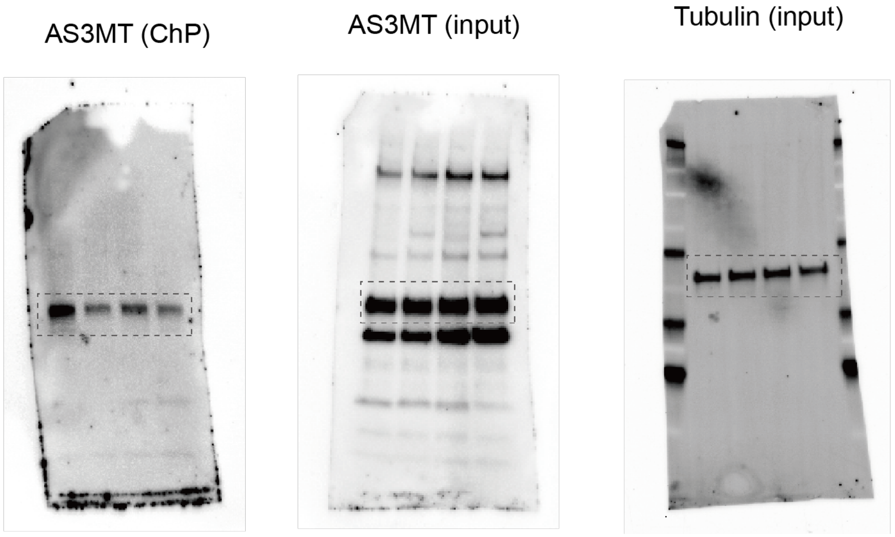

**Supplementary Fig. 9. Uncropped western blot images in Extended Data Fig. 6.**

### ED Fig. 8c

EphA2 (ChP)

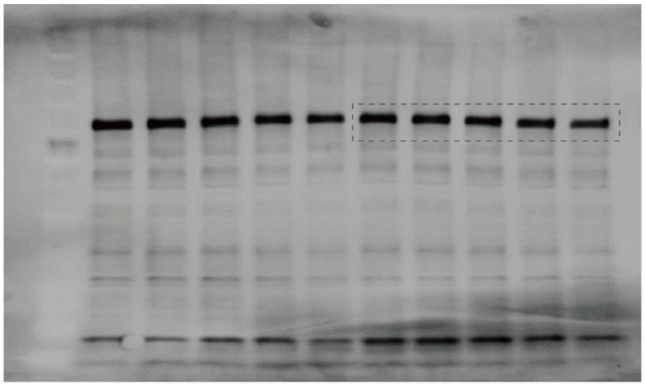

EphA2 (input)

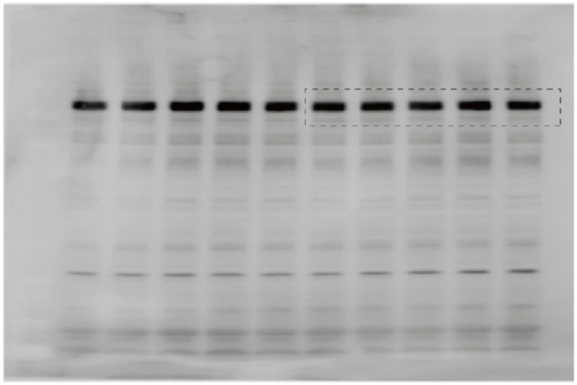

Actin (input)

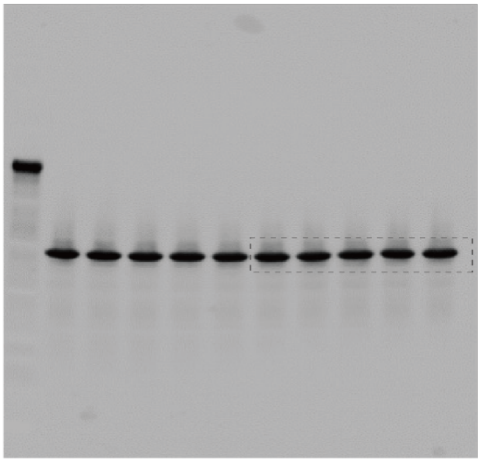

**Supplementary Fig. 10. Uncropped western blot images in Extended Data Fig. 8.**

ED Fig. 9c

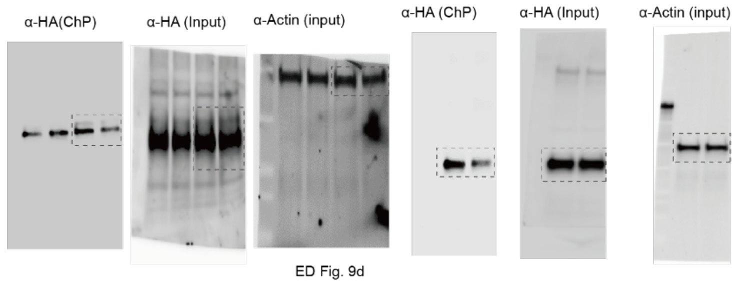

ED Fig. 9d

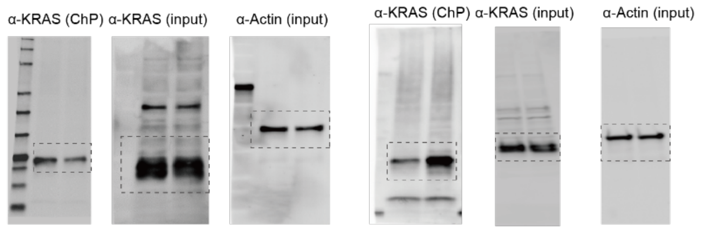

ED Fig. 9e

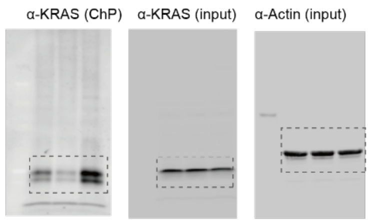

ED Fig. 9f

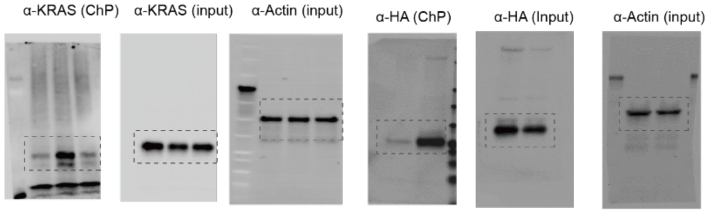

ED Fig. 9i

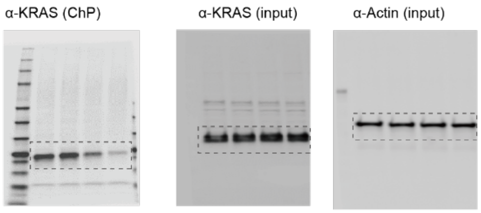

ED Fig. 9j

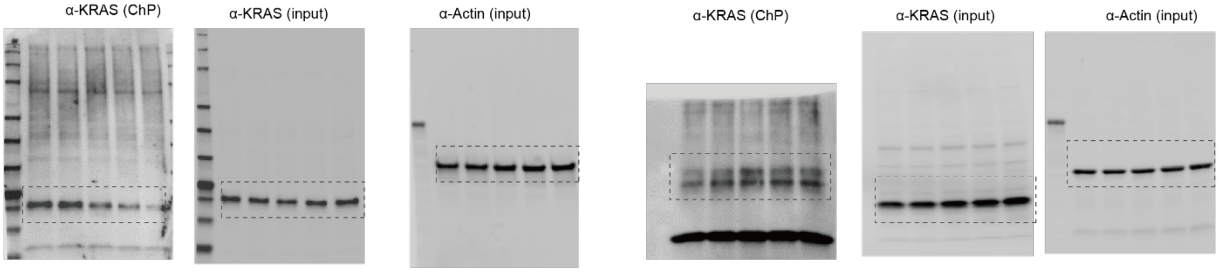

ED Fig. 9k

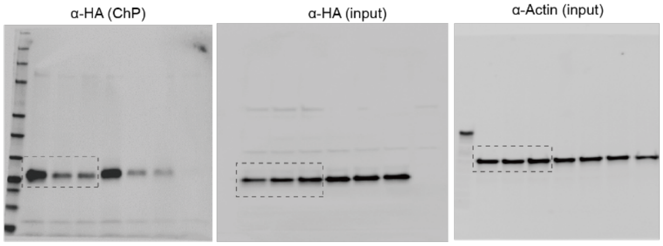

ED Fig. 9l

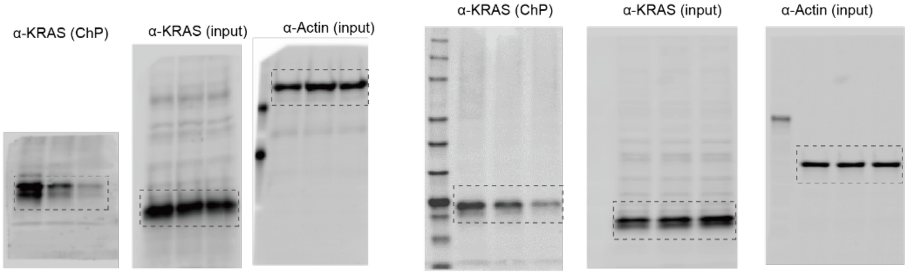

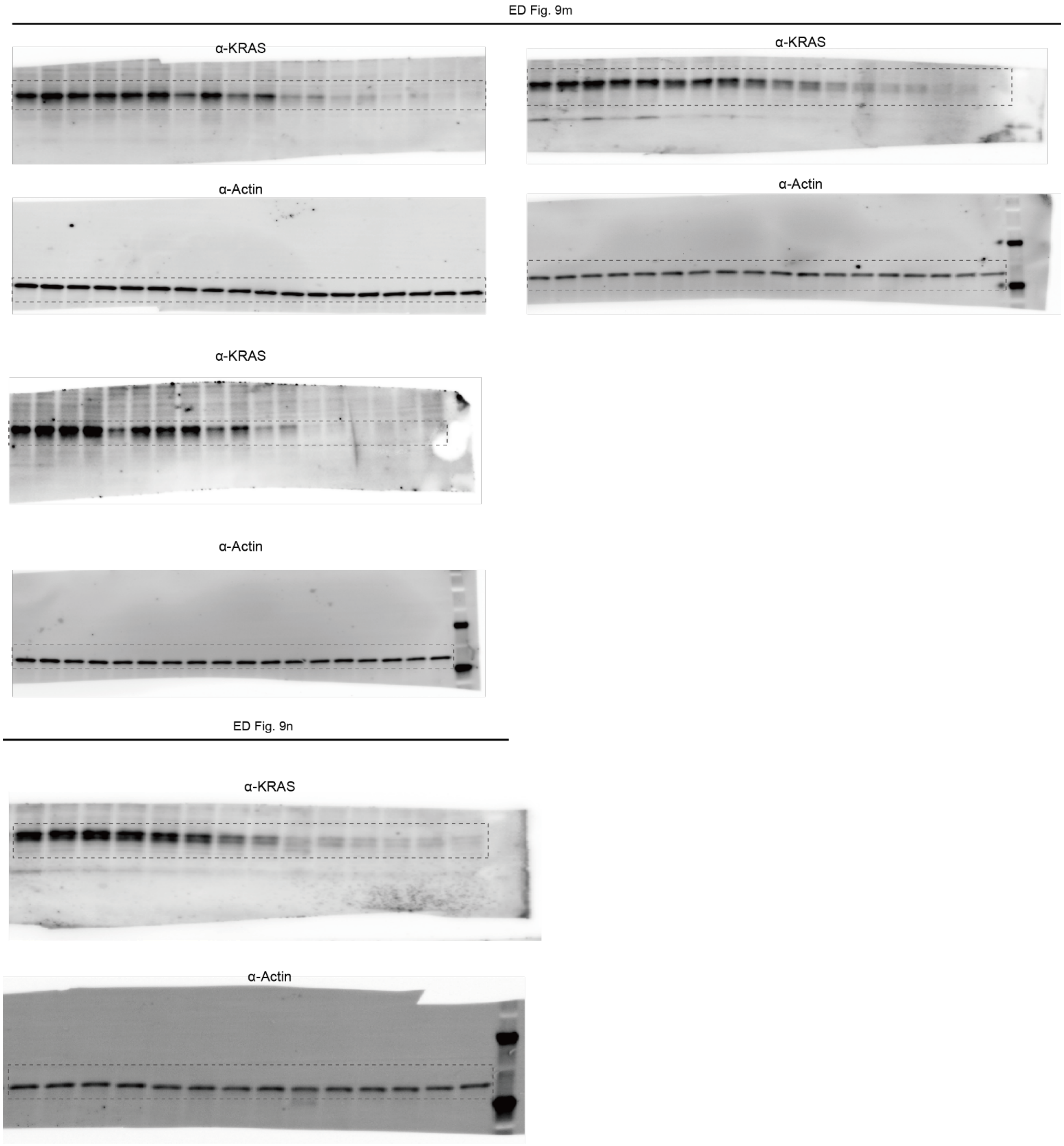

**Supplementary Fig. 11. Uncropped western blot images in Extended Data Fig. 9.**

ED Fig. 10a

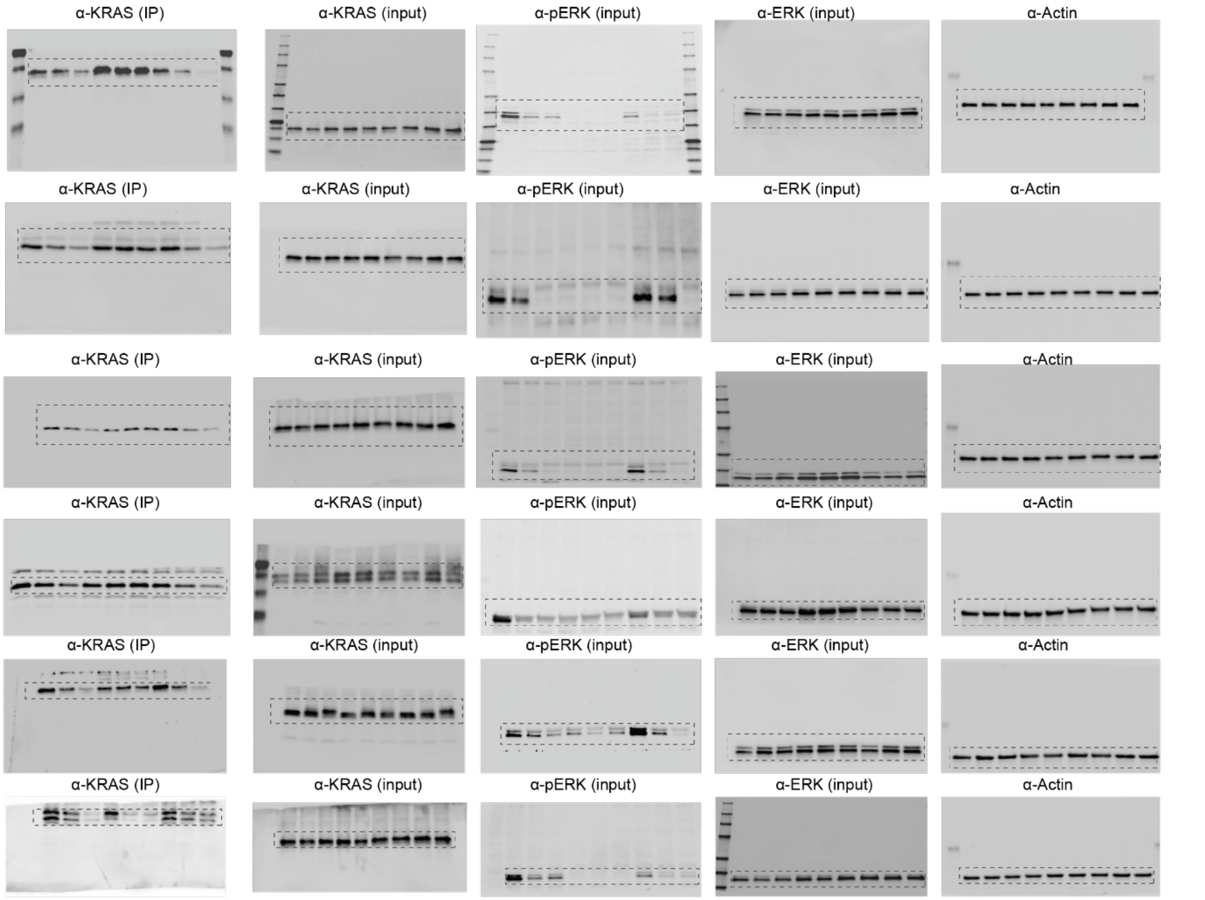

ED Fig. 10b

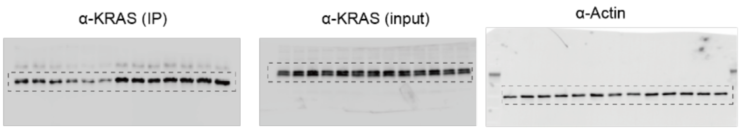

ED Fig. 10c

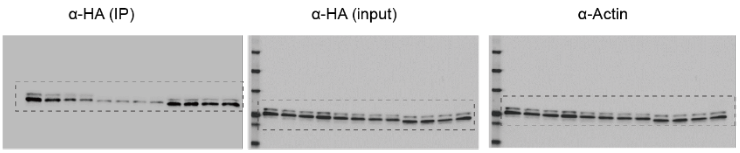

ED Fig. 10d

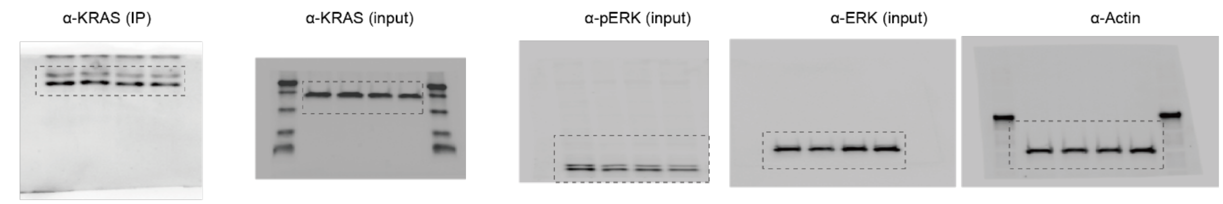

ED Fig. 10e

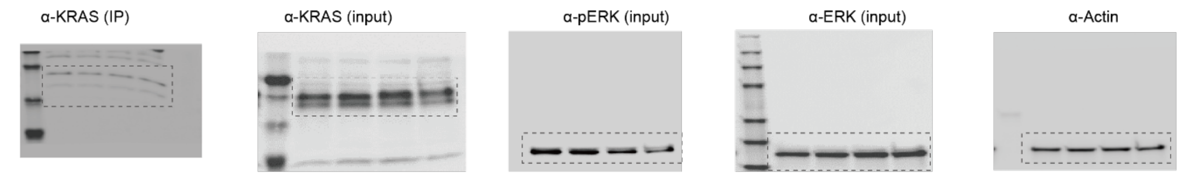

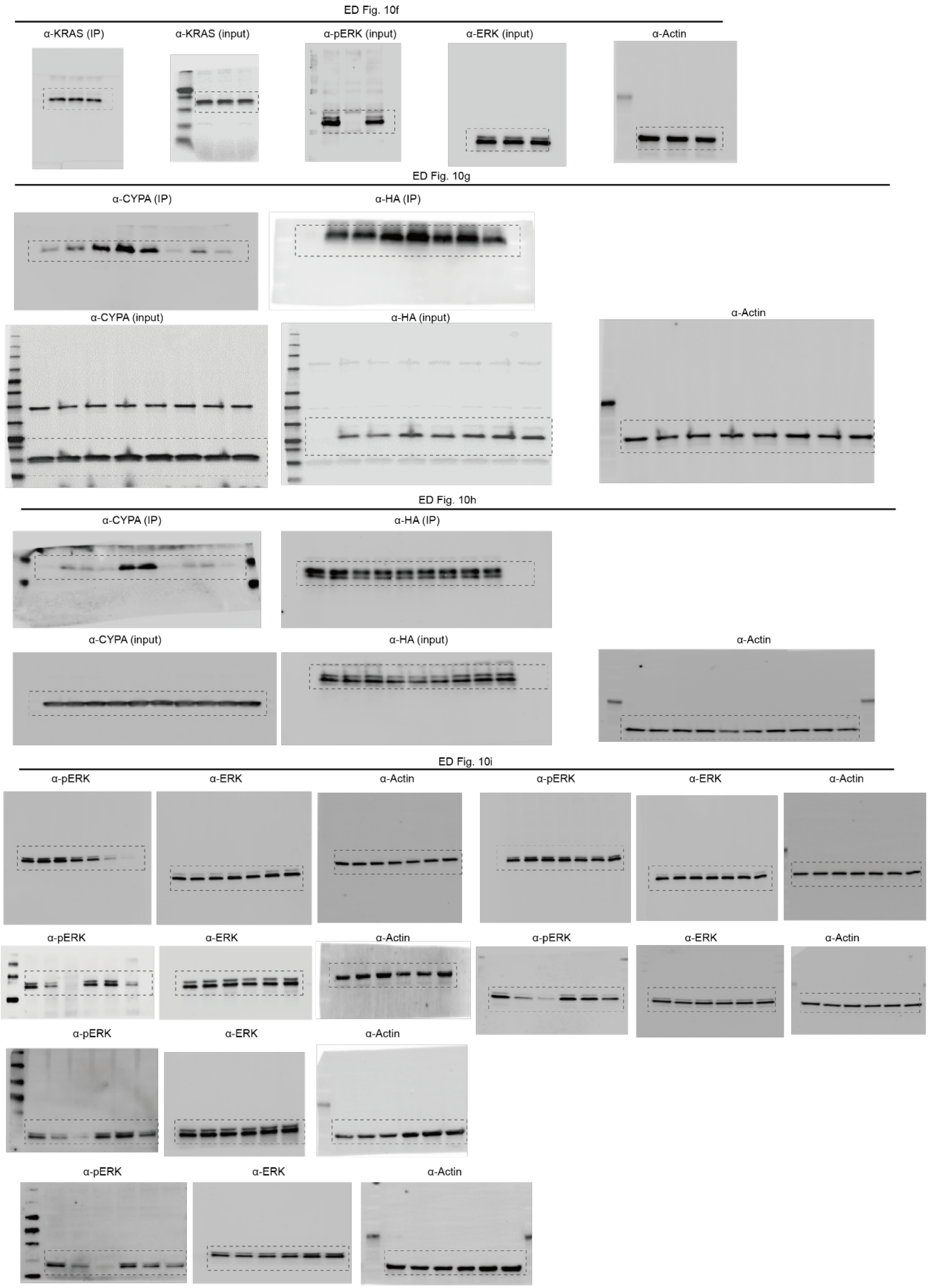

**Supplementary Fig. 12. Uncropped western blot images in Extended Data Fig. 10.**

**Supplementary Tables**

**Supplementary Table 1. Whole-protein proteomics data (P2, phosphorylation-dependent).**

See attached excel table.

**Supplementary Table 2. Whole-protein proteomics data (P3, phosphorylation-dependent).**

See attached excel table.

**Supplementary Table 3. Whole-protein proteomics data (P5, phosphorylation-dependent).**

See attached excel table.

**Supplementary Table 4. Whole-protein proteomics data (P6, phosphorylation-dependent).**

See attached excel table.

**Supplementary Table 5. Phosphorylation-dependent whole protein unenriched proteomics data.**

See attached excel table.

**Supplementary Table 6. Whole-protein proteomics data (P2 and P5, glycosylation-dependent).**

See attached excel table.

**Supplementary Table 7. Whole-protein proteomics data (P4 and P6, glycosylation-dependent).**

See attached excel table.

**Supplementary Table 8. Glycosylation-dependent whole protein unenriched proteomics data.**

See attached excel table.

**Supplementary Table 9. Phospho-proteomics enriched with $TiO_2$.**

See attached excel table

**Supplementary Table 10. Phospho-proteomics enriched with FeNTA.**

See attached excel table

**Supplementary Table 11. SoL phosphorylation-dependent proteomics data processed with Proteome Discoverer.**

See attached excel table

**Supplementary Table 12. SoL glycosylation-dependent proteomics data processed with Proteome Discoverer.**

See attached excel table

**Supplementary Table 13. SoL phosphorylation-dependent proteomics data processed with MSFragger.**

See attached excel table

**Supplementary Table 14. SoL glycosylation-dependent proteomics data processed with MSFragger.**

See attached excel table

**Supplementary Table 15. Reagent or resource used in this study**

| REAGENT or RESOURCE | SOURCE | IDENTIFIER |
|---|---|---|
| **Antibodies** | | |
| Anti-KRAS antibody (1:1000) | ProteinTech | 12063-1-AP |
| Anti-pERK antibody (1:1000) | Cell Signaling Technology | 9102 |
| Anti-ERK antibody (1:1000) | Cell Signaling Technology | 9101 |
| Phospho-Tyrosine (P-Tyr-1000) MultiMab (1:500) | Cell Signaling Technology | 8954 |
| JAK1 Monoclonal antibody (1:1000) | Proteintech | 66466-1-lg |
| ATP6V1E1 Polyclonal antibody (1:1000) | Proteintech | 15280-1-AP |
| EDEM3 Polyclonal antibody (1:1000) | Proteintech | 27310-1-AP |
| CDH2 Polyclonal antibody (1:1000) | Proteintech | 22018-1-AP |
| Human EPHA2 antibody (1:200) | R&D | AF3035 |
| Phospho-EphA2 (Tyr772) Antibody | Cell Signaling Technology | 8244 |
| NPC2 Polyclonal antibody (1:1000) | Proteintech | 19888-1-AP |
| Monoclonal anti-Flag M2 antibody (1:2000) | Sigma | F3165 |
| LAMTOR2/ROBLD3 (D7C10) Rabbit mAb (1:1000) | Cell Signaling Technology | 8145 |
| LAMTOR1 Rabbit pAb (1:2000) | Abclonal | A21557 |
| HA-Tag (C29F4) Rabbit mAb (1:2000) | Cell Signaling Technology | 3724 |
| Biotinylated PHA-L (1:200) | Vector laboratories | B-1115-2 |
| Sambucus Nigra Lectin (SNA, EBL), Biotinylated | Vector laboratories | B-1305-2 |
| Phospho-(Ser/Thr) Phe Antibody (1:1000) | Cell Signaling Technology | 9631 |
| PARP1 Polyclonal antibody (1:5000) | Cell Signaling Technology | 9541S |
| CAPS3 Polyclonal antibody (1:1000) | Cell Signaling Technology | 9662S |
| WWP2 Polyclonal antibody (1:1000) | Proteintech | 12197-1-AP |
| MARCKS Polyclonal antibody (1:1000) | Proteintech | 10004-1-lg |
| CD44 Polyclonal antibody (1:1000) | Proteintech | 15675-1-AP |
| CETN2 Polyclonal antibody (1:1000) | Proteintech | 15877-1-AP |
| AS3MT Polyclonal antibody (1:1000) | Proteintech | 27270-1-AP |
| DDOST Polyclonal antibody (1:1000) | Proteintech | 14916-1-AP |
| BiP Rabbit mAb (1:1000) | Cell Signaling Technology | 3177 |
| Anti-human LAMP-1 antibody | Biolegend | 328601 |
| Anti-Actin hFAB (1:10000) | Bio-Rad | 12004164 |
| Anti-Tubulin hFAB (1:10000) | Bio-Rad | 12004165 |

| | | |
|---|---|---|
| DyLight™ 488 Donkey anti-rabbit IgG (minimal x-reactivity) Antibody | Biolegend | 406404 |
| Alexa Fluor® 647 anti-mouse IgG1 Antibody | Biolegend | 406617 |
| Goat anti-Mouse IgG, HRP (1:10000) | Abcam | ab6789 |
| Goat anti-Rabbit IgG, HRP (1:10000) | Abcam | ab6721 |
| HRP-Conjugated Streptavidin (1:10000) | GeneTex | GTX27403 |
| **Cell Lines** | | |
| MDA-MB-231 | ATCC | HTB-26 |
| HCC44 | DSMZ | ACC 534 |
| MIA PaCa2 | ATCC | CRL-1420 |
| HEK293T | ATCC | CRL-3216 |
| Lenti-X™ 293T cells | Takara Bio | 632180 |
| H460 | ATCC | HTB-177 |
| H1975 | ATCC | CRL-5908 |
| **Critical commercial assays** | | |
| Trans-Blot Turbo RTA transfer Kit, LF, PVDF | BioRad | 1704275 |
| CellTiter-Glo Luminescent Cell Viability Assay | Promega | G7570 |
| Active Ras Pull-Down and Detection Kit | Thermo Fisher Scientific | 16117 |
| DC Protein Assay | Bio-Rad | 5000112 |
| Fe-NTA Phosphopeptide Enrichment Kit | Thermo Fisher Scientific | A32992 |
| Pierce™ High pH Reversed-Phase Peptide Fractionation Kit | Thermo Fisher Scientific | 84868 |
| **Reagents** | | |
| Biotin-PEG3-Azide | Click Chemistry Tools | AZ104 |
| Rhodamine-azide | Synthesized in lab | - |
| Pierce Streptavidin Agarose | Thermo Fisher Scientific | 20353 |
| Pierce High pH Reversed-Phase Fractionation Kit | Thermo Fisher Scientific | 84868 |
| Lipofectamine 3000 Transfection Reagent | Invitrogen | L3000075 |
| Sequencing Grade Modified Trypsin | Promega | V5111 |
| LysC | NEB | P8109S |
| TMT10plex™ Isobaric Label Reagent Set | Thermo Fisher Scientific | 90406 |
| TMTpro™ 16plex Label Reagent Set | Thermo Fisher Scientific | A44520 |
| Halt Protease Inhibitor | Thermo Fisher Scientific | 78438 |
| Halt Protease and Phosphatase Inhibitor | Thermo Fisher Scientific | 78841 |
| DTT | Sigma | D0632 |
| Tris(2-carboxyethyl)phosphine | Research Products International | T26500-10 |

| | | |
|---|---|---|
| Dulbecco's phosphate-buffered saline | Corning | 21-031-CV |
| Titansphere Phos-TiO Bulk Media | GL Sciences | 5010-21315 |
| **Recombinant DNA** | | |
| HA-tagged KRAS | Addgene | 75282 |
| pMAX-GFP | Addgene | 177825 |
| pRK5-FLAG-p14 | Addgene | 42330 |
| **Software and Algorithms** | | |
| Proteome Discoverer (v3.0) | Thermo Scientific | - |
| GraphPad (v9 and v10) | Dotmatics | - |
| Xcalibur software (v4.1.50). | Thermo Scientific | - |
| ImageLab (v6.1.0) | Bio-Rad Laboratories | - |
| ImageJ (v1.53r17) | NIH | - |
| ZEN (v2.3) | Zeiss Microscopy | - |
| Imaris (v10.2) | Oxford Instruments | - |
| Office 365 | Microsoft | - |
| ChimeraX (v1.8) | UCSF | - |
| Python (v3.8.5) | Python Software Foundation | - |
| RStudio (v2024.12.1) | Posit | - |
| Illustrator (v27.8.1) | Adobe | - |

**Supplementary Note**

Synthesis of **3-(3-(but-3-yn-1-yl)-3H-diazirin-3-yl)-N-methylpropanamide** (**P1**), **1-(2- benzylpiperidin-1-yl)-3-(3-(but-3-yn-1-yl)-3H-diazirin-3-yl)propan-1-one** (**P2**), **3-(3-(but-3-yn-1-yl)-3H-diazirin-3-yl)-N-(2-oxo-2H-chromen-6-yl)propenamide** (**P3**), **(E)-N-(2-(3-(but-3-yn-1-yl)-3H-diazirin-3-yl)ethyl)-2,3-diphenylacrylamide** (**P4**), (**R)-N-benzyl-3-(3-(but-3-yn-1-yl)-3H-diazirin-3-yl)-N-(1-phenylethyl)propenamide** (*R*)-**P5**, **(S)-N-benzyl-3-(3-(but-3-yn-1-yl)-3H-diazirin-3-yl)-N-(1-phenylethyl)propenamide** (*S*)-**P5, (R)-3-(3-(but-3-yn-1-yl)-3H-diazirin-3-yl)-N-(1,2,3,4-tetrahydronaphthalen-1-yl)propenamide** (*R*)-**P6 , and (S)-3-(3-(but-3-yn-1-yl)-3H-diazirin-3-yl)-N-(1,2,3,4-tetrahydronaphthalen-1-yl)propenamide** (*S*)-**P6** were performed according to previously published literature procedures[1-3].

Cleavable biotin azide tag: light L-Valine tag was synthesized according to previously published literature procedures[3].

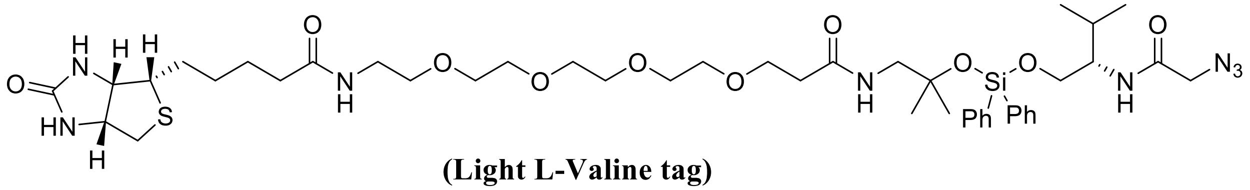

**(Light L-Valine tag)**
